# Functional characterization of *Ixodes* neuropeptide receptors

**DOI:** 10.1101/2025.06.30.662206

**Authors:** Cory A. Knox, Lea L. Kyle, Laura M. Szczesniak, Sarah E. Reks, Richard J. Wojcikiewicz, Barry E. Knox

## Abstract

Neuropeptidergic systems control feeding behaviors in animals, including arthropods. Given the wide variety of pathogens transmitted by ticks during hematophagy, there is an urgency to understand the neural mechanisms responsible for tick feeding behavior. We characterized three *Ixodes* signaling systems that are involved in feeding regulation in other arthropods: neuropeptide CCHamide (CCHa), short neuropeptide F (sNPF) and sulfakinin. RNAs encoding the preproneuropeptides and their receptors were characterized and cDNAs for the receptors were expressed in HEK293T cells by transient transfection. Activation of the receptors by synthetic peptides was monitored by a calcium release (FLIPR) fluorescence assay. There was a single receptor (CCHaR) activated by CCHa (NH_2_-SCKMYGHSCLGGH-amide) containing a disulfide bond with an EC_50_=12 pM, while a scrambled cyclic peptide was inactive at 1 μM. Of the two *Ixodes* NPY-like receptors, NPYLR1A was activated by sNPF (NH_2_-GGRSPSLRLRF-amide) with an EC_50_=1.9 nM. NPYLR1B did not respond to 10 μM sNPF. A single sulfakinin receptor was activated by a sulfated sulfakinin (NH_2_-SDDY(SO_3_H)GHMRF-amide) with an EC_50_=220 pM but not by 1 μM non-sulfated sulfakinin. The *Ixodes* GPCRs were able to couple to endogenous HEK293T G-protein(s). Surprisingly, human but not *Ixodes* GNAQ restored CCHaR responsiveness in HEK293T cells with GNAQ/GNA12 disruptions. Quantitative RT-PCR analysis indicated that all three receptors were expressed in the synganglion. CCHaR/CCHa were found at high levels in the midgut from unfed ticks, and CCHaR expression in the midgut was confirmed by RNAScope *in situ* hybridization. These results establish ligand-receptor identities for three central neuropeptide systems in *Ixodes* and set the stage for structure-function and physiological investigations.

## Introduction

Hard ticks transmit a wide variety of pathogens and are the most common disease vectors in the US (Eisen and Eisen, 2018; Eisen et al., 2017). For example, *Ixodes scapularis* (black-legged tick) carries the spirochete *Borrelia burgdorferi* which causes Lyme disease, the most frequently reported vector-born disease in the United States (Eisen, 2018; Eisen and Eisen, 2018; Kugeler et al., 2021; Nelson et al., 2015). Ticks are obligate hematophagous ectoparasites and spread pathogens to their hosts during active blood-feeding (Eisen, 2018; Eisen and Eisen, 2018; Randolf, 2008; Randolph, 1998; Spielman et al., 1985). Ixodids feed three times during their lifespan followed each time by molting (Oliver, 1989), with extended periods between blood meals (Needham and Teel, 1991). Feeding in non-nidicolous ticks is a series of behaviors that include questing, engagement, attachment and engorgement (Waladde and Rice, 1982). Feeding is initiated from either a quiescent or dormant (via diapause) state through the complex and not well-understood interactions of energy stores (Alasmari and Wall, 2020, 2021; Herrmann and Gern, 2012; Randolph et al., 2002) and water balance (Benoit and Denlinger, 2010; Needham and Teel, 1991), temperature (Alasmari and Wall, 2021; Ogden et al., 2004; Randolph et al., 2002; Thomas et al., 2019), and diurnal (Lees and Milne, 1951; Schulze et al., 2001) or seasonal (Belozerov, 2010; Randolph et al., 2002) photoperiods. Due to the prolonged nature of tick feeding, there is a significant gap in understanding of the central neural pathways and molecular signaling mechanisms that control tick feeding neurophysiology. Neuropeptidergic systems (neuropeptides and neuropeptide G-protein coupled receptors, GPCRs) are ubiquitous (Elphick et al., 2018; Jekely, 2013; Mirabeau and Joly, 2013) and play important roles in metazoan feeding (Arora and Anubhuti, 2006; Geary, 2014; Jekely et al., 2018; Nusbaum et al., 2017; Smith et al., 2014; Sobrino Crespo et al., 2014; Wang et al., 2015). In invertebrates, particularly the insects *D. melanogaster* (Bhumika, 2018; Nassel and Zandawala, 2019; Pool and Scott, 2014) and *Ae. aegypti* (Duvall et al., 2019), appetite, feeding and satiety are regulated by multiple neuropeptidergic systems (Caers et al., 2012; Pool and Scott, 2014; Schoofs et al., 2017). Although the networks controlling such behaviors are complex, three classes of neuropeptides are most thoroughly described in insects (*see* Discussion) and thus are attractive as potential feeding modulators in ticks: the short neuropeptide F (sNPF)/sNPF receptor (sNPFR) and neuropeptide CCHamide (CCHa)/receptor (CCHaR) systems which commonly promote food intake and sulfakinin (SK)/receptor (SKR) which commonly reduces food intake.

Previous studies have partially characterized tick neuropeptides and their associated GPCRs neuropeptide systems using peptidomics, bioinformatics or RNA expression (Christie, 2008; Dinglasan et al., 2013; Egekwu et al., 2014; Gulia-Nuss et al., 2016; Hansen et al., 2011; Neupert et al., 2009; Waldman et al., 2022). Here, we describe the *Ixodes* RNAs encoding CCHa, sNPF and SK and their receptors. We characterized their functional activity using heterologous expression systems for the receptors (Caers et al., 2014; Hansen and Brauner-Osborne, 2009). We describe high-affinity activation of CCHa, sNPF and SK receptors by synthetic neuropeptides, through the endogenous Gq signaling pathway of HEK293T cells. We determine the relative expression levels in unfed ticks using RT-PCR and RNAScope.

## Materials and Methods

### Ticks

Unfed adult *Ixodes scapularis* (NR-42510) and *Ixodes pacificus* (NR-44385) were deposited by the Centers for Disease Control and Prevention and obtained through BEI Resources, NIAID, NIH. They were housed at 22°C in humidified incubators until use.

### Phylogeny and sequence analysis

Sequences in Acari were identified using the standard protein BLAST algorithm and both the non-redundant protein sequences (https://blast.ncbi.nlm.nih.gov/) and VectorBase (https://vectorbase.org/). Sequences (see Data Supplements for accession numbers and sequences used) were aligned (Larkin et al., 2007) using ClustalW2 (www.clustal.org/clustal2) or SnapGene (v6.1, www.snapgene.com). Sequence alignments were further refined, and phylogenetic trees and estimated evolutionary distances inferred using MEGA (v. 12, www.megasoftware.net) software (Kumar et al., 2018; Stecher et al., 2020). Alignments were prepared for figures using Illustrator (Adobe, Inc). DNA sequences of the prepropeptide coding regions were determined by direct sequencing of PCR products derived from both *I. scapularis* and *I. pacificus* RNA (primers are listed in Suppl. Table 2). Predicted signal peptide sequences within the prepropeptides were identified using the SignalP-6.0 (Teufel et al., 2022a) webserver (https://services.healthtech.dtu.dk/service.php?SignalP). Logos were created using WEBLogo3.0 (www.weblogo.threeplusone.com). Secondary structure models of the *Ixodes* GPCRs were made using Protter (wlab.ethz.ch/protter/) and modified in Illustrator. For comparison with human GPCRs (Kooistra et al., 2021), snake plots (www.gpcrdb.org) of human receptor orthologs (endothelin receptor type B (EDNRB, PDB accession number 5GLH) for CCHaR, NPY1 receptor (NPY1R, PDB accession number 7VGX) for sNPF receptor and cholecystokinin receptor type A (CCKAR, PDB accession number 7F8Y) for sulfakinin receptor) were modified in Illustrator. The positions in the transmembrane helices are numbered using the Ballesteros-Weinstein scheme (Isberg et al., 2015). Sequences for the *Ixodes* G-protein alpha subunits were collected from NCBI and compared with human orthologs using MEGA software. A snake plot of human GNAQ was downloaded from GPCR database and modified in Illustrator.

### Expression plasmids

The *Ixodes scapularis* expression cDNAs and the associated amino acid sequences are shown in Supplemental Data (Suppl. Fig. 1). A cDNA encoding *Ixodes scapularis* CCHamide receptor (accession number XM_002415437) was assembled from fragments (nucleotides 1-1053) obtained by RT-PCR from whole adult tick RNA and synthetic primers containing the C-terminus (nucleotides 1054-1107) and the 1D4 epitope (TETSQVAPA, nucleotides 1108-1137). A cDNA encoding Ixodes scapularis sulfakinin receptor (accession number EEC11794) was synthesized following codon optimization for expression in human cells and addition of the ID4 epitope at the C-terminus by gBlocks gene fragments (IDT, Inc. www.idtdna.com). Both receptor cDNAs were directionally cloned in pcDNA3.1. cDNAs encoding two putative sNPF/NPF receptors (NPYLR1A, accession number KC439540 and NPYLR1B, accession number KC439541) in the mammalian expression vector pME18S were provided by Dr. L. Vosshall (Rockefeller University) and have been previously described (Liesch et al., 2013). A cDNA encoding human G protein alpha subunit 15 (accession number NM_002068.4) in the expression vector pcDNA3.1 was provided by Dr. D. Logothetis (Northeastern University). A cDNA encoding *Ixodes scapularis* G protein alpha q subunit (accession number XP_029846564.1) was synthesized following codon optimization for expression in human cells by gBlocks gene fragments (IDT). The IsGNAQ cDNA was directionally cloned into pcDNA3.1. The human G protein alpha q subunit in pcDNA3.1 was obtained from the cDNA Resource Center (www.cdna.org). All cDNA sequences were confirmed by Sanger sequencing (Genwiz, Inc, www.genewiz.com) on both strands.

### Cell lines and transfections

HEK293T cells were obtained from ATCC (CRL-3216, www.atcc.org) and low passage number cells were maintained in DMEM/F12 supplemented with 10% fetal bovine serum and penicillin-streptomycin in 5% CO_2_ at 37C. HEK293 cells with CRISPR/Cas9-mediated genome-editing to eliminate expression of both Gq and G11 alpha subunits (dKO) described previously (Schrage et al., 2015) were provided by G. Milligan (University of Glasgow) and A. Babwah (Rutgers University) and grown in DMEM with 10% fetal bovine serum and penicillin-streptomycin in 5% CO_2_ at 37C. The day prior to transfections, cells were split into either 6-well (4.2 x 10^5^ cells/well) or 10 cm (2.8 x 10^6^ cells/well) plates. Expression plasmid DNA (60 µg) and Fugene6 reagent (90 µl) were incubated with serum-free DMEM in a final volume of 248 µl for 5 min at room temperature. Receptor and GNA plasmids (human GNA15, human GNAQ or *Ixodes* GNAQ) were used in equal amounts. When GNA cDNAs were not included, pcDNA3.1 was used instead. The DNA-Fugene transfection mixture was then added to the cells and incubated about 16 h. The transfected cells were trypsinized, distributed into 96-well polylysine-treated plates (4.2 x 10^5^ cells/well) and further incubated for another 24 h.

### FLIPR/calcium release assay

Transfected cells were used to measure neuropeptide-dependent calcium mobilization assay using the FLIPR©/Calcium 6 assay kit (R8190, www.moleculardevices.com) as described previously (Szczesniak et al., 2021). Transfected cells were incubated with Calcium 6 dye in phenol-free DMEM at 37 C for 2 h. Synthetic peptides (Suppl. Table 1) were obtained from Genemed Synthesis (www.genemedsyn.com), stocks prepared by dissolving in H_2_O and then diluted in phenol-free DMEM for the assay. As positive controls, carbachol (Millipore-Sigma Chemical) and ionomycin (1 mM in DMSO, Millipore-Sigma Chemical) were used. Calcium release was measured using a FlexStation®3 Benchtop Multi-Mode Microplate Reader (www.moleculardevices.com) using excitation 485 nm / emission 525 nm. The fluorescence data from each well was normalized to the pre-stimulus baseline fluorescence (which was set to 1.0) and then replicate wells (n = 3 –12) from each plate were averaged in Excel (www.microsoft.com). Data analysis, non-linear fits to estimate K_d_ values and figures were prepared using GraphPad Prism version 10.5 for Mac (www.graphpad.com).

### qRT-PCR

To measure the variation of receptor and neuropeptide RNA expression levels, whole unfed adult *Ixodes scapularis* or dissected tissues (synganglion, midgut, salivary gland and ovary), were frozen and then used to extract total RNA using the miRNeasy Mini Kit (Qiagen, www.qiagen.com). Samples were homogenized in QIAzol lysis buffer (Qiagen) using the TissueRuptor (Qiagen), and the standard miRNeasy (Qiagen) protocol was followed. RNA quality and quantity was assessed using the RNA6000 Nano kit on the Agilent 2100 Bioanalyzer. cDNA was then synthesized using the Quantitect Reverse Transcription kit (Qiagen). Primers (Suppl. Table 2) were designed using the RealTime qPCR PrimerQuest Tool (IDT) or NCBI Primer-Blast (www.ncbi.nlm.nih.gov/tools/primer-blast/) and obtained from IDT. Real-time PCR was performed on a real-time cycler (CFX384, www.bio-rad.com) using cDNA (produced from 0-50 ng/μl total RNA), 2.5 μM primer mix and SYBR Green I master mix (lifescience.roche.com). PCR amplification was performed as follows: 95 C for 5 m, then 39 cycles of 95 C for 10 s, 60 C for 10 s and 70 C for 30 s. A product melting curve was obtained by heating the samples from 65-95 C (heating rate of 0.1 C per 1 s) with continuous fluorescence measurement. The crossing point was determined using CFX Maestro (v. 2, www.bio-rad.com). Ribosomal protein S4 (Koci et al., 2013) was used as the reference gene in ΔC_T_ comparisons (Schmittgen and Livak, 2008). Figures and statistical analysis were prepared using GraphPad Prism.

### RNAScope *in situ* hybridization

20ZZ probes for RNAScope *in situ* hybridization (Kersigo et al., 2018) were designed by ACD (acdbio.com) for CCHaR (targeting 89-961 of XM_002415437.2, C1 probe), SKR (targeting 2-1120 of XM_029970835.1, C3 probe), sNPFR/NPYLR1A (targeting 2-974 of KC439540.1, C4 probe) and scrambled CCHaR (targeting 89-961 of XM_002415437.2, C4 probe). RNAscope® Multiplex Fluorescent Reagent Kit v2 and RNAscope® 4-Plex Ancillary Kit for Multiplex Fluorescent Kit v2 with Opal 520 (FP1487001KT), Opal 570 (FP1488001KT), Opal 620, (FP1495001KT) and Opal 690 (FP1497001KT) from Akoya Biosciences were used. Unfed adult *Ixodes scapularis* synganglia and midgut were dissected and fixed in 4% paraformaldehyde in PBS overnight followed by ethanol dehydration and rehydration into PBS with 1% Tween-20 and 1% bovine serum albumin. Tissue was permeabilized using a minimal volume of Protease Plus reagent (ACD), incubated at RT for 5 min and then washed with freshly prepared glycine (2 mg/ml in PBST) being careful not to disturb the fragile tissue. Probe hybridization, fluorophore addition and amplification were performed according to the manufacturer’s instructions. Images were taken on either Perkin-Elmer Ultra VIEW Vox spinning disk confocal microscope or a Leica SP8 confocal microscope and analyzed using FIJI/ImageJ (Schindelin et al., 2012). Figures were prepared in Illustrator.

## Results

### Short neuropeptide F (sNPF) and receptor

The *I. scapularis* gene encoding prepropeptide sNPF has three exons and two large introns, the second of which interrupts the coding region (**Fig. 1a**). The gene encodes a protein of 105 amino acids with a 29 amino acid signal peptide predicted using SignalP-6.0 (Teufel et al., 2022b). There are two potential dibasic endoproteolytic cleavage sites (Hook et al., 2018) that yield a predicted processed sNPF peptide with the sequence GGRSPSLRLRFG. The carboxyl terminal glycine is an amidation site in many neuropeptides (Kumar et al., 2016), including those in *Ixodes* (Neupert et al., 2009). Sequencing of the single RT-PCR product from total RNA from unfed adult *I. scapularis* matched the predicted coding sequence (**Fig. 1b**). Unfed adult *I. pacificus* RNA also gave a single product whose sequence was very similar to *I. scapularis* (8 nucleotide substitutions leading to 8 variants in amino acid sequence over 353 bases, **Fig 1c** and Suppl. Fig. 2). A comparison of amino acid sequences shows high sequence similarity in the prepropeptide coding region amongst ticks, with less similarity with other arthropods outside the neuropeptide coding region. The predicted *Ixodes* peptide conforms to the sNPF consensus (**Fig. 1d**) with 8 out 12 identities, xGxxPxLRLRFG.

**Fig. 1.**
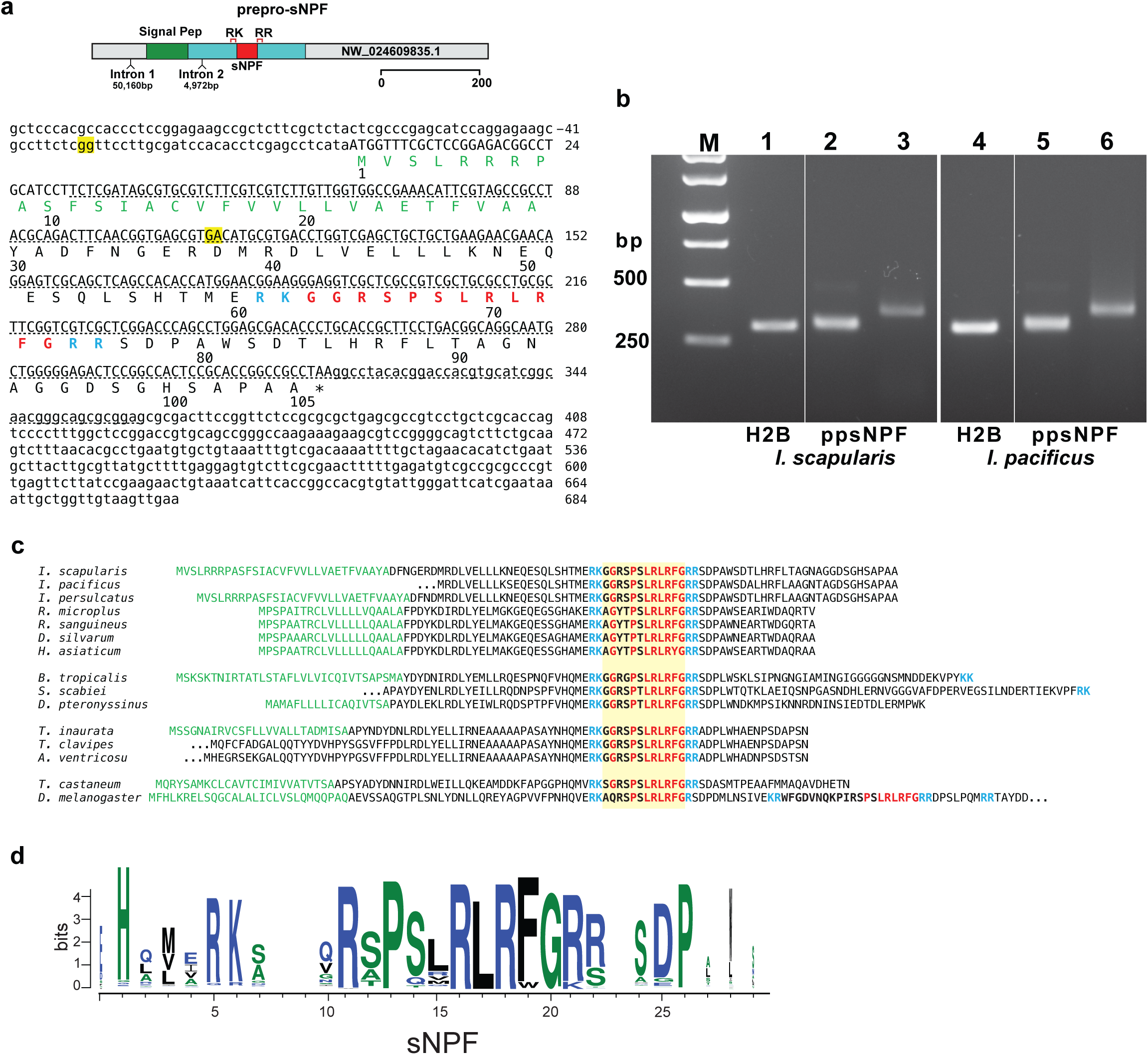
*Ixodes* sNPF. **a**. Organization of the preproneuropeptide sNPF gene showing the positions and sizes of the introns. The sequence of the sNPF mRNA and prepropeptide are shown. The predicted signal peptide is colored green, the potential dibasic endoproteolytic cleavage sites in blue, the predicted sNPF peptide in red. The splice sites are highlighted in yellow. The underlined region was confirmed by sequencing of *I. scapularis* and *I. pacificus* RT-PCR products (b). **b**. RT-PCR analysis of total RNA from unfed *I. scapularis* (lanes 1-3) and *I. pacificus* (lanes 4-6) using different primer pairs (F1-R1, lanes 2, 5 and F2-R2, lanes 3 and 6). Histone H2B primers were used as a control (lanes 1,4). **c**. Sequence alignments of the prepropeptide encoding regions of prepro-sNPF from selected arthropods. The predicted signal peptides are in green and conserved amino acid residues highlighted in red, the potential proteolytic sites are in blue, and the processed peptide is boxed in yellow. **d**. A logo representation of the amino acid sequence alignment of the processed peptide using 232 sequences from arthropods.

Two neuropeptide Y (NPY)-like receptors (NPYLR1A and NPYLR1B) from *I. scapularis* were previously identified by bioinformatics (Liesch et al., 2013). These two receptors are ∼60% identical over the seven transmembrane domain (excluding the divergent N– and C-termini, **Fig. 2a-c** and Suppl. Fig. 3). Both receptor sequences occur in *I. pacificus* and *I. persulcatus*, with a high degree of identity within each subgroup (Suppl. Fig. 4). Two receptors also are found in *Rhipicephalus* species, with a lower sequence identity ∼55%-70% to their *Ixodes* counterparts (**Fig. 2a,b)**. There is one receptor which clusters in the NPYLR1A group in the draft *Dermacentor* genome. Both *Ixodes* receptors are ∼60% identical to the *Drosophila* sNPFR receptor (**Fig. 2d)** and are most related to the vertebrate NPY/ /PrRP GPCRs with whom they share ∼35-40% identity in the transmembrane regions (Suppl Fig. 5). Of the 26 residues in NPY1R or PrRPR that interact with the carboxyl terminal RFamide portion of their cognate peptides ((Kang et al., 2023; Langley et al., 2022; Li et al., 2024; Park et al., 2022; Shen et al., 2024; Tang et al., 2021; Tang et al., 2022), nine (50%) are conserved in both *Ixodes* receptors (Suppl. Fig. 4).

**Fig. 2.**
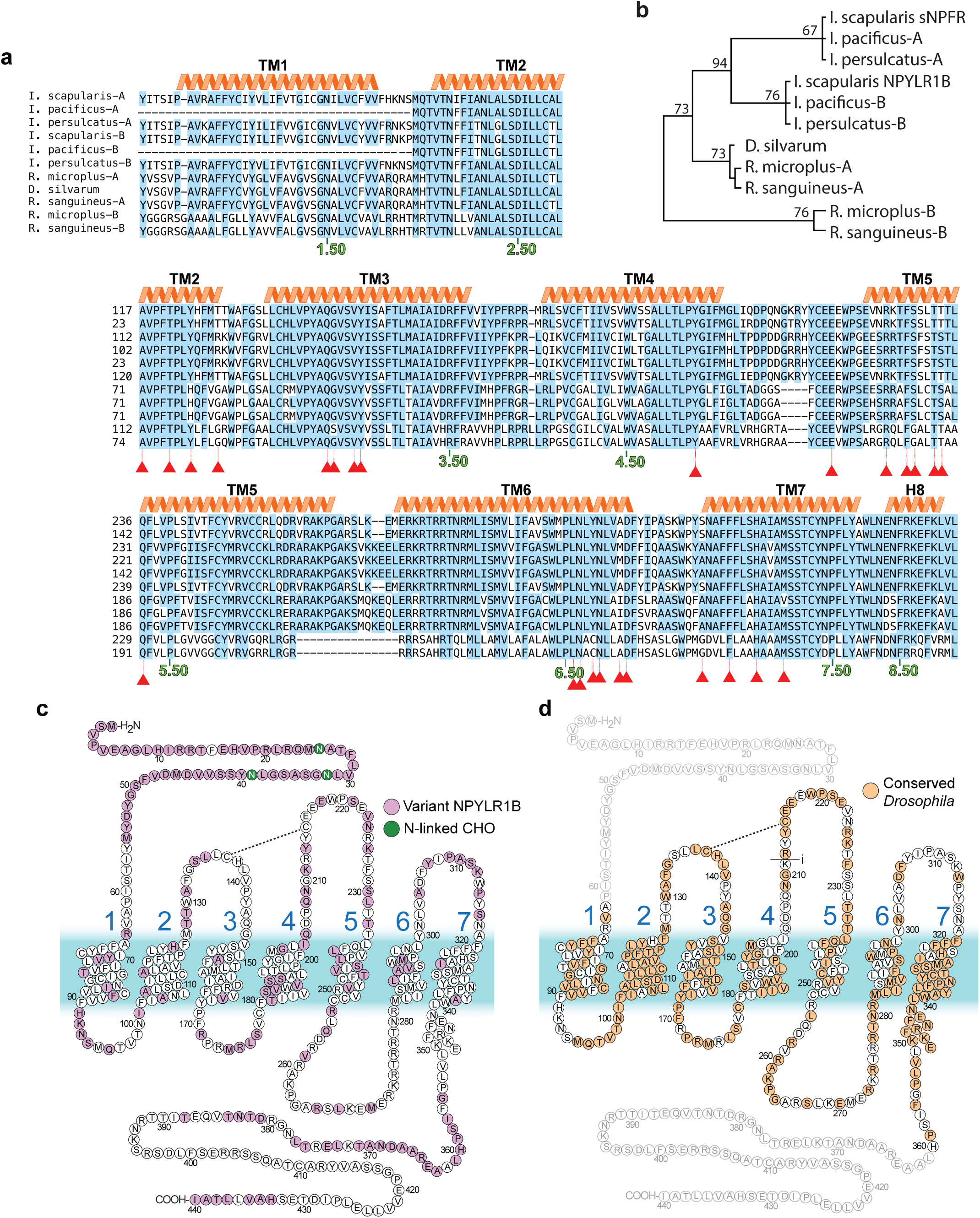
*Ixodes* sNPFR. **a**. Comparison of Ixodid sNPFR amino acid sequences. The two sequences from *Ixodes scapularis* (Dinglasan et al., 2013), originally termed NPYLR1A and NPYLR1B, are labeled A and B respectively. Conserved amino acids (>50%) are highlighted in blue. The positions in the transmembrane helices are numbered using the Ballesteros-Weinstein scheme (Isberg et al., 2015). Positions that interact with NPY in mammalian receptors are indicated with red triangles. **b**. A phylogenetic tree with the highest log likelihood (–3,676.10) inferred using the maximum likelihood method and JTT model (Mega12). The analytical procedure encompassed 11 amino acid sequences with 452 positions in the final dataset. **c**. A model of the secondary structure of NPYLR1A showing the amino acids that are different in NPYLR1B are shown in purple. The N-terminal N-linked glycosylation acceptor sites are shown in green and a disulfide bond between C137-C215 is indicated. **d.** Secondary structure model of sNPFR showing the amino acid positions conserved between *Ixodes scapularis* (NPYLR1A) and *Drosophila melanogaster* sNPFR in orange. The N– and C-termini (*greyed-out*) are not well-conserved and were not included in the comparison.

The NYPL1A receptor responded to Head Peptide-I derived from mosquito heads, an RFamide peptide having the sequence (pyroGlu)RP(hydroxyPro)SLKTRFamide, with an EC_50_ of 427 nM while NPYLR1B had a much higher EC50 of 10 μM (Liesch et al., 2013). To determine whether either *Ixodes* receptor was activated by *Ixodes* sNPF, we employed a 293T cell-based calcium signaling assay (Caers et al., 2014). To facilitate coupling of the *Ixodes* receptors to endogenous calcium-release, a promiscuous G protein alpha subunit (human GNA15) was co-transfected with the receptors (Caers et al., 2014). *Ixodes* NYPL1A responded to the sNPF peptide but did not respond to with a scrambled sNPF sequence (Suppl. Table 1) at concentrations up to 0.1 μM peptide (**Fig. 3a)**. By contrast, *Ixodes* NPYLR1B did not respond to sNPF up to 0.675 μM (**Fig. 3b**). The NPYLR1A receptor had an EC_50_ = 1.9 nM [1.7, 2.1] (mean [95% Confidence Interval]) with respect to sNPF (**Fig. 3c**). These results identify NPYL1A receptor as the *Ixodes* sNPFR receptor. The ligand for NPYLR1B, which is highly conserved in hard ticks, remains unknown. A comparison of amino acids predicted to be in the ligand binding pocket for sNPFR and NPYLR1B suggests potential steric differences that might influence binding of sNPF (Suppl. Fig. 5).

**Fig. 3.**
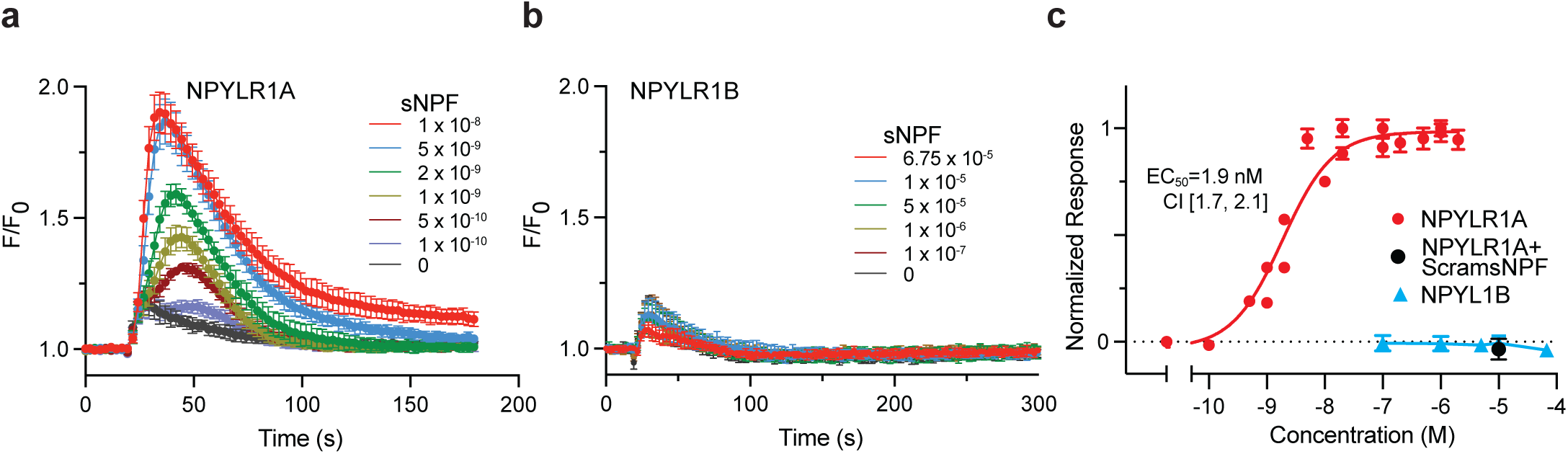
Functional analysis of *Ixodes* sNPFR and NPYLR1B in response to sNPF in transfected 293T cells. **a.** Normalized Ca6 fluorescence responses of cells transfected with NPYLR1A and treated at t = 20 s with synthetic sNPF at the indicated molar concentrations. The symbols are averages of three wells each containing 0.5-1 x 10^5^ cells and error bars are standard deviations. The injection of buffer alone (0) caused a small increase in fluorescence. For reference, maximum responses (*not shown*) elicited by carbachol (375 μM) ranged from 1.8 – 2.0. **b.** Normalized fluorescence responses of cells transfected with NPYLR1B and treated at t = 20 s with synthetic sNPF at the indicated molar concentrations. **c.** Concentration-response relationships for cells transfected with NPYLR1A (*red* or *black*) or NPYLR1B (*blue*). Averaged responses from individual experiments (NPYLR1A, n = 12; NPYLR1A+ScramsNPF, n = 6; NPYLR1B, n = 6) were normalized to the maximum response for that experiment and plotted. Each symbol represents the mean ± SEM. The line is a non-linear fit to the Hill equation with coefficient equal to 1. The EC_50_ and 95% confidence interval is shown for sNPFR responses.

### Sulfakinin neuropeptidergic system

Previous *I. scapularis* synganglion proteomic data identified two peptides in the sulfakinin (SK) family, NH_2_-QDDDYGHMRFa and NH_2_-SDDYGHMRFa (Neupert et al., 2009). A cDNA identified by RT-PCR from *I. scapularis* predicts (**Fig. 4a**) a prepropeptide SK of 100 amino acids encoding peptides flanked by two potential dibasic endoproteolytic cleavage sites (Hook et al., 2018). The carboxyl terminal cleavage site is immediately followed by the termination codon. The two SK peptides are separated by a single Arg. Processing in invertebrates typically occurs if there is a basic amino acid residue in position –4, –6 or –8 (Veenstra, 2000). However, in the tick sequences, basic amino acids occur at –3 (Arg) and –5 (His). Both SK peptides have RFG carboxyl terminal motifs, with a carboxyl terminal amidation site (Kumar et al., 2016). There is a 21 amino acid predicted signal peptide (Teufel et al., 2022b). Although the gene for prepro-SK has not yet been annotated for *I. scapularis*, the ∼300 bp cDNA sequence was found in the ISE6 cell line (Miller et al., 2018) and PalLabHiFi (De et al., 2023) *I. scapularis* genome assemblies, as well as in *I. persulcatus* (Suppl. Fig. 6). Since there are no interruptions in the genomic sequence or obvious splice sites compared to the cDNA sequence, we infer that the prepro-SK gene contains the coding region as a single exon. Sequencing of the single RT-PCR product from total RNA from unfed adult *I. scapularis* matched the predicted coding sequence (**Fig. 4b**). Unfed adult *I. pacificus* RNA also gave a single product with one *I. scapularis* primer pair but not the other (**Fig. 4b**). The sequence of the partial prepro-SK RNA was very similar to *I. scapularis* (5 nucleotide substitutions leading to 1 variant in amino acid sequence over 294 bases, **Fig 4c** and Suppl. Fig. 7). Comparison with other Ixodids shows similar prepro-SK peptides (Suppl. Fig. 7), with tandem SK peptides separated by a single arginine and differing in one residue (a conservative change Asp vs Glu) in position 2 of SK1 (**Fig. 4c**). The Ixodid SK sequence logos show both highly conserved (**Fig. 4d**).

**Fig. 4.**
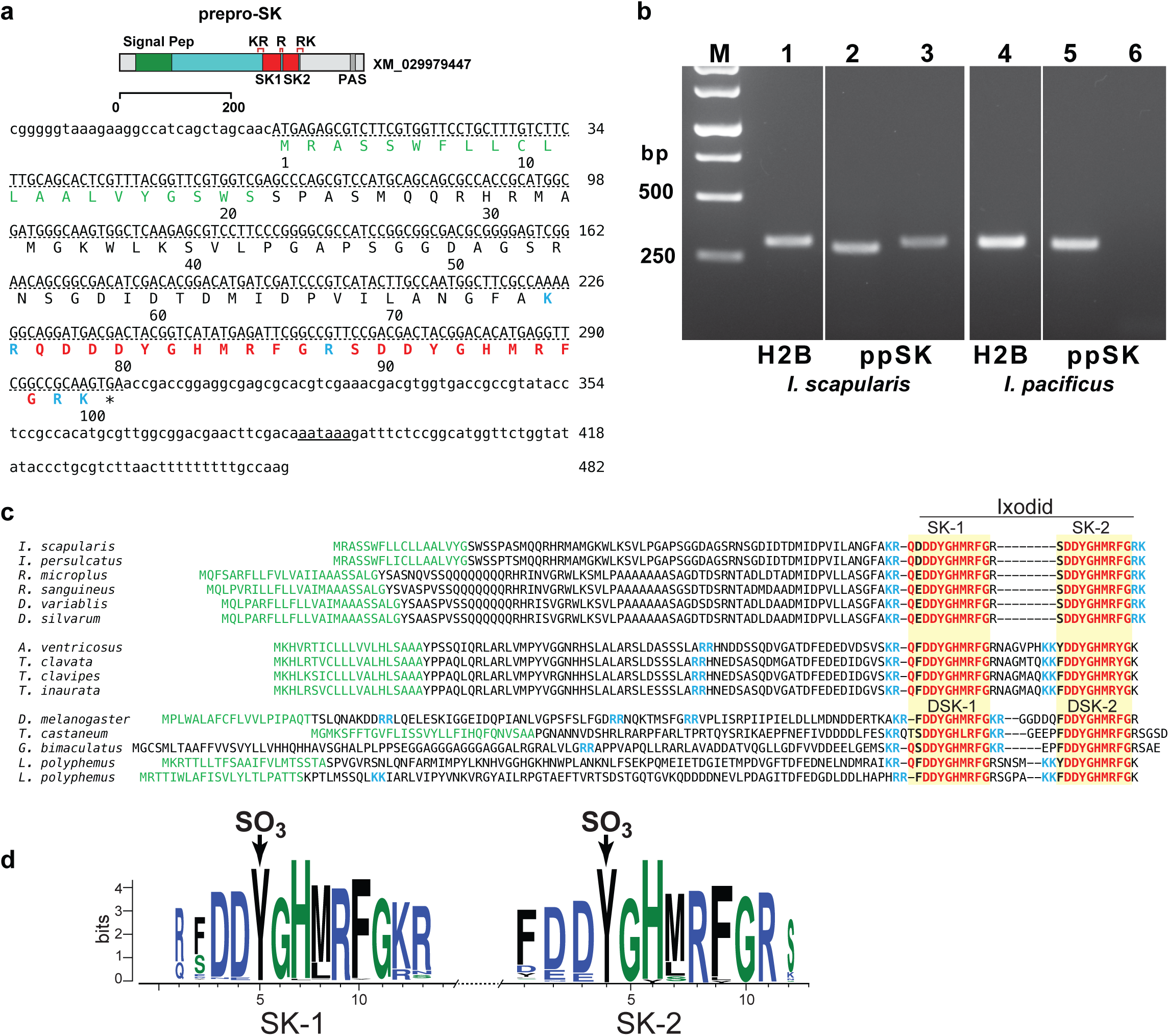
*Ixodes* sulfakinin. **a.** Organization and sequence of the prepro-sulfakinin transcript are shown. The predicted signal peptide is colored green, the potential dibasic endoproteolytic cleavage sites in blue, the predicted processed sulfakinin peptides in red. The underlined region was confirmed by sequencing of the *I. scapularis* and *I. pacificus* RT-PCR products (see b). The sequence of the I. pacificus was only partial. **b.** RT-PCR analysis of total RNA from unfed *I. scapularis* (lanes 1-3) and *I. pacificus* (lanes 4-6) using different primer pairs (F1-R1, lanes 2, 5 and F2-R2, lanes 3 and 6). The second set did not amplify a product in *I. pacificus*. Histone H2B was used as a control (lanes 1 and 4). **c.** Sequence alignments of the prepropeptide encoding regions of prepro-SK from selected arthropods (sequences and accession numbers are provided in the Supplemental Data). The predicted signal peptides are in green (*G. bimaculatus* did not have a predicted signal peptide) and conserved neuropeptide amino acid residues highlighted in red, the potential proteolytic sites in blue, and the final processed peptides are boxed in yellow. DSK-1 and DSK-2 refer to the Drosophila sulfakinin peptides. **d.** A logo representation of the amino acid sequence alignment of the two predicted processed peptides from 272 arthropod sequences. The arrow indicates the sulfation sites.

A single sulfakinin receptor was identified in the *I. scapularis* genome (Cassens et al., 2025; De et al., 2023; Miller et al., 2018) and three other *Ixodes* species (**Fig. 5a**). The *I. hexagonus* and *I. pacificus* sequences were partial, but all of the sequences were >93% identical at the amino acid level (Suppl. Fig. 8) and virtually identical in the seven transmembrane domains (**Fig. 5a**). There are orthologous sulfakinin receptors in other tick species that share >70% amino acid identity with the *I. scapularis* protein sequence, and we term this subgroup SKR2. In addition, both *Rhipicephalus* and *Dermacentor* species have two additional sulfakinin receptors paralogs with a lower amino acid sequence (∼50%) to the *Ixodes* SKR2. There were sulfakinin receptors in selected mite species with >40% amino acid identity, a similar level to the two sulfakinin receptors in *Drosophila melanogaster* (**Fig. 5c**, Suppl. Figs. 9 and 10). We term this subgroup of sulfakinin receptors SKR1. Thus, *Ixodes* species appear to have a single sulfakinin receptor.

**Fig. 5.**
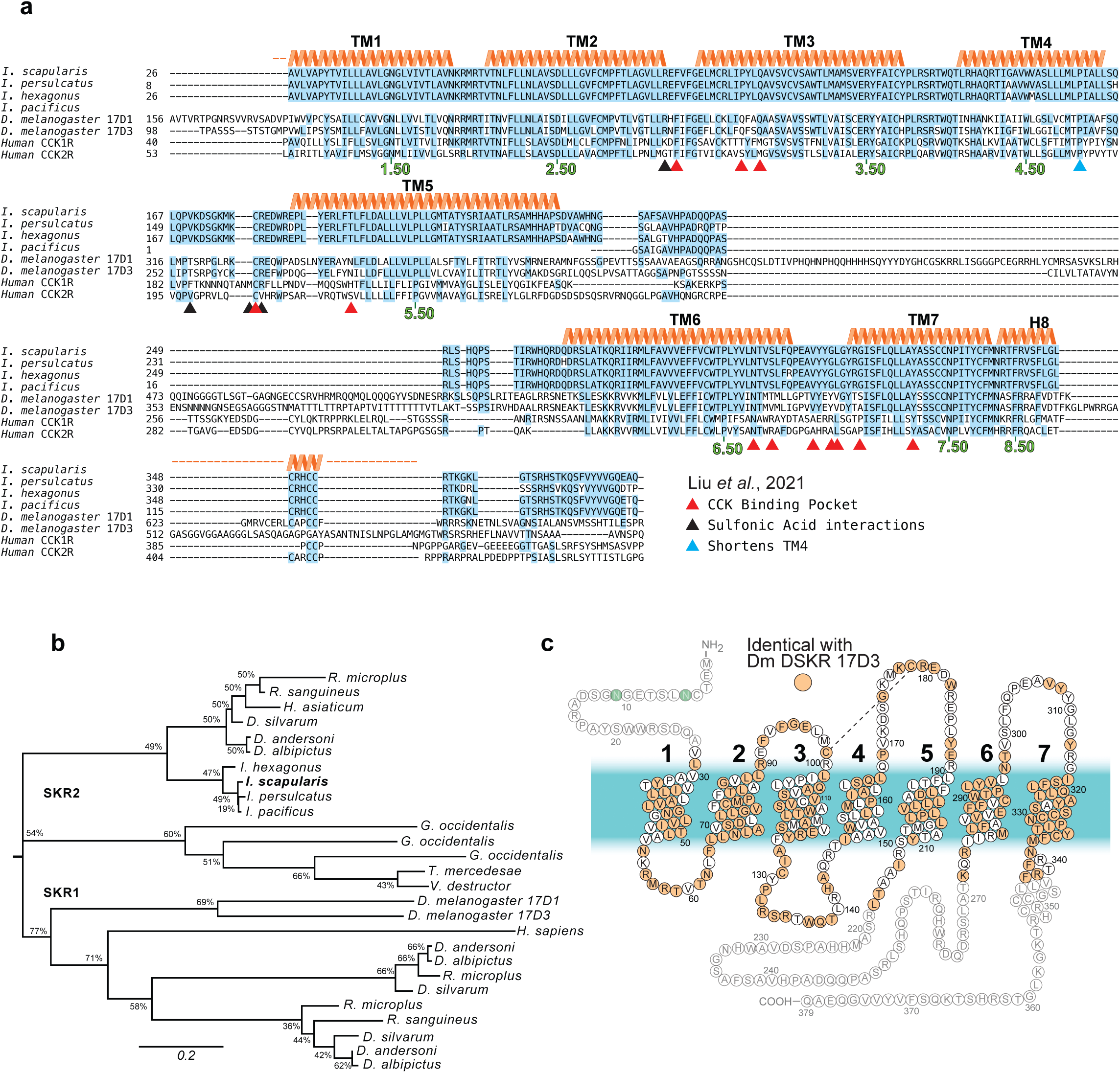
*Ixodes* SKR. **a.** Comparison of sulfakinin receptor amino acid sequences. Amino acids conserved in all sequences are highlighted in blue. The positions in the transmembrane helices are numbered using the Ballesteros-Weinstein scheme. The termini were variable and could not be aligned so they have been omitted. **b.** Phylogenetic tree was inferred using the maximum likelihood method and JTT and the tree with the highest log likelihood (–14,147.37) is shown (MEGA12). The analytical procedure encompassed 27 amino acid sequences with 772 positions in the final dataset. **c.** A model of the secondary structure of SKR showing the amino acids conserved between *Ixodes scapularis* and *Drosophila melanogaster* in orange. The N– and C-termini (*greyed-out*) are not well-conserved and were not included in the comparison.

Both subgroups have high homology in the transmembrane and extracellular loop regions and share significant homology to the human CCK receptors, including P^4.59^ (**Fig. 5a**, blue triangle) which causes an unwinding and thus shortening of TM4 helices in CCKRs compared to other GPCRs (Ding et al., 2022; Mobbs et al., 2021; Zhang et al., 2021). In addition, there is conservation of amino acids that interact with or form a binding pocket for the sulfated tyrosine of their respective neuropeptides, particularly R197^ECL2^ (R90^ECL2^ in *I. scapularis* SKR) which forms an essential hydrogen bond with the sulfonic acid group is found in all of the sulfakinin receptors and other amino acids in the putative sulfakinin binding pocket are also conserved with either CCK1R or CCK2R (**Fig. 5a** and Suppl. Fig. 11).

To determine whether *Ixodes* sulfakinin receptor was activated by *Ixodes* sulfakinin, we employed the 293T cell-based calcium signaling assay. To facilitate coupling of the *Ixodes* receptors to endogenous calcium-release, a promiscuous G protein alpha subunit (human GNA15) was co-transfected with the receptors. *Ixodes* SKR responded (**Fig. 6a and c**) to the sulfated peptide NH2-SDDY(SO_3_)GHMRF-amide with an EC_50_ = 220 pM [191, 260] (mean [95% Confidence Interval]) but did not respond to unsulfated 0.1 μM peptide and very weakly (<10% maximum) 1 μM unsulfated peptide (**Fig. 6b and c)**. These results identify this receptor as the single *Ixodes* sulfakinin receptor with a high affinity to the native sulfated peptide.

**Fig. 6.**
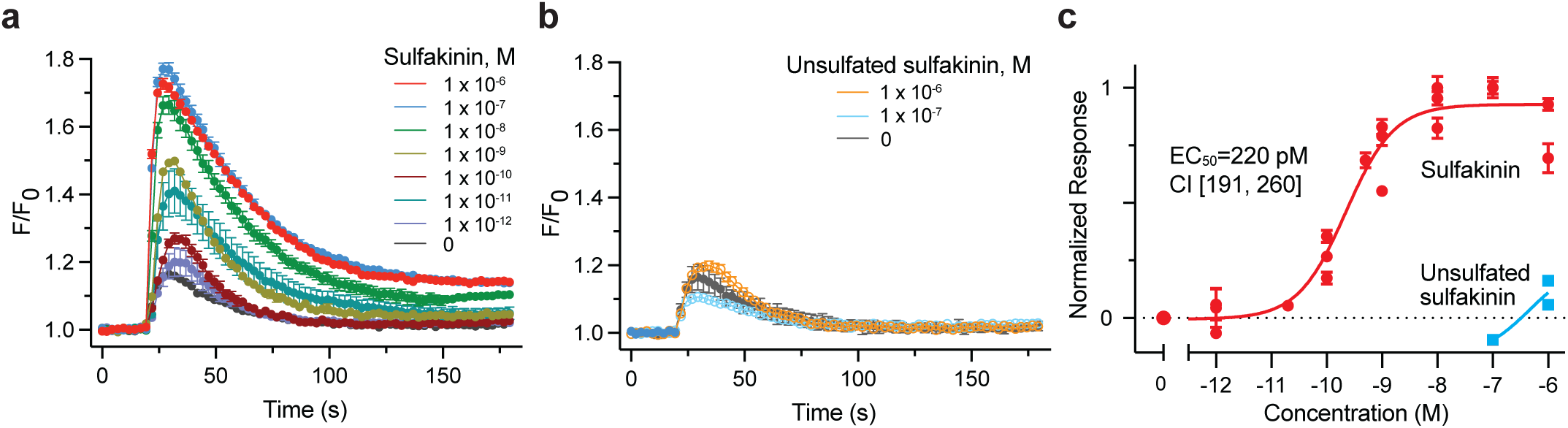
Functional analysis of *Ixodes* SKR in response to sNPF in transfected 293T cells. **a.** Normalized Ca6 fluorescence responses of cells transfected with SKR and treated at t = 20 s with synthetic sulfakinin (containing sulfated tyrosine at position 4) at the indicated molar concentrations. The symbols are averages of three wells each containing 0.5-1 x 10^5^ cells and error bars are standard deviations. The injection of buffer alone (0) caused a small increase in fluorescence. For reference, maximum responses (*not shown*) elicited by carbachol (375 μM) ranged from 1.8 – 2.2. **b.** Normalized fluorescence responses of cells transfected with SKR and treated at t = 20 s with synthetic sulfakinin without sulfated tyrosine at the indicated molar concentrations. The symbols are averages of three wells each containing 0.5-1 x 10^5^ cells and error bars are standard deviations. The responses were not different by paired t-test (p > 0.05). **c.** Concentration-response relationships for cells transfected with SKR. Averaged responses from individual experiments were normalized to the maximum response for that experiment and plotted. Each symbol represents the mean ± SEM (n = 8 – 12). The line is a non-linear fit to the Hill equation with coefficient equal to 1. The EC_50_ and 95% confidence interval is shown for SKR responses elicited by the sulfated SK.

### CCHamide neuropeptidergic system

A genomic region encoding the mRNA encoding prepro-CCHa (Hansen et al., 2011) was identified in the *I. scapularis* genome spanning more than 22 kbp (**Fig. 7a**). RT-PCR using primers at the ends of the predicted prepro-CCHa peptide produced two bands with both *I. scapularis* and *I. pacificus* whole tick RNA (**Fig. 7b**). Sequencing showed that exon 3 occurs in the upper PCR products but not in the lower products in both species. Thus, the CCHa gene produces alternative spliced mRNAs. Both prepro-CCHa peptides (127 and 117 amino acids in length) contain a 24 amino acid predicted signal peptide and multiple potential dibasic endoproteolytic cleavage sites, one of which occurs in the alternatively spliced exon 3. The conserved sequence encoding the CCHa peptide is interrupted by an intron. There are two predicted neuropeptides from the primary cleavage sites: a 24 amino acid peptide NH_2_-EASGGQPSIDSCKMYGHSCLGGHG from the long transcript and a 16-mer NH_2_-NNSCKMYGHSCLGGHG from the shorter transcript. Both peptides are likely amidated on the carboxyl terminal glycine. CCHa peptides were not reported in the previous peptidomics from *I. scapularis* synganglia (Neupert et al., 2009). Predicted prepro-CCHa from other ticks and mites (**Fig. 7c**) have a similar structure as encoded by the *I. scapularis* shorter transcript. Several predicted prepro-CCHa sequences from representative arthropods encode a conserved 14 amino acid CCHa peptide with a signal peptide and a carboxyl terminal dibasic cleavage signal immediately following the carboxyl glycine of the CCHa peptide. Alignment of arthropod CCHa sequences show a highly conserved 14-mer peptide with invariant cysteines residues at positions 2 and 9 **(Fig.7d)**. Sequencing of peptides from *Ixodes* ssp. is required to establish the native CCHa peptide(s).

**Fig. 7.**
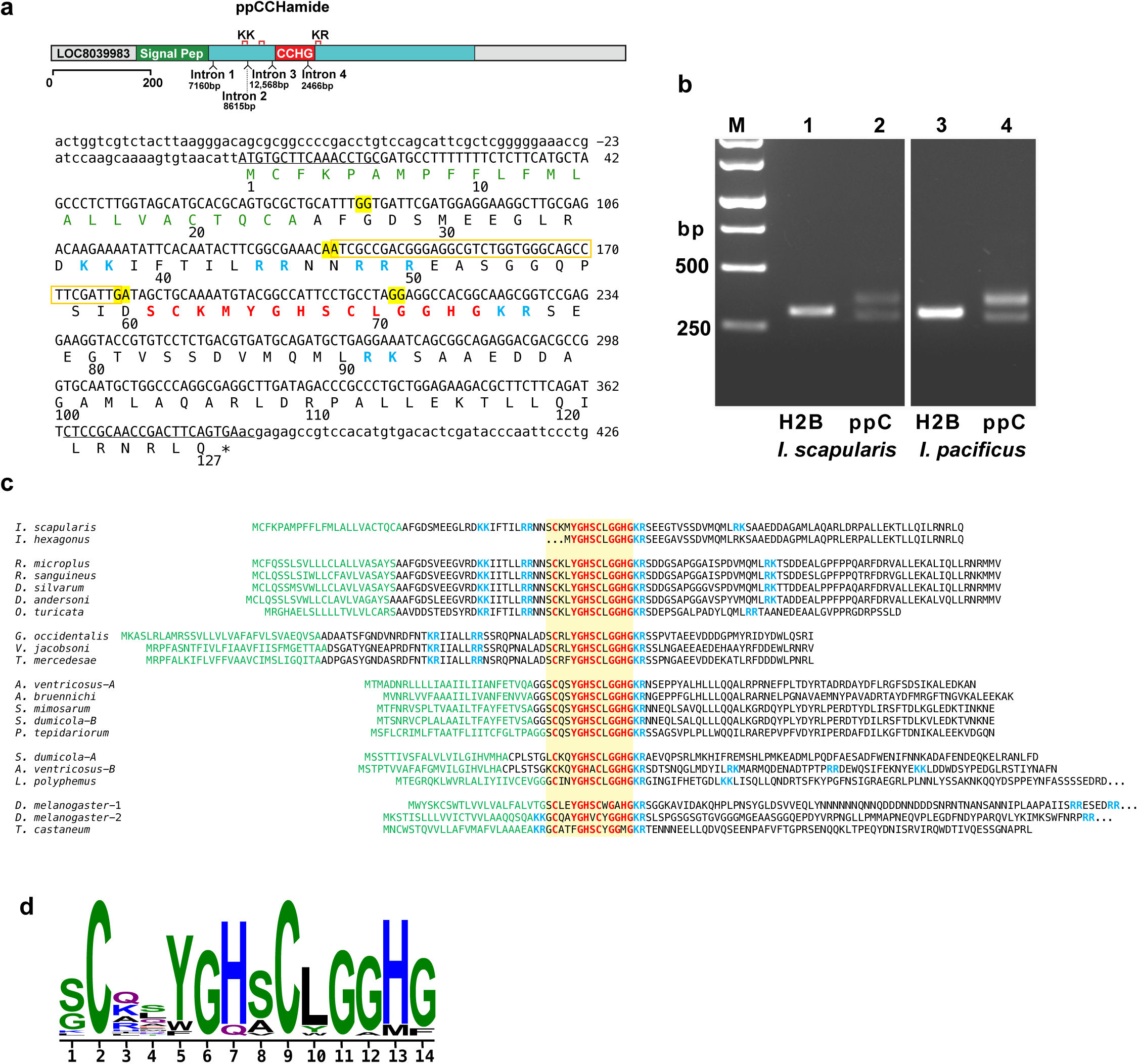
*Ixodes* prepro-CCHa gene. **a.** Organization of the prepro-CCHa gene showing the positions and sizes of the introns. The sequence of the mRNA and prepro-CCHa are shown. The predicted signal peptide is colored *green*, the potential dibasic endoproteolytic cleavage sites in *blue*, the predicted processed CCH peptide in *red*. The splice sites are highlighted in *yellow*. Primers used in RT-PCR are underlined. The alternatively spliced exon 3 is boxed in *yellow*. **b.** RT-PCR analysis of total RNA from unfed *I. scapularis* (lanes 1,2) and *I. pacificus* (lanes 3,4). Histone H2B was used as a control (lanes 1 and 3). **c.** Sequence alignments of the prepro-CCHa. The predicted processed peptides are boxed in yellow and conserved amino acid residues highlighted in red, the potential proteolytic sites are in blue, and the predicted signal sequence is in green. **d.** A logo representation of the amino acid sequence alignment of the processed peptides of 21 arthropod sequences.

A single CCHa receptor (CCHaR) was identified in the *I. scapularis* genome (Cassens et al., 2025; De et al., 2023; Miller et al., 2018) and by RT-PCR with *I. scapularis* and *I. pacificus* RNA (*data not shown*). A further search in the nonredundant protein sequence database returned a single CCHaR sequence in other tick and mite species (**Fig. 8a**). There are orthologous CCHaRs in other arthropod species (**Fig. 8b**), however in most species there are two receptors (**Fig. 8a** shows *D. melanogaster* sequences). The *Ixodes* spp. are >93% identical with each and >84% identical with other tick species and >50% identical to mite species (Suppl. Fig. 13). The *Drosophila* CCHaR sequences are more distantly related (∼40% identical) with the identities clustering in the transmembrane regions (**Fig. 8c**). The CCHaRs belong to family 12 of the rhodopsin β GPCRs (Elphick et al., 2018; Jekely, 2013; Mirabeau and Joly, 2013) with mammalian bombesin and endothelin receptors, which includes the neuromedin B receptor (NMBR/BB1), the gastrin-releasing peptide receptor (GRPR/BB2), an orphan BBN receptor subtype 3 (BRS-3/BB3), and endothelin (ET_A_ and ET_B_) receptors. The *Ixodes* spp. CCHaR are 30-35% identical to these human GPCRs (**Fig. 8a** and Suppl. Fig. 14).

**Fig. 8.**
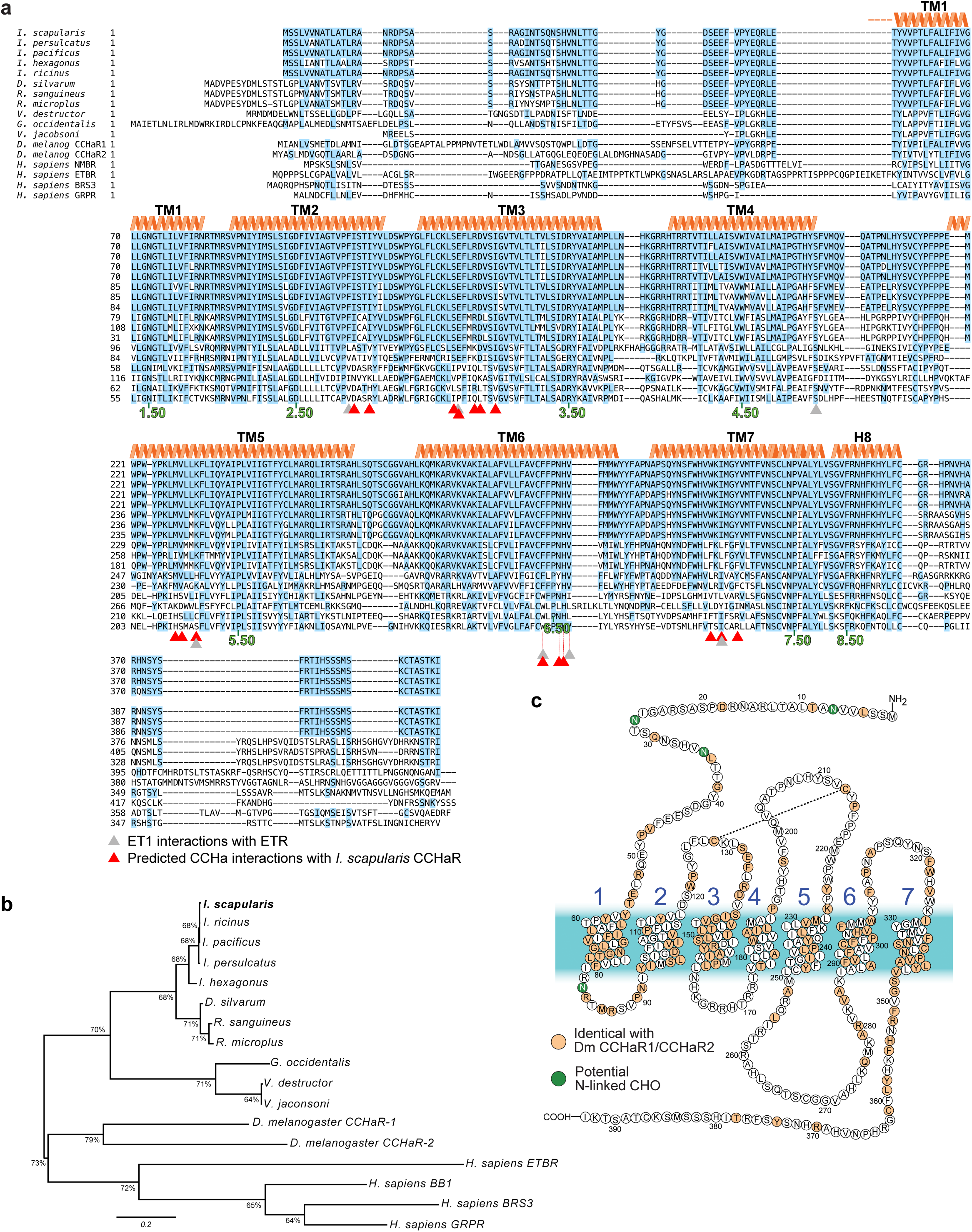
*Ixodes* CCHaR. **a.** Comparison of CCHa receptor (CCHaR) amino acid sequences. Amino acids conserved with *Ixodes* are highlighted in blue. Potential N-linked glycosylation sequences in green. The positions in the transmembrane helices are numbered using the Ballesteros-Weinstein scheme. The termini were variable and could not be aligned so they have been omitted. **b.** Phylogenetic tree was inferred using the maximum likelihood method and JTT and the tree with the highest log likelihood (–8,723.40) is shown (MEGA12). The analytical procedure encompassed 17 amino acid sequences with 578 positions in the final dataset. **c.** Model of the secondary structure of CCHaR showing the amino acids conserved between *Ixodes scapularis* and both *Drosophila melanogaster* receptors in orange.

Although the amino acid sequences between the CCHaR and the mammalian rhodopsin β family 12 GPCR’s neuropeptides are not similar, CCHa and endothelin peptides all have intramolecular disulfide bonds and the endothelins have a twisted structure. Structural modeling was performed on the CCHa neuropeptide, CCHaR and their interaction, using the human ET_B_ receptor structure (Ji et al., 2023; Shihoya et al., 2018; Shihoya et al., 2017) as an initial template. The amino acids predicted to interact with CCHa (**Fig. 8a**, *red arrows* and Suppl. Fig. 15) are identical among the tick and mite CCHaRs, with substantial identity with *Drosophila* receptors. The predicted binding site also shares significant homology with the human ET_B_R (**Fig. 8a**). The structure of the predicted binding pocket fits the cyclic CCHa with the peptide inserting into the receptor core, which is conserved in Ixodids (Suppl. Fig. 15). Positions equivalent to the ET1 binding site in ETHB (grey arrows) and those predicted for CCHaR (red arrows) are shown.

To determine whether *I. scapularis* CCHaR was activated by *I. scapularis* CCHa, we employed a 293T cell-based calcium signaling assay with human promiscuous GNA15 co-transfected with the receptors. *Ixodes* CCHaR responded (**Fig. 9a and b**) to the 13-mer peptide SCKMYGHSCLGGH-NH_2_ containing a disulfide bond between C2-C9 with an EC_50_ = 12 pM [11, 14] (mean and 95% CI). There were no responses to < 0.1 μM peptide with a scrambled CCHa sequence containing a disulfide bond in between different positions (C4-C12) or to a CCHa peptide with two additional positively charged residues (CCHaKR), both followed by terminal amides (**Fig. 9c**). When the peptide concentration was increased, there were responses to 10 μΜ CCHKRamide or 1-10 μM for scrambled CCHa. These results identify this receptor as the single *Ixodes* CCHa receptor that has a high affinity for the native peptide.

**Fig. 9.**
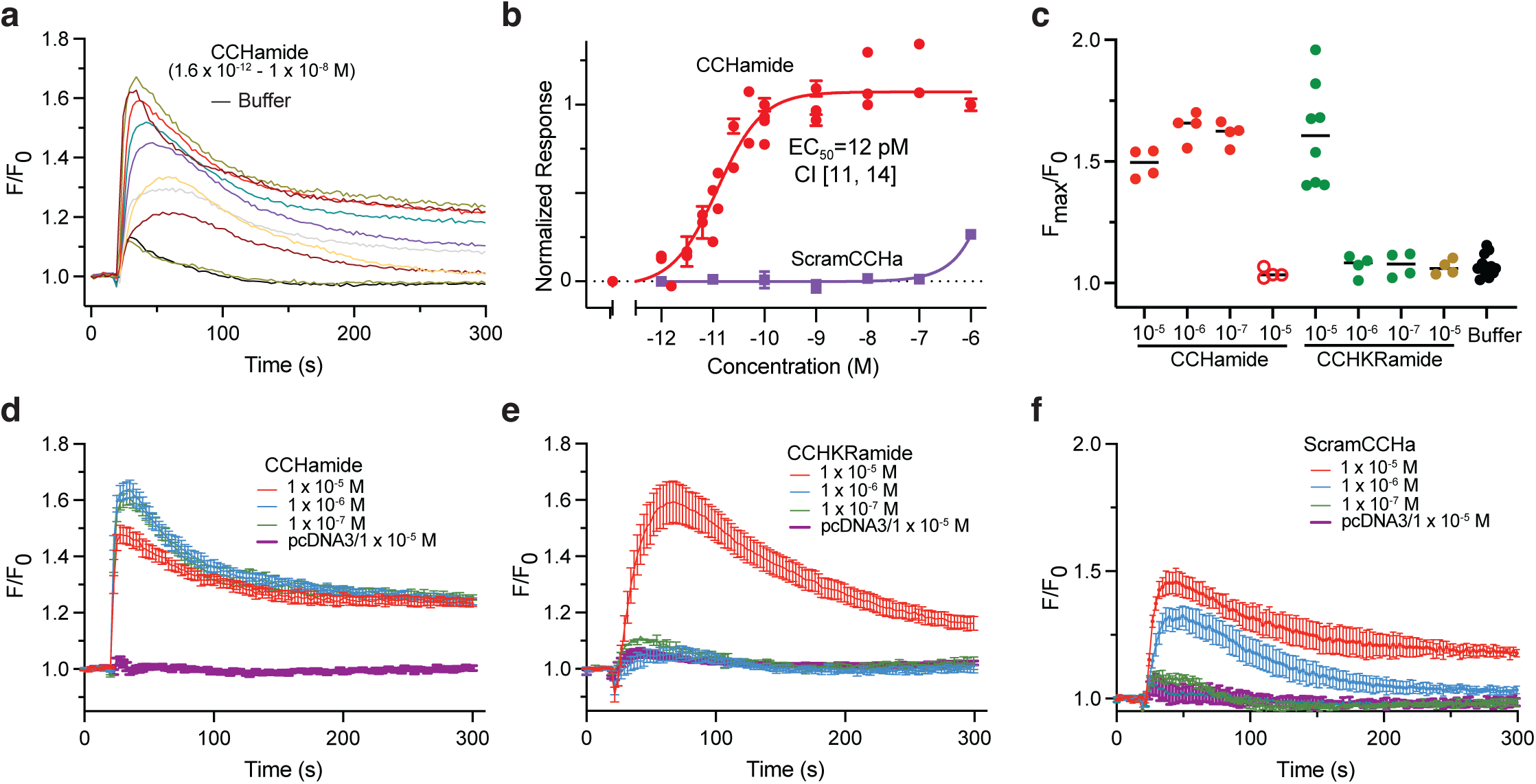
Functional analysis of *Ixodes scapularis* CCHaR in response to CCHamide in transfected 293T cells. a-c. Normalized Ca6 fluorescence responses of cells transfected with CCHaR or empty vector pcDNA3 plus the promiscuous human Gα15 subunit and treated at t = 20 s with the indicated molar concentrations of synthetic CCHa (a) or CCHKRa (b) both containing a disulfide bond between Cys2–Cys9 for CCHa or Cys4–Cys12 for the scrambled CCHamide sequence (c). The symbols are averages of three wells each containing 0.5-1 x 10^5^ cells and error bars are standard deviations (n = 4 – 8 wells). The injection of buffer alone (0) caused a small increase in fluorescence. For reference, maximum responses (*not shown*) elicited by carbachol (375 μM) ranged from 1.8 – 2.0. **d.** Comparisons of the maximum fluorescence changes for the individual wells in (a-c). **e.** Normalized fluorescence responses of individual wells containing cells transfected with CCHaR and treated at t = 20 s with synthetic CCHa with range of in M: 10^-8^, 10^-9^, 10^-10^, 5 x 10^-11^, 2.5 x 10^-11^, 6.25 x 10^-12^, 3.13 x 10^-12^, 1.56 x 10^-12^ and buffer alone. **f.** Concentration-response relationships for cells transfected with CCHaR and GNA15. Averaged responses from individual experiments were normalized to the maximum response for that experiment and plotted. Each symbol represents the mean ± SEM (n = 30). The line is a non-linear fit to the Hill equation with coefficient equal to 1. The EC_50_ and 95% confidence interval is shown for CCHa responses.

**Fig. 10.**
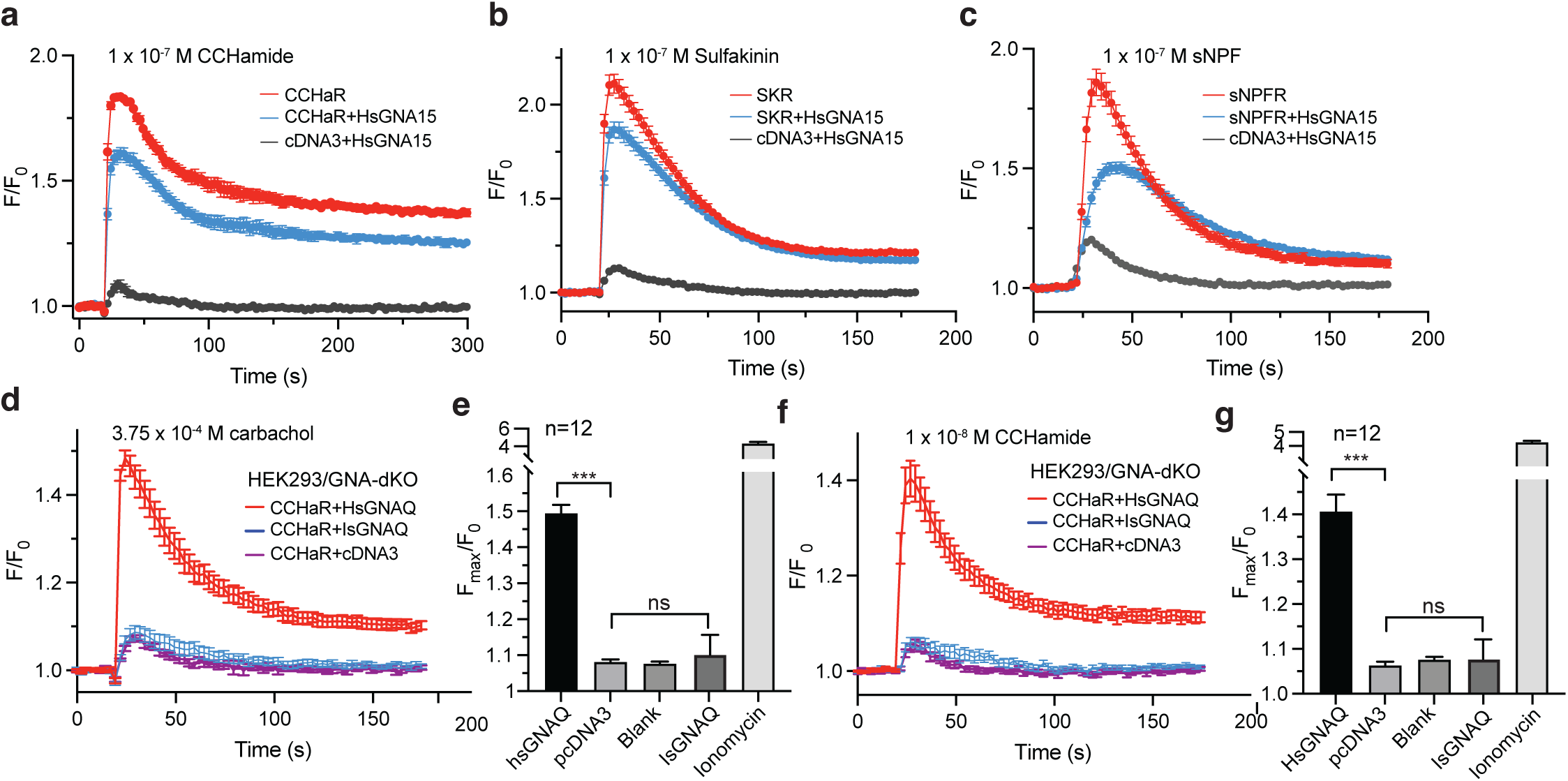
*Ixodes* GPCR coupling to G-proteins in 293T cells. **a.** Normalized Ca6 fluorescence responses (n = 4) from HEK293T cells transfected with CCHaR (*red*), CCHaR plus human GNA15 (*blue*) or empty pcDNA3 plus GNA15 (*black*) treated with 1 x 10^-7^ M CCHamide at t = 20 s. **b.** Normalized Ca6 fluorescence responses (n = 4) from HEK293T cells transfected with SKR (*red*), SKR plus human GNA15 (*blue*) or empty pcDNA3 plus GNA15 (*black*) treated with 1 x 10^-7^ M sulfakinin at t = 20 s. **c.** Normalized Ca6 fluorescence responses (n = 4) from HEK293T cells transfected with sNPFR (*red*), sNPFR plus human GNA15 (*blue*) or empty pcDNA3 plus GNA15 (*black*) treated with 1 x 10^-7^ M sNPF at t = 20 s. The injection of buffer alone (0) caused a small increase in fluorescence. **d-g**. Mutant HEK293T cells (n = 12) with both GNAQ and GNA11 eliminated (GNA-dKO) with transfected with CCHaR and human (HsGNAQ, *red*)), *Ixodes scapularis* (IsGNAQ, *blue*) or pcDNA3 (*purple*). Cells were treated with 375 μM carbachol (d, e) or 1 x 10^-8^ M CCHamide (f, g). Ionomycin (1.25 μM) was used as positive control for cell viability. Each symbol represents the mean ± SEM. e, g. Comparison of the average maximum responses for each group of dKO cells from d and f, respectively. ***, p<0.001; ns, not significant at p<0.05. Error bars are SEM (n = 12).

### *I. scapularis* receptor responses depend upon GNAQ

The responses of *Ixodes* neuropeptide receptors (**Figs. 3, 5, and 9**) were performed in 293T cells co-transfected with human GNA15, which couples to many different GPCRs (Giannone et al., 2010; Offermanns and Simon, 1995). 293T cells express two of the four Gq alpha family subunits that couple to phospholipase C, GNAQ and GNA11 but not GNA14 or GNA15 (Atwood et al., 2011; Schrage et al., 2015). We tested whether the *Ixodes* receptors couple to the endogenous 293T subunits by transfecting cells with only receptors but not with GNA15 (**Fig. 10**). Cells transfected with each *Ixodes* receptor elicited robust responses to its cognate neuropeptide. In fact, the responses were consistently larger than when co-transfected with human GNA15. This may reflect reduced expression of the receptors caused by competition with GNA15, since both cDNAs are driven by the strong CMV promoter.

To determine whether *Ixodes* CCHa receptors can couple to GNAQ, we utilized a cell-line in which both Gq and G11 alpha subunits (dKO-293T) are disrupted (Schrage et al., 2015). Cells transfected with *Ixodes* CCHaR failed to produce responses in dKO-293T cells when stimulated with 100 nM neuropeptide which elicit maximal responses in 293T cells or with carbachol which stimulates endogenous muscarinic receptors (**Fig. 10d**). When human GNAQ was co-transfected with the *Ixodes* neuropeptide receptors in dKO-293T cells, both CCHa– and carbachol-elicited responses were restored (**Fig. 10d-g**). Thus, the *Ixodes* CCHa receptor is able to couple through Gq, similar to its human homologs (Masuho et al., 2023).

We identified the amino acid sequences of the *Ixodes* G protein alpha subunits, which have not been previously described. We found nine complete *Ixodes scapularis* coding regions (**Fig. 11a**): two GNAS, GNAI, GNAI2, GNAO, GNA12, GNA11 and two GNAQ. We found a similar array of GNA sequences in other *Ixodes* species (**Fig. 11b** and Suppl. Fig. 16, although not all of the sequences were complete coding regions). The two predicted GNAQ sequences (X1 and X2) are about 96% identical with the largest variation between positions 309-319. The X1 sequence is about 85% identical to human GNAQ while the X2 form is more divergent (again within the same region as it diverges with X1). To determine whether the *Ixodes* GNAQ could functionally couple to *Ixodes* CCHaR and human Gβγ/PLC, a cDNA with humanized codon usage was synthesized and co-transfected into 293T cells dKO with CCHaR. Despite the high sequence similarity to the human GNAQ **(Fig.11c)**, the *Ixodes* GNAQ did not restore responsiveness to either carbachol (**Fig. 10d, e**) or CCHa in cells transfected with CCHaR (**Fig. 10f, g**).

**Fig. 11.**
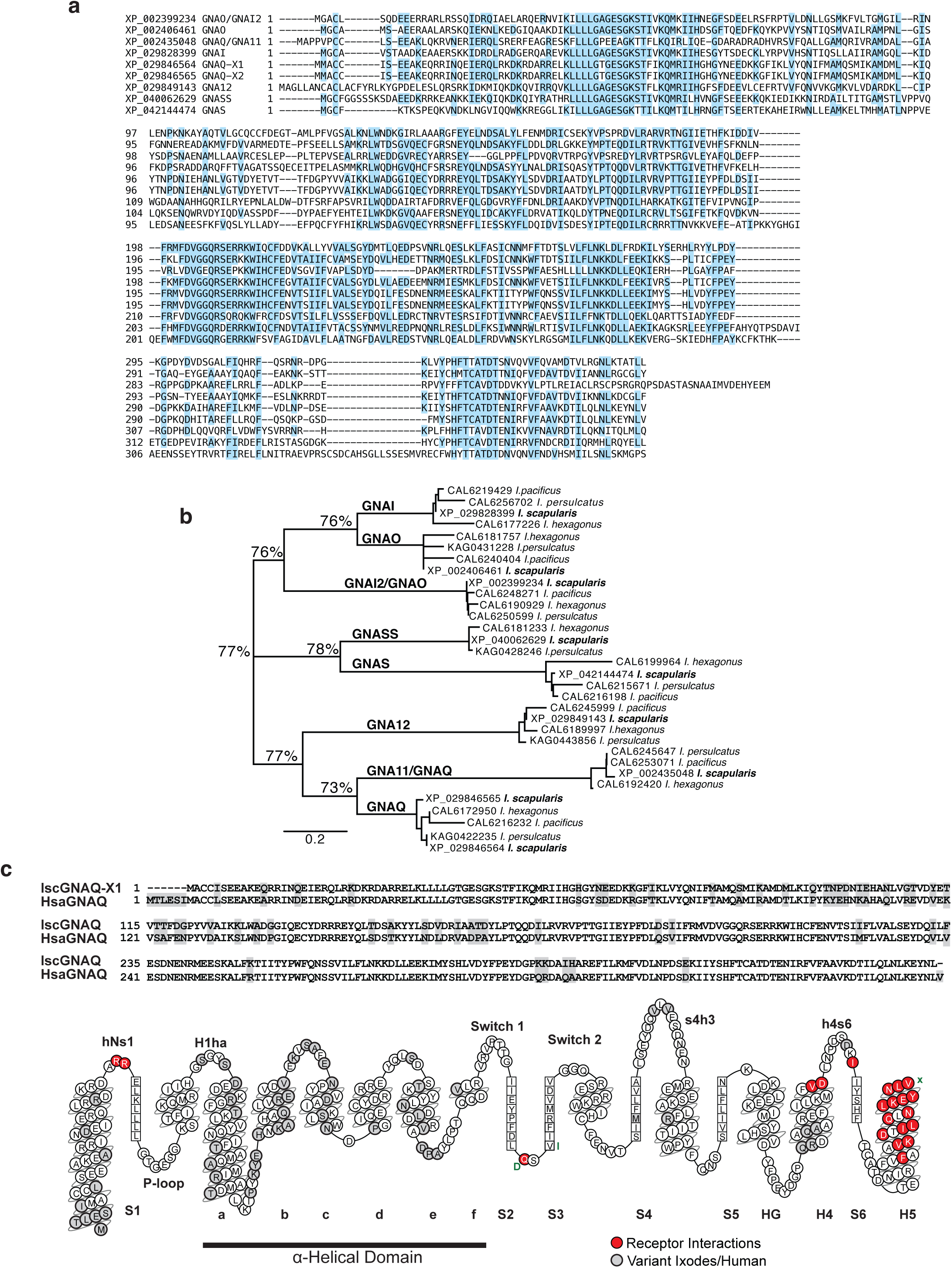
*Ixodes* G-protein alpha subunits. **a.** Comparison of *Ixodes* GNA amino acid sequences. Amino acids conserved in all sequences are highlighted in blue. **b.** Phylogenetic tree was inferred using the maximum likelihood method and JTT and the tree with the highest log likelihood (–8,973.08) is shown (MEGA12). The analytical procedure encompassed 32 amino acid sequences with 467 positions in the final dataset. **c.** Alignment of human and *Ixodes* scapularis GNAQ-X1 with the variant positions highlighted in gray. A snake plot of *Ixodes* GNAQ highlighting the secondary structure and functional regions.

### Expression analysis of *Ixodes* neuropeptidergic systems

The expression patterns of the GPCRs in specific body tissues (synganglion, midgut, salivary gland and ovary) was investigated by qRT-PCR. sNPFR was prominently expressed in the synganglion and to a much lower extent in the salivary gland and almost undetectable in the midgut in both males and females (**Fig. 12a**). SKR was expressed predominantly in the synganglion at a much higher level than sNPFR and very low levels in other tissues (**Fig. 12b**). CCHaR was expressed at higher levels in all tissues, with highest levels in the synganglion (**Fig. 12c**). We examined the expression of genes for the neuropeptides using RNA extracted from whole unfed nymphs (**Fig. 12d**). cDNA derived from both CCHa and SK genes were detected at similar levels, while sNPF cDNA was found at very low levels, the C_T_ values were too high to accurately measure expression suggesting very little transcript was present (**Fig. 12d**). We observed SK expression in synganglion but not in midgut in unfed adults (**Fig. 12e**). CCHa was expressed in the synganglion at very high levels near RPS4 and also at high levels in the midgut in unfed adults (**Fig. 12f**). The expression levels in unfed adults were similar between males and females.

**Fig. 12.**
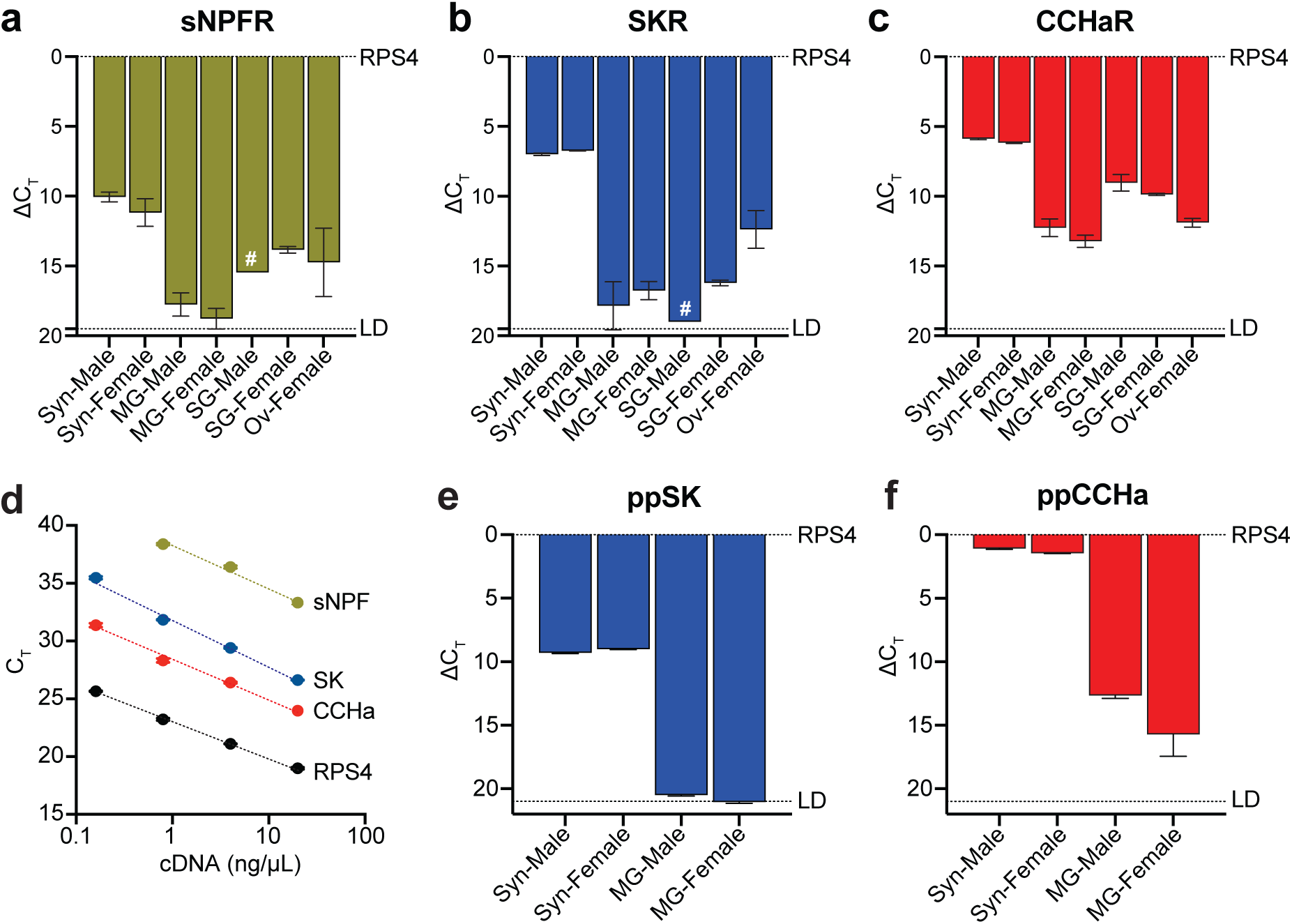
Transcript abundance in various tissues using quantitative real time PCR. Relative transcript abundance of *Ixodes* receptors (**a-c**) and preproneuropeptides (**d-f**) was estimated using qRT-PCR. Threshold cycles of the target RNA were subtracted (ΔC_T_) from the threshold cycles of the ribosomal protein S4 (RPS4) as a reference RNA. In **d**, C_T_ are plotted as a function of input cDNA derived from unfed nymphs. Detection limit (LD) is indicated. Each sample was measured in triplicate and error bars represent standard deviations. The dashed line represents the limit of detection for the samples. RNA from both male (M) and female (F) adult unfed ticks was extracted from synganglia (Syn), midgut (MG), salivary gland (SG) and ovary (Ov). # indicates the C_T_ was from a single sample.

We investigated the expression of GPCRs in unfed adult tissues using whole mount *in situ* hybridization RNAScope. Midgut stained prominently with the CCHaR probe but not with a probe derived from a scrambled version of the CCHaR coding region (**Fig. 13b-f**). No staining was observed for sNPFR or SKR. There was strong staining with CCHaR probes in the anterior segment of the midgut, nearest to the synganglion, and less intense staining in more posterior segments of the midgut. There was also sporadic staining of cells in the synganglion **(Fig. 13g)**. SKR and sNPFR receptor probes also gave signals in the synganglion (**Fig. 13g, h, i**), however challenges with balancing permeability or access of the probes while maintaining tissue integrity during the whole mount *in situ* hybridization prevented consistent staining of the synganglion.

**Fig. 13.**
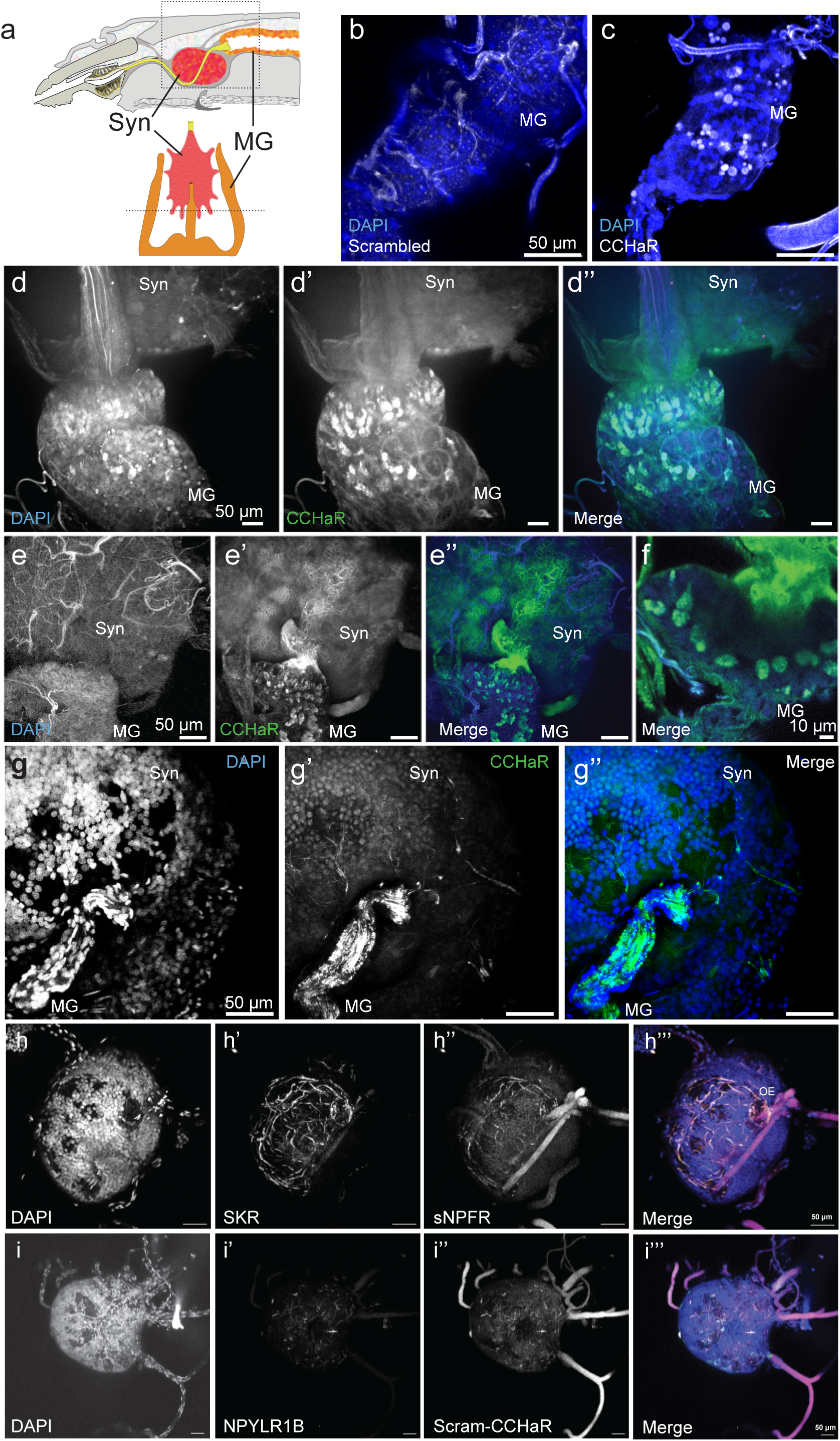
Detection of CCHaR in *Ixodes* midgut and synganglion. **a.** Diagram of the regions dissected for the *in situ* hybridization. Tissues were hybridized with either a set of RNAScope probes to (**b**) a scrambled RNA sequence or (**c-g**) to CCHaR RNA. **h**. A synganglia stained with multiplexed RNAScope probes to SKR (h’) and sNPFR (h’’) and counter-stained with DAPI (h). **i**. A synganglia stained with multiplexed RNAScope probes to NPYLR1B (i’) and a probe designed using a scrambled CCHaR sequence (i’’) and counter-stained with DAPI (i). Merged images are also shown and all tissues were counterstained with DAPI. Synganglia (Syn), esophagus (OE) and midgut (MG) are labeled. Scale bar, 50 μm.

## Discussion

We compared the coding regions of *Ixodes* genes encoding the three receptors and their preproneuropeptides and the *Ixodes* G alpha and beta subunits, with other hard ticks and human homologs. There was high amino acid conservation amongst the tick proteins. Moreover, amino acids that interact with ligands or G-protein alpha subunits were conserved in all three receptor subtypes. We used a calcium release fluorescence assay in transiently transfected HEK293T cells to characterize the activation of three GPCRs by their neuropeptides. We found that all receptors exhibited high affinity interactions with their neuropeptides, having EC_50_ in the pM (CCHaR and SKR) or nM (sNPFR) range.

The human GPCR homologs of the receptors studied here signal through Gq as well as other G protein pathways: i) three ETBR through Gs, Gi Gq; ii) ETAR through Gq; 3) NMBR/GRPR through Gq; iv) BRS3 through Gs, Gq and G12 constitutively; v) NPYR isoforms through Gi (mainly Gi2) and Gq; vi) CCKR1through Gq/11, also Gs and G13; and vii) CCKR2 through Gq. Here, we show that all three *Ixodes* GPCRs were able to couple to human Gq to activate calcium release. A puzzling result was the failure of *Ixodes* GNAQ to restore CCHa responsiveness to GNAQ/GNA12 mutant cell line transfected with CCHaR given the high degree of sequence identity between the human and *Ixodes* GNAQ. It is not clear why the *Ixodes* GNAQ fails to couple CCHaR to the endogenous 293T phospholipase C. The *Ixodes* GNAQ may fail to assemble with human GNB. 293T cells express five GNB genes, most prominently GNB1 and GNB2, and to a lower level GNB5 (Atwood et al., 2011), which contact alpha subunits at multiple interfaces (Oldham and Hamm, 2008). We compared the amino acid sequences of the *Ixodes* G protein beta subunits, which have not been previously described, to the human GNB family (Suppl. Fig. 18). We found two complete coding regions, one that grouped with the human GNB1 and another with GNB5. The *Ixodes* GNB1 was about 83% identical to the human homolog while the *Ixodes* GNB5 was about 71% (Suppl. Fig. 19). A comparison of the variant positions of *Ixodes* GNAQ and GNB1 with the structure of Gq bound to a GPCR (Wu et al., 2024) shows that the variant positions are not in interface regions with receptor or between the G protein subunits (Suppl. Fig. 20). While there are no obvious reasons that suggest that *Ixodes* GNAQ would not assemble with 293T GNB subunits, future work is needed to determine whether they in fact form the G-protein trimer or to identify the amino acid(s) that prevent formation of a hybrid *Ixodes*/human trimer. An alternative explanation for the lack of functional rescue by *Ixodes* GNAQ could be a failure to interact with human PLCβ. We searched for *Ixodes* PLCβ coding regions, which have been previously described We found one complete sequence, related to the human PLCβ family, which is only 46% identical (Suppl. Fig. 21). Moreover, a mapping of the *Ixodes* GNAQ and PLCβ sequences onto the mammalian GNAQ-PLCβ complex (Lyon et al., 2013) shows many amino acid variants in the interface between GNAQ and PLCβ (Suppl. Fig. 21). Thus, it is possible that *Ixodes* GNAQ may not interact effectively with human PLCβ3. However, the possibility of unique interaction interfaces for tick G-protein-effector complexes offers a potential avenue to explore for developing specific agents to disrupt tick signaling pathways without interfering with mammalian counterparts. Future work will be needed to clarify the signaling pathway discrepancies.

We found using quantitative RT-PCR analysis and RNAScope *in situ* hybridization that CCHa and CCHaR were found at high levels in the midgut, indicating that CCHa levels are high in unfed adults and may play a role in appetite or food seeking in ticks. However, further work is needed to characterize whether or how these neuropeptides change in response to feeding. The CCHa (Nagata, 2016b) receptors, and the related excitatory peptide (Morishita and Minakata, 2016), are orthologs of vertebrate neuromedin-B/bombesin/endothelin receptors (Elphick et al., 2018). The arthropod peptides are named for a conserved pair of cysteine residues that form a disulfide bond and an amidated C-terminal His residue. In *Drosophila* (Hansen et al., 2011) and other insects, there are two CCHa and receptor genes (Elphick et al., 2018). dmCCHa-1 modulates sensory responses and olfactory behavior (Farhan et al., 2013). dmCCHa-2 has a role in feeding, however, there are conflicting data on its mechanism of action (Hansen et al., 2011; Li et al., 2013; Lin et al., 2019; Ren et al., 2015; Sano et al., 2015). In one study, disruption of the peptide *ccha-2* reduced food intake by larval and adult flies, with an accompanying reduction in locomotion (Ren et al., 2015). In another study, no differences in larval feeding with dmCCHa-2 receptor disruption were observed (Sano et al., 2015). dmCCHa-2 mRNA levels were sensitive to nutrition state and also the type of nutrients in the food (Millington et al., 2021), decreasing during starvation and recovering after re-feeding (Sano et al., 2015). Moreover, *ccha2* mutants showed significant alterations of insulin-like peptide (*Dilp*) levels in the brain (Ren et al., 2015; Sano et al., 2015). In blowflies, injection of dmCCHa-2 increased proboscis extension in response to food (Ida et al., 2012). In pea aphids, reduction of CCHa-2 receptor caused changes in feeding behaviors, lower appetite and reduced feeding (Shahid et al., 2021). In the two-spotted cricket nymphs, reduction of CCHa-2 receptor increased while injection of CCHa-2 decreased food intake (Zhu et al., 2022). In the hematophagous kissing bug, reduction of CCHa-2 did not reduce the mass of blood ingested but increased post-prandial diuresis (Capriotti et al., 2019). Although the role of CCHa remains unknown in tick feeding behavior, there is growing cross-species evidence suggesting its necessity to feeding activity and food intake.

The invertebrate short NPF (sNPF) neuropeptide system is of significance to feeding in many animals which has a consensus sequence of xPxLRLR(F/W)amide (Nagata, 2016a), is conserved in Protostomia, but has limited sequence similarity with invertebrate NPF or vertebrate NPY peptides (Fadda et al., 2019; Nassel and Wegener, 2011) beyond the terminal RFamide. However, the receptors that recognize sNPF and NFP are distantly related, sharing a common ancestor with NPY receptors (Elphick et al., 2018). Insects have multiple peptides and receptors with differing roles depending upon the species (Fadda et al., 2019). In *Drosophila*, overexpression of sNPF increased feeding while reduction of sNPF decreased feeding (Lee et al., 2004). sNPFs regulate *Dilps* and body size (Lee et al., 2008). In mosquitos, sNPFs have a number of effects that promote feeding (Fadda et al., 2019) and small molecules targeted to an sNPF receptor alters blood feeding behaviors in mosquitos (Duvall et al., 2019). In honeybees, topical sNPF enhanced food intake and appetitive responses (Bestea et al., 2022), and stimulated locomotor activity akin to starvation in cockroaches (Mikani et al., 2015). In pea aphids, silencing sNPF or the receptor decreased stylet probing and delayed phloem ingestion (Amir et al., 2022). On the other hand, injection of sNPF decreased and reducing the receptor increased feeding in the desert locust, (Dillen et al., 2013) and injected sNPF delayed feeding in silk moth (Nagata et al., 2011). The second *Ixodes* NPY-related receptor, NPYLR2B, (Liesch et al., 2013) was not activated by sNPF in the HEK293T expression system. Among the possible explanations are: i) NPYLR2B binds another neuropeptide, rather than sNPF; ii) NPYLR2B does not couple to Gq but rather to another G protein; iii) NPYLR2B does not fold well or traffic to the plasma membrane. Further work is needed to deorphanize this receptor.

The arthropod sulfakinins (Nassel and Wu, 2022) with a sulfated tyrosine have central roles in food intake. In *Drosophila*, genetic manipulations showed that sulfakinins regulated food intake, starvation resistance and *dilp* expression (Soderberg et al., 2012) and other behaviors, such as odor preferences (Nichols et al., 2008) and aggression (Wu et al., 2020). In the red flour beetle, reducing sulfakinin (Yu et al., 2013) or sulfakinin receptor (Yu and Smagghe, 2014) by RNAi stimulated and injecting the peptide (Yu et al., 2013) reduced food intake. This behavior was also found in the kissing bug (Al-Alkawi et al., 2017; Bloom et al., 2019), brown planthopper (Guo et al., 2015), cricket (Meyering-Vos and Muller, 2007), cockroach (Maestro et al., 2001), desert locust (Wei et al., 2000), and blow fly (Downer et al., 2007). Sulfakinins also influence gut and heart contractions or regulate Neuropeptide F (which itself stimulates feeding in vertebrates) (Nassel and Wu, 2022).

Ticks spread pathogens to their hosts during active hematophagy, and thus transmission of pathogens could be interrupted by impairing this aspect of tick biology. An underdeveloped approach to combat disease spread is to target the tick’s central neural circuitry controlling feeding behaviors. A model of possible functions for the three neuropeptide systems suggested by studies in other organisms is stimulation of food intake by CCHa and sNPF and feeding suppression by sulfakinin (Suppl. Fig. 22).

Future experiments designed to test these hypotheses require that the neuropeptide signaling systems be further characterized. The experiments reported here develop the foundation for next steps aimed at discovering their role *in vivo* and to uncover agents that disrupt their function. In summary, we have provided new information on G protein signaling cascades in *Ixodes*, but further work is needed to identify the G proteins and effectors to which the CCHa, sNPF and SK receptors couple. Next steps to investigate any links of these neuropeptides signaling pathways to feeding behavior are necessary. Our results set the stage for further studies to define their physiological roles in feeding. Moreover, structural biology offers the possibility of protein-ligand or protein-protein interactions unique to ticks that could be targets for developing agents to disrupt feeding and disease transmission.

## Acknowledgements

We acknowledge the generous support of the Hoepner Family Fund, pilot grants from the Hendrick’s Fund and Center for Environmental Health and Medicine (SUNY Upstate Medical University), Dr. David Amberg (Office of VP for Research, SUNY Upstate Medical University), an Unrestricted Grant to the Department of Ophthalmology & Visual Science from Research to Prevent Blindness (RBP) and National Institutes of Health grants DK107944 (RJW), GM121621 (RJW) and GM134638 (LMS). We thank Drs. Jeffrey Amack and Jushua Wang (SUNY Upstate Medical University) and Leica Center of Excellence for Advanced Microscopy for assistance with the microscopy.

## Supplemental Data

**Supplemental Table 1**: Synthetic peptides used in calcium release assay.

**Supplemental Table 2**: PCR primers used for qRT-PCR and cDNA characterization.

**Supplemental Fig. 1**: GPCR coding regions used for expression in HEK293T cells.

**Supplemental Fig. 2**: Sequences of prepro-sNPF coding regions.

**Supplemental Fig. 3**: Sequences of sNPFR coding regions.

**Supplemental Fig. 4**: sNPFR amino acid sequences identity matrix.

**Supplemental Fig. 5**: *Ixodes scapularis* sNPFR Structural Models. **a**. A snake plot showing the 2D sequence of NPYLR1A transmembrane regions mapped onto the structure based upon human NPY2R. The amino and carboxyl termini and the intracellular loops (ICL) are omitted. Putative disulfide bond in the extracellular region is shown by the dashed line. The amino acids conserved between the two proteins are colored as indicated. The GPCR residue numbering by Ballesteros and Weinstein for each helix is shown in black. **b**. Structural models of sNPFR and NPYLR1B generated using Swiss-Model (swissmodel.expasy.org) and based upon PrRPR (9K27). A transverse view of the carbon backbone showing the transmembrane domains for all three models are closely aligned. The amino acids in the conserved ligand binding pocket (Fig. 2) are shown in stick representation (sNPFR in red and NPYR1B in blue) along with amino acids 28-31 of PrRP (VGRF-NH_2_) shown in gold with sphere representation. **c**. View of the ligand binding pocket from the extracellular face. The last five amino acid of sNPF (LRLRFNH_2_) modeled into the sNPFR are shown in green and two amino acids positions in the ligand binding pocket that differ between sNPFR and NPYL1B are shown. The potential steric conflict between Q71 and M252 and sNPF, which could prevent high affinity binding, are illustrated.

**Supplemental Fig. 6**: Sequences of prepro-SK coding regions

**Supplemental Fig. 7**: Alignment of prepro-SK PCR and genomic sequences

**Supplemental Fig. 8**: Sequences of SKR coding regions

**Supplemental Fig. 9**: SKR amino acid sequences identity matrix

**Supplemental Fig. 10**: Alignment of SKR Sequences

**Supplemental Fig. 11**: *Ixodes scapularis* SKR Structural Models. **a**. A snake plot showing the 2D sequence of SKR transmembrane regions mapped onto the structure based upon human CCKR. The amino and carboxyl termini and the intracellular loops (ICL) are omitted. Putative disulfide bond in the extracellular region is shown by the dashed line. The amino acids conserved between SKR and the two human CCKR proteins are colored as indicated. The GPCR residue numbering by Ballesteros and Weinstein for each helix is shown in black. **b**. Structural models of SKR generated using Swiss-Model (swissmodel.expasy.org) and based upon CCKAR (7EZM). A transverse view of the carbon backbone showing the transmembrane domains for both are closely aligned. The amino acids in the conserved ligand binding pocket (Fig. 5) are shown in stick representation (in red) along with amino acids of CCK8 (DY(SO_3_)MGWMDF-NH_2_) shown in gold with sphere representation. **c**. View of the ligand binding pocket from the extracellular face. The sulfated tyrosine from CCK8 modeled into the SKR is shown in green. The conserved amino acids predicted to interact with SK are shown in red.

**Supplemental Fig. 12**: Sequences of prepro-CCHa coding regions

**Supplemental Fig. 13**: Sequences of CCHaR coding regions

**Supplemental Fig. 14**: CCHaR amino acid sequence identity matrix

**Supplemental Fig. 15**: *Ixodes scapularis* CCHaR Structural Models. **a**. A snake plot showing the 2D sequence of CCHaR transmembrane regions mapped onto the structure based upon human ETH_B_R. The amino and carboxyl termini and the intracellular loops (ICL) are omitted. Putative disulfide bond in the extracellular region is shown by the dashed line. The amino acids conserved between SKR and the two human CCKR proteins are colored as indicated. **b**. Structural model of CCHaR generated using Swiss-Model (swissmodel.expasy.org) and based upon human ETH_B_R (5GLI). A transverse view of the carbon backbone showing the transmembrane domain and the extracellular face at the top. **c**. A model of CCHa including the disulfide bond was generated with PEP-FOLD2.1 (https://mobyle2.rpbs.univ-paris-diderot.fr/cgi-bin/portal.py#forms::PEP-FOLD). **d**. A model of CCHaR with CCHa in the binding pocket if shown (same orientation as in b. **e**. The amino acids predicted to be in the ligand binding pocket (Fig. 8) are shown in stick representation. CCHa has been removed for clarity.

**Supplemental Fig. 16**: Sequences of *Ixodes* GNA coding regions

**Supplemental Fig. 17**: Sequences of GNA coding regions used for expression

**Supplemental Fig. 18**: Sequences of *Ixodes* GNB coding regions

**Supplemental Fig. 19**: GNB amino acid sequence identity matrix between human and *Ixodes* subunit amino acids.

**Supplemental Fig. 20**: GNAQ-Kisspeptin Structure. **a**. Alignment of the human, mouse and *Ixodes scapularis* (XP_029846564) GNAQ subunit sequences, with amino acids matching the human sequence highlighted in blue. **b**. Alignment of the human and *Ixodes scapularis* (XP_002411384) GNB1 subunit sequences, with amino acids variant with the human sequence boxed and shaded in yellow, similar amino acid substitutions are shown in blue, non-conservative changes in red. **c**. The human kisspeptide receptor in complex with mouse Gq (8XGS) is shown. The amino acid positions that vary between the mouse and *Ixodes* G-protein subunit are highlighted with a sphere representation. The orientation of the kisspeptin receptor is opposite to the presentations in other figures, with the extracellular face at the bottom.

**Supplemental Fig. 21**: **a**. A comparison of sequences from the human phospholipaseC β3 and the closest *Ixodes* scapularis homolog (XP_029835062) coding regions. The conserved amino acids are in black, similar residues in blue and non-conservative changes in red. **b**. Structure of human phospholipaseC β3 and mouse GNAQ signaling complex (7SQ2). The amino acid positions that vary between the mouse and *Ixodes* GNAQ unit are highlighted with a sphere representation.

**Supplemental Fig. 22:** A schematic showing the possible roles of CCHa, sNPF and SK in tick feeding.

## References

1. Al-Alkawi, H., Lange, A.B., Orchard, I., 2017. Cloning, localization, and physiological effects of sulfakinin in the kissing bug, Rhodnius prolixus. Peptides 98, 15–22. 10.1016/j.peptides.2016.12.017.

2. Alasmari, S., Wall, R., 2020. Determining the total energy budget of the tick Ixodes ricinus. Exp Appl Acarol 80(4), 531–541. 10.1007/s10493-020-00479-1.

3. Alasmari, S., Wall, R., 2021. Metabolic rate and resource depletion in the tick Ixodes ricinus in response to temperature. Exp Appl Acarol 83(1), 81–93. 10.1007/s10493-020-00568-1.

4. Amir, M.B., Shi, Y., Cao, H., Ali, M.Y., Ahmed, M.A., Smagghe, G., Liu, T.X., 2022. Short Neuropeptide F and Its Receptor Regulate Feeding Behavior in Pea Aphid (Acyrthosiphon pisum). Insects 13(3). 10.3390/insects13030282.

5. Arora, S., Anubhuti, 2006. Role of neuropeptides in appetite regulation and obesity--a review. Neuropeptides 40(6), 375–401. 10.1016/j.npep.2006.07.001.

6. Atwood, B.K., Lopez, J., Wager-Miller, J., Mackie, K., Straiker, A., 2011. Expression of G protein-coupled receptors and related proteins in HEK293, AtT20, BV2, and N18 cell lines as revealed by microarray analysis. BMC Genomics 12, 14. 10.1186/1471-2164-12-14.

7. Belozerov, V.N., 2010. Seasonal adaptations in the life cycles of mites and ticks: comparative and evolutionary aspects, Trends in Acarology. pp. 319–326. 10.1007/978-90-481-9837-5_51.

8. Benoit, J.B., Denlinger, D.L., 2010. Meeting the challenges of on-host and off-host water balance in blood-feeding arthropods. J Insect Physiol 56(10), 1366–1376. 10.1016/j.jinsphys.2010.02.014.

9. Bestea, L., Paoli, M., Arrufat, P., Ronsin, B., Carcaud, J., Sandoz, J.C., Velarde, R., Giurfa, M., de Brito Sanchez, M.G., 2022. The short neuropeptide F regulates appetitive but not aversive responsiveness in a social insect. iScience 25(1), 103619. 10.1016/j.isci.2021.103619.

10. Bhumika, A.K.S., 2018. Regulation of feeding behavior in Drosophila through the interplay of gustation, physiology and neuromodulation. Front Biosci, Landmark, 23, 2016–2027.

11. Bloom, M., Lange, A.B., Orchard, I., 2019. Identification, Functional Characterization, and Pharmacological Analysis of Two Sulfakinin Receptors in the Medically-Important Insect Rhodnius prolixus. Sci Rep 9(1), 13437. 10.1038/s41598-019-49790-x.

12. Caers, J., Peymen, K., Suetens, N., Temmerman, L., Janssen, T., Schoofs, L., Beets, I., 2014. Characterization of G protein-coupled receptors by a fluorescence-based calcium mobilization assay. J Vis Exp (89), e51516. 10.3791/51516.

13. Caers, J., Verlinden, H., Zels, S., Vandersmissen, H.P., Vuerinckx, K., Schoofs, L., 2012. More than two decades of research on insect neuropeptide GPCRs: an overview. Front Endocrinol (Lausanne) 3, 151. 10.3389/fendo.2012.00151.

14. Capriotti, N., Ianowski, J.P., Gioino, P., Ons, S., 2019. The neuropeptide CCHamide2 regulates diuresis in the Chagas disease vector Rhodnius prolixus. J Exp Biol 222(Pt 10). 10.1242/jeb.203000.

15. Cassens, J., Oliva Chavez, A.S., Tufts, D.M., Zhong, J., Faulk, C., Oliver, J.D., 2025. Whole Genome Sequencing Reveals Clade-Specific Genetic Variation in Blacklegged Ticks. Ecol Evol 15(2), e70987. 10.1002/ece3.70987.

16. Christie, A.E., 2008. Neuropeptide discovery in Ixodoidea: an in silico investigation using publicly accessible expressed sequence tags. Gen Comp Endocrinol 157(2), 174–185. 10.1016/j.ygcen.2008.03.027.

17. De, S., Kingan, S.B., Kitsou, C., Portik, D.M., Foor, S.D., Frederick, J.C., Rana, V.S., Paulat, N.S., Ray, D.A., Wang, Y., Glenn, T.C., Pal, U., 2023. A high-quality Ixodes scapularis genome advances tick science. Nat Genet 55(2), 301–311. 10.1038/s41588-022-01275-w.

18. Dillen, S., Zels, S., Verlinden, H., Spit, J., Van Wielendaele, P., Vanden Broeck, J., 2013. Functional characterization of the short neuropeptide F receptor in the desert locust, Schistocerca gregaria. PLoS One 8(1), e53604. 10.1371/journal.pone.0053604.

19. Ding, Y., Zhang, H., Liao, Y.Y., Chen, L.N., Ji, S.Y., Qin, J., Mao, C., Shen, D.D., Lin, L., Wang, H., Zhang, Y., Li, X.M., 2022. Structural insights into human brain-gut peptide cholecystokinin receptors. Cell Discov 8(1), 55. 10.1038/s41421-022-00420-3.

20. Dinglasan, R.R., Liesch, J., Bellani, L.L., Vosshall, L.B., 2013. Functional and Genetic Characterization of Neuropeptide Y-Like Receptors in Aedes aegypti. PLoS Neglected Tropical Diseases 7(10). 10.1371/journal.pntd.0002486.

21. Downer, K.E., Haselton, A.T., Nachman, R.J., Stoffolano, J.G., Jr., 2007. Insect satiety: sulfakinin localization and the effect of drosulfakinin on protein and carbohydrate ingestion in the blow fly, Phormia regina (Diptera: Calliphoridae). J Insect Physiol 53(1), 106–112. 10.1016/j.jinsphys.2006.10.013.

22. Duvall, L.B., Ramos-Espiritu, L., Barsoum, K.E., Glickman, J.F., Vosshall, L.B., 2019. Small-Molecule Agonists of Ae. aegypti Neuropeptide Y Receptor Block Mosquito Biting. Cell 176(4), 687–701 e685. 10.1016/j.cell.2018.12.004.

23. Egekwu, N., Sonenshine, D.E., Bissinger, B.W., Roe, R.M., 2014. Transcriptome of the female synganglion of the black-legged tick Ixodes scapularis (Acari: Ixodidae) with comparison between Illumina and 454 systems. PLoS One 9(7), e102667. 10.1371/journal.pone.0102667.

24. Eisen, L., 2018. Pathogen transmission in relation to duration of attachment by Ixodes scapularis ticks. Ticks Tick Borne Dis 9(3), 535–542. 10.1016/j.ttbdis.2018.01.002.

25. Eisen, R.J., Eisen, L., 2018. The Blacklegged Tick, Ixodes scapularis: An Increasing Public Health Concern. Trends Parasitol 34(4), 295–309. 10.1016/j.pt.2017.12.006.

26. Eisen, R.J., Kugeler, K.J., Eisen, L., Beard, C.B., Paddock, C.D., 2017. Tick-Borne Zoonoses in the United States: Persistent and Emerging Threats to Human Health. ILAR J 58(3), 319–335. 10.1093/ilar/ilx005.

27. Elphick, M.R., Mirabeau, O., Larhammar, D., 2018. Evolution of neuropeptide signalling systems. J Exp Biol 221(Pt 19). 10.1242/jeb.193342.

28. Fadda, M., Hasakiogullari, I., Temmerman, L., Beets, I., Zels, S., Schoofs, L., 2019. Regulation of Feeding and Metabolism by Neuropeptide F and Short Neuropeptide F in Invertebrates. Front Endocrinol (Lausanne) 10, 64. 10.3389/fendo.2019.00064.

29. Farhan, A., Gulati, J., Grobetae-Wilde, E., Vogel, H., Hansson, B.S., Knaden, M., 2013. The CCHamide 1 receptor modulates sensory perception and olfactory behavior in starved Drosophila. Sci Rep 3, 2765. 10.1038/srep02765.

30. Geary, N., 2014. A physiological perspective on the neuroscience of eating. Physiol Behav 136, 3–14. 10.1016/j.physbeh.2014.03.022.

31. Giannone, F., Malpeli, G., Lisi, V., Grasso, S., Shukla, P., Ramarli, D., Sartoris, S., Monsurro, V., Krampera, M., Amato, E., Tridente, G., Colombatti, M., Parenti, M., Innamorati, G., 2010. The puzzling uniqueness of the heterotrimeric G15 protein and its potential beyond hematopoiesis. J Mol Endocrinol 44(5), 259–269. 10.1677/JME-09-0134.

32. Gulia-Nuss, M., Nuss, A.B., Meyer, J.M., Sonenshine, D.E., Roe, R.M., Waterhouse, R.M., Sattelle, D.B., de la Fuente, J., Ribeiro, J.M., Megy, K., Thimmapuram, J., Miller, J.R., Walenz, B.P., Koren, S., Hostetler, J.B., Thiagarajan, M., Joardar, V.S., Hannick, L.I., Bidwell, S., Hammond, M.P., Young, S., Zeng, Q., Abrudan, J.L., Almeida, F.C., Ayllon, N., Bhide, K., Bissinger, B.W., Bonzon-Kulichenko, E., Buckingham, S.D., Caffrey, D.R., Caimano, M.J., Croset, V., Driscoll, T., Gilbert, D., Gillespie, J.J., Giraldo-Calderon, G.I., Grabowski, J.M., Jiang, D., Khalil, S.M., Kim, D., Kocan, K.M., Koci, J., Kuhn, R.J., Kurtti, T.J., Lees, K., Lang, E.G., Kennedy, R.C., Kwon, H., Perera, R., Qi, Y., Radolf, J.D., Sakamoto, J.M., Sanchez-Gracia, A., Severo, M.S., Silverman, N., Simo, L., Tojo, M., Tornador, C., Van Zee, J.P., Vazquez, J., Vieira, F.G., Villar, M., Wespiser, A.R., Yang, Y., Zhu, J., Arensburger, P., Pietrantonio, P.V., Barker, S.C., Shao, R., Zdobnov, E.M., Hauser, F., Grimmelikhuijzen, C.J., Park, Y., Rozas, J., Benton, R., Pedra, J.H., Nelson, D.R., Unger, M.F., Tubio, J.M., Tu, Z., Robertson, H.M., Shumway, M., Sutton, G., Wortman, J.R., Lawson, D., Wikel, S.K., Nene, V.M., Fraser, C.M., Collins, F.H., Birren, B., Nelson, K.E., Caler, E., Hill, C.A., 2016. Genomic insights into the Ixodes scapularis tick vector of Lyme disease. Nat Commun 7, 10507. 10.1038/ncomms10507.

33. Guo, M., Qu, X., Qin, X.Q., 2015. Bombesin-like peptides and their receptors: recent findings in pharmacology and physiology. Curr Opin Endocrinol Diabetes Obes 22(1), 3–8. 10.1097/MED.0000000000000126.

34. Hansen, K.B., Brauner-Osborne, H., 2009. FLIPR assays of intracellular calcium in GPCR drug discovery. Methods Mol Biol 552, 269–278. 10.1007/978-1-60327-317-6_19.

35. Hansen, K.K., Hauser, F., Williamson, M., Weber, S.B., Grimmelikhuijzen, C.J., 2011. The Drosophila genes CG14593 and CG30106 code for G-protein-coupled receptors specifically activated by the neuropeptides CCHamide-1 and CCHamide-2. Biochem Biophys Res Commun 404(1), 184–189. 10.1016/j.bbrc.2010.11.089.

36. Herrmann, C., Gern, L., 2012. Do the level of energy reserves, hydration status and Borrelia infection influence walking by Ixodes ricinus (Acari: Ixodidae) ticks? Parasitology 139(3), 330–337. 10.1017/S0031182011002095.

37. Hook, V., Lietz, C.B., Podvin, S., Cajka, T., Fiehn, O., 2018. Diversity of Neuropeptide Cell-Cell Signaling Molecules Generated by Proteolytic Processing Revealed by Neuropeptidomics Mass Spectrometry. J Am Soc Mass Spectrom 29(5), 807–816. 10.1007/s13361-018-1914-1.

38. Ida, T., Takahashi, T., Tominaga, H., Sato, T., Sano, H., Kume, K., Ozaki, M., Hiraguchi, T., Shiotani, H., Terajima, S., Nakamura, Y., Mori, K., Yoshida, M., Kato, J., Murakami, N., Miyazato, M., Kangawa, K., Kojima, M., 2012. Isolation of the bioactive peptides CCHamide-1 and CCHamide-2 from Drosophila and their putative role in appetite regulation as ligands for G protein-coupled receptors. Front Endocrinol (Lausanne) 3, 177. 10.3389/fendo.2012.00177.

39. Isberg, V., de Graaf, C., Bortolato, A., Cherezov, V., Katritch, V., Marshall, F.H., Mordalski, S., Pin, J.P., Stevens, R.C., Vriend, G., Gloriam, D.E., 2015. Generic GPCR residue numbers – aligning topology maps while minding the gaps. Trends Pharmacol Sci 36(1), 22–31. 10.1016/j.tips.2014.11.001.

40. Jekely, G., 2013. Global view of the evolution and diversity of metazoan neuropeptide signaling. Proc Natl Acad Sci U S A 110(21), 8702–8707. 10.1073/pnas.1221833110.

41. Jekely, G., Melzer, S., Beets, I., Kadow, I.C.G., Koene, J., Haddad, S., Holden-Dye, L., 2018. The long and the short of it – a perspective on peptidergic regulation of circuits and behaviour. J Exp Biol 221(Pt 3). 10.1242/jeb.166710.

42. Ji, Y., Duan, J., Yuan, Q., He, X., Yang, G., Zhu, S., Wu, K., Hu, W., Gao, T., Cheng, X., Jiang, H., Eric Xu, H., Jiang, Y., 2023. Structural basis of peptide recognition and activation of endothelin receptors. Nat Commun 14(1), 1268. 10.1038/s41467-023-36998-9.

43. Kang, H., Park, C., Choi, Y.K., Bae, J., Kwon, S., Kim, J., Choi, C., Seok, C., Im, W., Choi, H.J., 2023. Structural basis for Y2 receptor-mediated neuropeptide Y and peptide YY signaling. Structure 31(1), 44–57 e46. 10.1016/j.str.2022.11.010.

44. Kersigo, J., Pan, N., Lederman, J.D., Chatterjee, S., Abel, T., Pavlinkova, G., Silos-Santiago, I., Fritzsch, B., 2018. A RNAscope whole mount approach that can be combined with immunofluorescence to quantify differential distribution of mRNA. Cell Tissue Res 374(2), 251–262. 10.1007/s00441-018-2864-4.

45. Koci, J., Simo, L., Park, Y., 2013. Validation of internal reference genes for real-time quantitative polymerase chain reaction studies in the tick, Ixodes scapularis (Acari: Ixodidae). J Med Entomol 50(1), 79–84. 10.1603/me12034.

46. Kooistra, A.J., Mordalski, S., Pandy-Szekeres, G., Esguerra, M., Mamyrbekov, A., Munk, C., Keseru, G.M., Gloriam, D.E., 2021. GPCRdb in 2021: integrating GPCR sequence, structure and function. Nucleic Acids Res 49(D1), D335–D343. 10.1093/nar/gkaa1080.

47. Kugeler, K.J., Schwartz, A.M., Delorey, M.J., Mead, P.S., Hinckley, A.F., 2021. Estimating the Frequency of Lyme Disease Diagnoses, United States, 2010-2018. Emerg Infect Dis 27(2), 616–619. 10.3201/eid2702.202731.

48. Kumar, D., Mains, R.E., Eipper, B.A., 2016. 60 YEARS OF POMC: From POMC and alpha-MSH to PAM, molecular oxygen, copper, and vitamin C. J Mol Endocrinol 56(4), T63–76. 10.1530/JME-15-0266.

49. Kumar, S., Stecher, G., Li, M., Knyaz, C., Tamura, K., 2018. MEGA X: Molecular Evolutionary Genetics Analysis across Computing Platforms. Mol Biol Evol 35(6), 1547–1549. 10.1093/molbev/msy096.

50. Langley, D.B., Schofield, P., Jackson, J., Herzog, H., Christ, D., 2022. Crystal structures of human neuropeptide Y (NPY) and peptide YY (PYY). Neuropeptides 92, 102231. 10.1016/j.npep.2022.102231.

51. Larkin, M.A., Blackshields, G., Brown, N.P., Chenna, R., McGettigan, P.A., McWilliam, H., Valentin, F., Wallace, I.M., Wilm, A., Lopez, R., Thompson, J.D., Gibson, T.J., Higgins, D.G., 2007. Clustal W and Clustal X version 2.0. Bioinformatics 23(21), 2947–2948. 10.1093/bioinformatics/btm404.

52. Lee, K.S., Kwon, O.Y., Lee, J.H., Kwon, K., Min, K.J., Jung, S.A., Kim, A.K., You, K.H., Tatar, M., Yu, K., 2008. Drosophila short neuropeptide F signalling regulates growth by ERK-mediated insulin signalling. Nat Cell Biol 10(4), 468–475. 10.1038/ncb1710.

53. Lee, K.S., You, K.H., Choo, J.K., Han, Y.M., Yu, K., 2004. Drosophila short neuropeptide F regulates food intake and body size. J Biol Chem 279(49), 50781–50789. 10.1074/jbc.M407842200.

54. Lees, A.D., Milne, A., 1951. The seasonal and diurnal activities of individual sheep ticks (Ixodes ricinus L). Parasitology 41(3-4), 189–208. 10.1017/s0031182000084031.

55. Li, S., Torre-Muruzabal, T., Sogaard, K.C., Ren, G.R., Hauser, F., Engelsen, S.M., Podenphanth, M.D., Desjardins, A., Grimmelikhuijzen, C.J., 2013. Expression patterns of the Drosophila neuropeptide CCHamide-2 and its receptor may suggest hormonal signaling from the gut to the brain. PLoS One 8(10), e76131. 10.1371/journal.pone.0076131.

56. Li, Y., Yuan, Q., He, X., Zhang, Y., You, C., Wu, C., Li, J., Xu, H.E., Zhao, L.H., 2024. Molecular mechanism of prolactin-releasing peptide recognition and signaling via its G protein-coupled receptor. Cell Discov 10(1), 91. 10.1038/s41421-024-00724-6.

57. Liesch, J., Bellani, L.L., Vosshall, L.B., 2013. Functional and genetic characterization of neuropeptide Y-like receptors in Aedes aegypti. PLoS Negl Trop Dis 7(10), e2486. 10.1371/journal.pntd.0002486.

58. Lin, S., Senapati, B., Tsao, C.H., 2019. Neural basis of hunger-driven behaviour in Drosophila. Open Biol 9(3), 180259. 10.1098/rsob.180259.

59. Lyon, A.M., Dutta, S., Boguth, C.A., Skiniotis, G., Tesmer, J.J., 2013. Full-length Galpha(q)-phospholipase C-beta3 structure reveals interfaces of the C-terminal coiled-coil domain. Nat Struct Mol Biol 20(3), 355–362. 10.1038/nsmb.2497.

60. Maestro, J.L., Aguilar, R., Pascual, N., Valero, M.L., Piulachs, M.D., Andreu, D., Navarro, I., Belles, X., 2001. Screening of antifeedant activity in brain extracts led to the identification of sulfakinin as a satiety promoter in the German cockroach. Are arthropod sulfakinins homologous to vertebrate gastrins-cholecystokinins? Eur J Biochem 268(22), 5824–5830. 10.1046/j.0014-2956.2001.02527.x.

61. Masuho, I., Kise, R., Gainza, P., Von Moo, E., Li, X., Tany, R., Wakasugi-Masuho, H., Correia, B.E., Martemyanov, K.A., 2023. Rules and mechanisms governing G protein coupling selectivity of GPCRs. Cell Rep 42(10), 113173. 10.1016/j.celrep.2023.113173.

62. Meyering-Vos, M., Muller, A., 2007. RNA interference suggests sulfakinins as satiety effectors in the cricket Gryllus bimaculatus. J Insect Physiol 53(8), 840–848. 10.1016/j.jinsphys.2007.04.003.

63. Mikani, A., Watari, Y., Takeda, M., 2015. Brain-midgut cross-talk and autocrine metabolastat via the sNPF/CCAP negative feed-back loop in the American cockroach, Periplaneta americana. Cell Tissue Res 362(3), 481–496. 10.1007/s00441-015-2242-4.

64. Miller, J.R., Koren, S., Dilley, K.A., Harkins, D.M., Stockwell, T.B., Shabman, R.S., Sutton, G.G., 2018. A draft genome sequence for the Ixodes scapularis cell line, ISE6. F1000Res 7, 297. 10.12688/f1000research.13635.1.

65. Millington, J.W., Brownrigg, G.P., Chao, C., Sun, Z., Basner-Collins, P.J., Wat, L.W., Hudry, B., Miguel-Aliaga, I., Rideout, E.J., 2021. Female-biased upregulation of insulin pathway activity mediates the sex difference in Drosophila body size plasticity. Elife 10. 10.7554/eLife.58341.

66. Mirabeau, O., Joly, J.S., 2013. Molecular evolution of peptidergic signaling systems in bilaterians. Proc Natl Acad Sci U S A 110(22), E2028–2037. 10.1073/pnas.1219956110.

67. Mobbs, J.I., Belousoff, M.J., Harikumar, K.G., Piper, S.J., Xu, X., Furness, S.G.B., Venugopal, H., Christopoulos, A., Danev, R., Wootten, D., Thal, D.M., Miller, L.J., Sexton, P.M., 2021. Structures of the human cholecystokinin 1 (CCK1) receptor bound to Gs and Gq mimetic proteins provide insight into mechanisms of G protein selectivity. PLoS Biol 19(6), e3001295. 10.1371/journal.pbio.3001295.

68. Morishita, F., Minakata, H., 2016. Chapter 75 – GGNG Peptides, in: Takei, Y., Ando, H., Tsutsui, K. (Eds.), Handbook of Hormones. Academic Press, San Diego, pp. 454–456. 10.1016/B978-0-12-801028-0.00075-1.

69. Nagata, S., 2016a. Chapter 42 – Short Neuropeptide F, in: Takei, Y., Ando, H., Tsutsui, K. (Eds.), Handbook of Hormones. Academic Press, San Diego, pp. 357–358. 10.1016/B978-0-12-801028-0.00042-8.

70. Nagata, S., 2016b. Chapter 84 – CCHamide, in: Takei, Y., Ando, H., Tsutsui, K. (Eds.), Handbook of Hormones. Academic Press, San Diego, pp. 475–476. 10.1016/B978-0-12-801028-0.00084-2.

71. Nagata, S., Morooka, N., Matsumoto, S., Kawai, T., Nagasawa, H., 2011. Effects of neuropeptides on feeding initiation in larvae of the silkworm, Bombyx mori. Gen Comp Endocrinol 172(1), 90–95. 10.1016/j.ygcen.2011.03.004.

72. Nassel, D.R., Wegener, C., 2011. A comparative review of short and long neuropeptide F signaling in invertebrates: Any similarities to vertebrate neuropeptide Y signaling? Peptides 32(6), 1335–1355. 10.1016/j.peptides.2011.03.013.

73. Nassel, D.R., Wu, S.F., 2022. Cholecystokinin/sulfakinin peptide signaling: conserved roles at the intersection between feeding, mating and aggression. Cell Mol Life Sci 79(3), 188. 10.1007/s00018-022-04214-4.

74. Nassel, D.R., Zandawala, M., 2019. Recent advances in neuropeptide signaling in Drosophila, from genes to physiology and behavior. Prog Neurobiol 179, 101607. 10.1016/j.pneurobio.2019.02.003.

75. Needham, G.R., Teel, P.D., 1991. Off-host physiological ecology of ixodid ticks. Annu Rev Entomol 36, 659–681. 10.1146/annurev.en.36.010191.003303.

76. Nelson, C.A., Saha, S., Kugeler, K.J., Delorey, M.J., Shankar, M.B., Hinckley, A.F., Mead, P.S., 2015. Incidence of Clinician-Diagnosed Lyme Disease, United States, 2005-2010. Emerg Infect Dis 21(9), 1625–1631. 10.3201/eid2109.150417.

77. Neupert, S., Russell, W.K., Predel, R., Russell, D.H., Strey, O.F., Teel, P.D., Nachman, R.J., 2009. The neuropeptidomics of Ixodes scapularis synganglion. J Proteomics 72(6), 1040–1045. 10.1016/j.jprot.2009.06.007.

78. Nichols, R., Egle, J.P., Langan, N.R., Palmer, G.C., 2008. The different effects of structurally related sulfakinins on Drosophila melanogaster odor preference and locomotion suggest involvement of distinct mechanisms. Peptides 29(12), 2128–2135. 10.1016/j.peptides.2008.08.010.

79. Nusbaum, M.P., Blitz, D.M., Marder, E., 2017. Functional consequences of neuropeptide and small-molecule co-transmission. Nat Rev Neurosci 18(7), 389–403. 10.1038/nrn.2017.56.

80. Offermanns, S., Simon, M.I., 1995. G alpha 15 and G alpha 16 couple a wide variety of receptors to phospholipase C. J Biol Chem 270(25), 15175–15180. 10.1074/jbc.270.25.15175.

81. Ogden, N.H., Lindsay, L.R., Beauchamp, G., Charron, D., Maarouf, A., O’Callaghan, C.J., Waltner-Toews, D., Barker, I.K., 2004. Investigation of relationships between temperature and developmental rates of tick Ixodes scapularis (Acari: Ixodidae) in the laboratory and field. J Med Entomol 41(4), 622–633. 10.1603/0022-2585-41.4.622.

82. Oldham, W.M., Hamm, H.E., 2008. Heterotrimeric G protein activation by G-protein-coupled receptors. Nat Rev Mol Cell Biol 9(1), 60–71. 10.1038/nrm2299.

83. Oliver, J.H., 1989. Biology and Systematics of Ticks (Acari:Ixodida). Annual Review of Ecology and Systematics 20, 397–430.

84. Park, C., Kim, J., Ko, S.B., Choi, Y.K., Jeong, H., Woo, H., Kang, H., Bang, I., Kim, S.A., Yoon, T.Y., Seok, C., Im, W., Choi, H.J., 2022. Structural basis of neuropeptide Y signaling through Y1 receptor. Nat Commun 13(1), 853. 10.1038/s41467-022-28510-6.

85. Pool, A.H., Scott, K., 2014. Feeding regulation in Drosophila. Curr Opin Neurobiol 29, 57–63. 10.1016/j.conb.2014.05.008.

86. Randolf, S.E., 2008. The impact of tick ecology on pathogen transmission dynamics, Ticks: Biology, Disease and Control, edited by Alan S. Bowman, and Patricia A. Nuttall. pp. 40–72.

87. Randolph, S.E., 1998. Ticks are not Insects: Consequences of Contrasting Vector Biology for Transmission Potential. Parasitol Today 14(5), 186–192.

88. Randolph, S.E., Green, R.M., Hoodless, A.N., Peacey, M.F., 2002. An empirical quantitative framework for the seasonal population dynamics of the tick Ixodes ricinus. Int J Parasitol 32(8), 979–989. 10.1016/s0020-7519(02)00030-9.

89. Ren, G.R., Hauser, F., Rewitz, K.F., Kondo, S., Engelbrecht, A.F., Didriksen, A.K., Schjott, S.R., Sembach, F.E., Li, S., Sogaard, K.C., Sondergaard, L., Grimmelikhuijzen, C.J., 2015. CCHamide-2 Is an Orexigenic Brain-Gut Peptide in Drosophila. PLoS One 10(7), e0133017. 10.1371/journal.pone.0133017.

90. Sano, H., Nakamura, A., Texada, M.J., Truman, J.W., Ishimoto, H., Kamikouchi, A., Nibu, Y., Kume, K., Ida, T., Kojima, M., 2015. The Nutrient-Responsive Hormone CCHamide-2 Controls Growth by Regulating Insulin-like Peptides in the Brain of Drosophila melanogaster. PLoS Genet 11(5), e1005209. 10.1371/journal.pgen.1005209.

91. Schindelin, J., Arganda-Carreras, I., Frise, E., Kaynig, V., Longair, M., Pietzsch, T., Preibisch, S., Rueden, C., Saalfeld, S., Schmid, B., Tinevez, J.Y., White, D.J., Hartenstein, V., Eliceiri, K., Tomancak, P., Cardona, A., 2012. Fiji: an open-source platform for biological-image analysis. Nat Methods 9(7), 676–682. 10.1038/nmeth.2019.

92. Schmittgen, T.D., Livak, K.J., 2008. Analyzing real-time PCR data by the comparative C(T) method. Nat Protoc 3(6), 1101–1108. 10.1038/nprot.2008.73.

93. Schoofs, L., De Loof, A., Van Hiel, M.B., 2017. Neuropeptides as Regulators of Behavior in Insects. Annu Rev Entomol 62, 35–52. 10.1146/annurev-ento-031616-035500.

94. Schrage, R., Schmitz, A.L., Gaffal, E., Annala, S., Kehraus, S., Wenzel, D., Bullesbach, K.M., Bald, T., Inoue, A., Shinjo, Y., Galandrin, S., Shridhar, N., Hesse, M., Grundmann, M., Merten, N., Charpentier, T.H., Martz, M., Butcher, A.J., Slodczyk, T., Armando, S., Effern, M., Namkung, Y., Jenkins, L., Horn, V., Stossel, A., Dargatz, H., Tietze, D., Imhof, D., Gales, C., Drewke, C., Muller, C.E., Holzel, M., Milligan, G., Tobin, A.B., Gomeza, J., Dohlman, H.G., Sondek, J., Harden, T.K., Bouvier, M., Laporte, S.A., Aoki, J., Fleischmann, B.K., Mohr, K., Konig, G.M., Tuting, T., Kostenis, E., 2015. The experimental power of FR900359 to study Gq-regulated biological processes. Nat Commun 6, 10156. 10.1038/ncomms10156.

95. Schulze, T.L., Jordan, R.A., Hung, R.W., 2001. Effects of selected meterological factors on diurnal questing of Ixodes scapularis and Amblyomma Americanum (Acari: Ixodidade). J Med Entomol 38(2), 318–324. 10.1603/0022-2585-38.2.318.

96. Shahid, S., Shi, Y., Yang, C., Li, J., Ali, M.Y., Smagghe, G., Liu, T.X., 2021. CCHamide2-receptor regulates feeding behavior in the pea aphid, Acyrthosiphon pisum. Peptides 143, 170596. 10.1016/j.peptides.2021.170596.

97. Shen, S., Deng, Y., Shen, C., Chen, H., Cheng, L., Wu, C., Zhao, C., Yang, Z., Hou, H., Wang, K., Shao, Z., Deng, C., Ye, F., Yan, W., 2024. Structural basis of neuropeptide Y signaling through Y(1) and Y(2) receptors. MedComm (2020) 5(7), e565. 10.1002/mco2.565.

98. Shihoya, W., Izume, T., Inoue, A., Yamashita, K., Kadji, F.M.N., Hirata, K., Aoki, J., Nishizawa, T., Nureki, O., 2018. Crystal structures of human ET(B) receptor provide mechanistic insight into receptor activation and partial activation. Nat Commun 9(1), 4711. 10.1038/s41467-018-07094-0.

99. Shihoya, W., Nishizawa, T., Yamashita, K., Inoue, A., Hirata, K., Kadji, F.M.N., Okuta, A., Tani, K., Aoki, J., Fujiyoshi, Y., Doi, T., Nureki, O., 2017. X-ray structures of endothelin ET(B) receptor bound to clinical antagonist bosentan and its analog. Nat Struct Mol Biol 24(9), 758–764. 10.1038/nsmb.3450.

100. Smith, W.W., Thomas, J., Liu, J., Li, T., Moran, T.H., 2014. From fat fruit fly to human obesity. Physiol Behav 136, 15–21. 10.1016/j.physbeh.2014.01.017.

101. Sobrino Crespo, C., Perianes Cachero, A., Puebla Jimenez, L., Barrios, V., Arilla Ferreiro, E., 2014. Peptides and food intake. Front Endocrinol (Lausanne) 5, 58. 10.3389/fendo.2014.00058.

102. Soderberg, J.A., Carlsson, M.A., Nassel, D.R., 2012. Insulin-Producing Cells in the Drosophila Brain also Express Satiety-Inducing Cholecystokinin-Like Peptide, Drosulfakinin. Front Endocrinol (Lausanne) 3, 109. 10.3389/fendo.2012.00109.

103. Spielman, A., Wilson, M.L., Levine, J.F., Piesman, J., 1985. Ecology of Ixodes dammini-borne human babesiosis and Lyme disease. Annu Rev Entomol 30, 439–460. 10.1146/annurev.en.30.010185.002255.

104. Stecher, G., Tamura, K., Kumar, S., 2020. Molecular Evolutionary Genetics Analysis (MEGA) for macOS. Mol Biol Evol 37(4), 1237–1239. 10.1093/molbev/msz312.

105. Szczesniak, L.M., Bonzerato, C.G., Schulman, J.J., Bah, A., Wojcikiewicz, R.J.H., 2021. Bok binds to a largely disordered loop in the coupling domain of type 1 inositol 1,4,5-trisphosphate receptor. Biochem Biophys Res Commun 553, 180–186. 10.1016/j.bbrc.2021.03.047.

106. Tang, T., Hartig, C., Chen, Q., Zhao, W., Kaiser, A., Zhang, X., Zhang, H., Qu, H., Yi, C., Ma, L., Han, S., Zhao, Q., Beck-Sickinger, A.G., Wu, B., 2021. Structural basis for ligand recognition of the neuropeptide Y Y(2) receptor. Nat Commun 12(1), 737. 10.1038/s41467-021-21030-9.

107. Tang, T., Tan, Q., Han, S., Diemar, A., Lobner, K., Wang, H., Schuss, C., Behr, V., Morl, K., Wang, M., Chu, X., Yi, C., Keller, M., Kofoed, J., Reedtz-Runge, S., Kaiser, A., Beck-Sickinger, A.G., Zhao, Q., Wu, B., 2022. Receptor-specific recognition of NPY peptides revealed by structures of NPY receptors. Sci Adv 8(18), eabm1232. 10.1126/sciadv.abm1232.

108. Teufel, F., Almagro Armenteros, J.J., Johansen, A.R., Gislason, M.H., Pihl, S.I., Tsirigos, K.D., Winther, O., Brunak, S., von Heijne, G., Nielsen, H., 2022a. SignalP 6.0 predicts all five types of signal peptides using protein language models. Nat Biotechnol. 10.1038/s41587-021-01156-3.

109. Teufel, F., Almagro Armenteros, J.J., Johansen, A.R., Gislason, M.H., Pihl, S.I., Tsirigos, K.D., Winther, O., Brunak, S., von Heijne, G., Nielsen, H., 2022b. SignalP 6.0 predicts all five types of signal peptides using protein language models. Nat Biotechnol 40(7), 1023–1025. 10.1038/s41587-021-01156-3.

110. Thomas, C.E., Burton, E.S., Brunner, J.L., 2019. Environmental Drivers of Questing Activity of Juvenile Black-Legged Ticks (Acari: Ixodidae): Temperature, Desiccation Risk, and Diel Cycles. J Med Entomol. 10.1093/jme/tjz126.

111. Veenstra, J.A., 2000. Mono– and dibasic proteolytic cleavage sites in insect neuroendocrine peptide precursors. Arch Insect Biochem Physiol 43(2), 49–63. 10.1002/(SICI)1520-6327(200002)43:2<49::AID-ARCH1>3.0.CO;2-M.

112. Waladde, S.M., Rice, M.J., 1982. The Sensory Basis of Tick Feeding Behaviour, in: Obenchain, F.D., Galun, R. (Eds.), Physiology of Ticks. Pergamon Press, New York, pp. 71–118.

113. Waldman, J., Xavier, M.A., Vieira, L.R., Logullo, R., Braz, G.R.C., Tirloni, L., Ribeiro, J.M.C., Veenstra, J.A., Silva Vaz, I.D., Jr., 2022. Neuropeptides in Rhipicephalus microplus and other hard ticks. Ticks Tick Borne Dis 13(3), 101910. 10.1016/j.ttbdis.2022.101910.

114. Wang, Y., Wang, M., Yin, S., Jang, R., Wang, J., Xue, Z., Xu, T., 2015. NeuroPep: a comprehensive resource of neuropeptides. Database (Oxford) 2015, bav038. 10.1093/database/bav038.

115. Wei, Z., Baggerman, G., R, J.N., Goldsworthy, G., Verhaert, P., De Loof, A., Schoofs, L., 2000. Sulfakinins reduce food intake in the desert locust, Schistocerca gregaria. J Insect Physiol 46(9), 1259–1265. 10.1016/s0022-1910(00)00046-9.

116. Wu, F., Deng, B., Xiao, N., Wang, T., Li, Y., Wang, R., Shi, K., Luo, D.G., Rao, Y., Zhou, C., 2020. A neuropeptide regulates fighting behavior in Drosophila melanogaster. Elife 9. 10.7554/eLife.54229.

117. Wu, Z., Chen, G., Qiu, C., Yan, X., Xu, L., Jiang, S., Xu, J., Han, R., Shi, T., Liu, Y., Gao, W., Wang, Q., Li, J., Ye, F., Pan, X., Zhang, Z., Ning, P., Zhang, B., Chen, J., Du, Y., 2024. Structural basis for the ligand recognition and G protein subtype selectivity of kisspeptin receptor. Sci Adv 10(33), eadn7771. 10.1126/sciadv.adn7771.

118. Yu, N., Nachman, R.J., Smagghe, G., 2013. Characterization of sulfakinin and sulfakinin receptor and their roles in food intake in the red flour beetle Tribolium castaneum. Gen Comp Endocrinol 188, 196–203. 10.1016/j.ygcen.2013.03.006.

119. Yu, N., Smagghe, G., 2014. Characterization of sulfakinin receptor 2 and its role in food intake in the red flour beetle, Tribolium castaneum. Peptides 53, 232–237. 10.1016/j.peptides.2013.12.011.

120. Zhang, X., He, C., Wang, M., Zhou, Q., Yang, D., Zhu, Y., Feng, W., Zhang, H., Dai, A., Chu, X., Wang, J., Yang, Z., Jiang, Y., Sensfuss, U., Tan, Q., Han, S., Reedtz-Runge, S., Xu, H.E., Zhao, S., Wang, M.W., Wu, B., Zhao, Q., 2021. Structures of the human cholecystokinin receptors bound to agonists and antagonists. Nat Chem Biol 17(12), 1230–1237. 10.1038/s41589-021-00866-8.

121. Zhu, Z., Tsuchimoto, M., Nagata, S., 2022. CCHamide-2 Signaling Regulates Food Intake and Metabolism in Gryllus bimaculatus. Insects 13(4). 10.3390/insects13040324.

