## Supplemental Materials for "Functional characterization of *Ixodes* neuropeptide receptors"

#### Synthetic Peptides Used for a calcium release (FLIPR) fluorescence assay

| Peptide | Sequence | Note |
|---|---|---|
| Isca sNPF | $NH_2$-GGRCPCLRLRF-amide | |
| Isca sNPF Scrambled | $NH_2$-RPLGRFGSLRS-amide | |
| Isca sulfakinin | $NH_2$-SDDY($SO_3H$)GHMRF-amide | Tyr4 sulfation |
| Isca sulfakinin – $SO_3H$ | $NH_2$-SDDYGHMRF-amide | No tyrosine sulfation |
| Isca CCHa | $NH_2$-SCKMYGHSCLGGH-amide | Disulfide Cys2-Cys9 |
| Isca CCHKRa | $NH_2$-SCKMYGHSCLGGHKR-amide | Disulfide Cys2-Cys9 |
| Isca CCHa Scrambled | $NH_2$-KSHCGLGHYSGCM-amide | Disulfide Cys4-Cys12 |

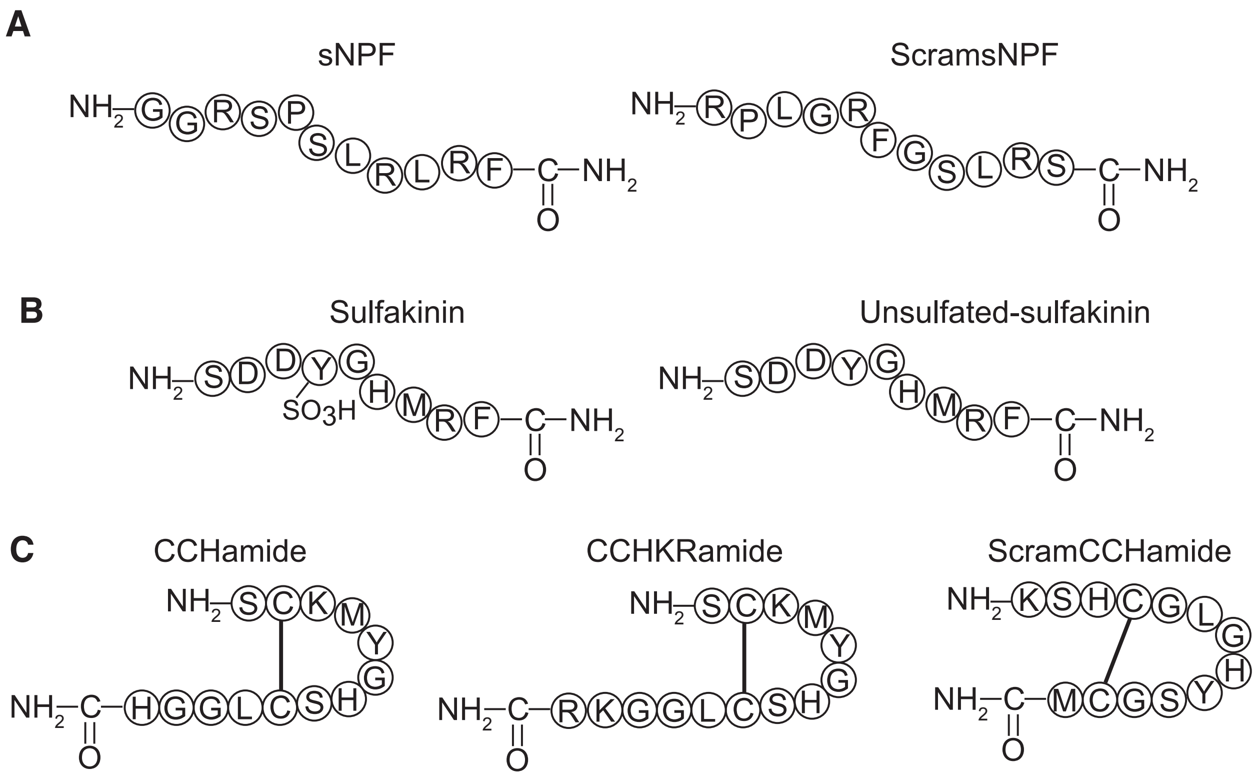

#### Primers used for RT-PCR

| Name | Target | Accession No. | Sequence | Location |
|---|---|---|---|---|
| **Primers used for qRT-PCR** | | | | |
| sNPFR-2F | sNPFR | KC439540 | gaaaggagttcaagctggttcta | 1037-1059 |
| sNPFR-1F | sNPFR | KC439540 | caagtctcgtagtgacctgttc | 1179-1200 |
| sNPFR-2R | sNPFR | KC439540 | tcaaggtgcgttccagttt | 1132-1114 |
| sNPFR-1R | sNPFR | KC43940 | ggtgtcaatgggtagctcaa | 1290-1271 |
| CCR-2F | CCHaR | XM_002415437 | gatgtggtactactttgctcca | 1020-1041 |
| CCR-1F | CCHaR | XM_002415437 | tgtggcattgtacctggttag | 1128-1148 |
| CCR-2R | CCHaR | XM_002415437 | caggagttgacgaaggtcatta | 1118-1097 |
| CCR-1R | CCHaR | XM_002415437 | ggtcctgaaggaatacgagttg | 1239-1218 |
| SKR-1F | SKR | XP_029826695 | caaccagaagcggtctactatg | 907-928 |
| SKR-2F | SKR | XP_029826695 | gtttcgcgtcagcttcct | 1020-1037 |
| SKR-1R | SKR | XP_029826695 | ttcatgaagcagtaggtgatgg | 1013-992 |
| SKR-2R | SKR | XP_029826695 | ctgtcctaccacgtacacaaac | 1128-1107 |
| RPS4-F | RPS4 | DQ066214 | ggtgaagaagattgtcaagcagag | 180-203 |
| RPS4-R | RPS4 | DQ066214 | tgaagccagcagggtagtg | 259-241 |
| psNPF-F | ppsNPF | XM_029980760 | ctgcaccgcttcctgac | 348-364 |
| psNPF-R | ppsNPF | XM_029980760 | tggtccgtgtaggccttag | 428-410 |
| pSKA-1F | ppSK | XP_029979447 | catacttgccaatggcttcg | 231-250 |
| pSKA-1R | ppSK | XP_029979447 | gaacctcatgtgtccgtagtc | 321-301 |
| pCCa-1F | ppCCHa | XM_029978301 | gatgcagatgctgaggaaatca | 305-326 |
| pCCa-1R | ppCCHa | XM_029978301 | ggttgcggagaatctgaagaag | 420-399 |
| **Primers used for RT-PCR** | | | | |
| sNPF-F1 | sNPF | XM_029980760 | atggtttcgctccggagacgg | 105-120 |
| sNPF-R1 | sNPF | XM_029980760 | cctcgagcctcataatggtttcgct | 91-115 |
| sNPF-F2 | sNPF | XM_029980760 | ttaggcggccggtgcgga | 396-413 |
| sNPF-R2 | sNPF | XM_029980760 | ctccgcgctgcccgttgc | 438-455 |
| CCHa-F1 | CCHa | XM_029978301 | atgtgcttcaaacctgcgatgc | 87-108 |
| CCHa-R1 | CCHa | XM_029978301 | gttcactgaagtcggttgcggag | 411-433 |
| SK-F1 | SK | XP_029979447 | caggaaccacgaagacgctctcat | 31-54 |
| SK-R1 | SK | XP_029979447 | tcacttgcggccgaacctcat | 313-333 |
| SK-R2 | SK | XP_029979447 | cgtgcgctcgcctccggt | 338-355 |
| H2B-F | H2B | XM_002402577 | cgcacgtccgacaagaagaagaaa | 128-151 |
| H2B-R | H2B | XM_002402577 | gaagaggtgtacttcgtgaccg | 414-435 |

#### DNA sequences used to express *Ixodes scapularis* GPCRs

>CCHaR-1D4 cDNA sequence of *Ixodes scapularis* CCHamide receptor used for expression in HEK293T cells

```
ATGAGTTCTCTTGTCGTCAACGCTACCCTAGCCACCCTCCGGGCAAACAGAGATCCATCTGCTT
CCCGAGCAGGCATCAATGCGTCACAGAATTCGCATGTCAACCTCACCACGGGCTATGGCGACTC
TGAAGAGTTCGTCCCATATGAGCAACGGCTGGAGACGTATGTTGTACCAACGCTGTTCGCATTG
ATATTTATCGTCGGGCTCCTGGGAAACGGAACGCTTATCCTCGTATTTATCAGAAACAGGACAA
TGCGAAGCGTTCCCAACATCTACATAATGAGCTTATCAATTGGCGATTTTATAGTAATCGCTGG
AACGGTGCCTTTTATCAGTACCATCTATGTGCTGGATTCGTGGCCCTACGGACTGTTTCTTTGT
AAACTCAGCGAGTTTCTCCGAGATGTCTCCATTGGTGTGACTGTACTGACGCTTACGGTCCTCA
GCATTGATAGGTACGTGGCCATTGCCATGCCACTGCTTAATCACAAAGGAAGACGACATACTAG
AAGAACGGTCACAATCCTTCTAGCCATTTCTGTGTGGATTGTGGCCATTTTAATGGCGATTCCA
GGGACCCACTATTCGTTTGTCATGCAAGTGCAGGCGACGCCCAACTTGCATTACAGTGTTTGCT
ACCCCTTCCCGCCGGAAATGTGGCCCTGGTATCCCAAGCTCATGGTCCTTCTGAAGTTCCTCAT
ACAGTACGCTATTCCACTGGTCATCATTGGCACCTTCTATTGCCTGATGGCCCGACAGCTTATT
CGGACATCGAGGGCTCACCTGTCGCAGACAAGCTGCGGTGGCGTTGCACATCTGAAGCAGATGA
AGGCTCGAGTGAAGGTCGCAAAGATTGCCCTTGCGTTCGTGCTTCTCTTCGCAGTCTGCTTTTT
TCCCAACCACGTATTCATGATGTGGTACTACTTCGCTCCAAACGCGCCGTCCCAGTACAACAGC
TTCTGGCACGTGTGGAAAATCATGGGATACGTAATGACCTTCGTCAACTCCTGCTTGAACCCTG
TGGCATTGTACCTGGTTAGCGGTGTGTTCAGAAACCATTTCAAGCACTATCTTTTTTGTGGGCG
GCATCCCAATGTGCACGCAACAGAGACCAGCCAAGTGGCGCCTGCCTAA
```

>SKR-1D4 cDNA sequence of humanized *Ixodes scapularis* sulfakinin receptor for expression in HEK293T cells

```
ATGGAAACTTGTAATCTCTCAACGGAAGGGAACGGTTCTGATGCGCGCCCCGCATATTCTTGGT
GGAGGTCAGACCAGGCCGTATTGGTTGCGCCATACACTGTTATTTTGCTTCTTGCCGTCTTGGG
AAACGGACTGGTGATTGTCACCCTTGCCGTCAACAAAAGAATGCGAACTGTTACCAATCTGTTC
CTTCTTAATTTGGCCGTCAGCGACTTGTTGCTGGGAGTTTTTTGTATGCCGTTTACTTTGGCAG
GGGTGCTTCTGCGCGAATTTGTGTTCGGCGAATTGATGTGTAGGTTGATTCCTTATCTTCAAGC
AGTAAGCGTCTGCGTATCCGCATGGACACTGATGGCAATGTCAGTTGAACGGTATTTCGCTATT
TGTTATCCGTTGAGGTCCCGCACATGGCAAACTCTTAGACACGCGCAAAGGACCATTGCAGCTG
TTTGGGTAGCTTCCTTGCTGCTTATGCTCCCAATCGCCCTCTTGTCTCAACTCCAGCCGGTCAA
GGACAGCGGTAAGATGAAATGTAGGGAAGATTGGCGAGAACCGCTTTACGAGCGGTTGTTCACA
CTCTTCTTGGATGCCCTCCTCCTCGTTCTCCCACTGTTGGGCATGACTGCGACATACTCCCGAA
TAGCAGCTACCCTCCGGTCAGCTATGCACCACGCCCCGAGCGACGTGGCTTGGCACAACGGTAG
CGCATTCAGTGCCGTACATCCGGCCGACCAGCAGCCTGCATCTAGGTTGTCACATCAACCGAGC
ACGATCCGATGGCACCAACGGGACCAGGATCGAAGCCTTGCGACCAAGCAGCGCATAATAAGGA
TGCTTTTTGCTGTTGTTGTTGAATTTTTTGTATGTTGGACGCCCCTTTACGTTCTGAACACTGT
GTCTCTGTTCCAACCAGAGGCTGTCTACTATGGACTTGGGTACCGAGGAATTTCCTTTTTGCAA
CTGCTGGCCTATGCCAGTTCCTGTTGCAATCCTATTACCTACTGTTTTATGAACAGGACATTCC
GGGTATCTTTTCTGGGTCTGTGTCGACATTGCTGTCGGACAAAGGGCAAACTCGGAACCTCACG
ACACAGTACAAAACAAAGCTTTGTATACGTCGTCGGGCAAGAGGCACAGACCGAGACCAGTCAG
GTTGCGCCAGCATGA
```

> KC439540 cDNA sequence of *Ixodes scapularis* sNPF receptor (NPYLR1A) for expression in HEK293T cells

ATGTCAGTGCCTGTTGAAGCTGGATTGCATATCAGGCGGACGTTCGAGCACGTCCCCAGGCTAC
GCCAGATGAACGCAACCTTTCTCGTGCTCAATGGCTCGGCCAGTGGACTCAACTACAGCAGCGT
GGTGGACATGGATGTCTTCTCCGGCTATGATTACATGTACATCACAAGCATCCCTGCTGTGCGC
GCCTTCTTCTACTGCATCTACGTGCTCATTTTTGTGACCGGAATATGTGGCAACATCCTCGTCT
GCTTTGTCGTCTTCCACAAAAACTCCATGCAAACCGTCACCAACATTTTTATCGCCAATCTGGC
ACTCTCGGACATCCTGCTGTGTGCGCTTGCGGTGCCCTTCACGCCTCTGTACCACTTCATGACG
ACATGGGCCTTCGGAAGTCTGCTGTGTCACCTTGTCCCCTACGCGCAGGGTGTTAGCGTATACA
TCTCCGCCTTCACGCTCATGGCCATCGCCATCGACAGATTTTTTGTGGTCATCTACCCCTTCCG
ACCTCGAATGCGGCTCTCAGTTTGCTTCACAATCATCGTCAGCGTGTGGGTGAGCAGTGCCTTG
CTGACGTTACCCTACGGTATCTTCATGGGCCTCATCCAGGATCCACAGAACGGGAAACGATACT
ACTGTGAGGAGGAGTGGCCCTCCGAGGTAAACCGGAAGACCTTCAGCTCGCTCACCACCACGCT
TCAGTTCTTGGTGCCGTTGAGCATTGTCACCTTTTGCTACGTGCGCGTTTGCTGTCGCTTGCAA
GACCGGGTGCGCGCTAAGCCAGGGGCACGCTCACTCAAGGAAATGGAGCGGAAGCGGACGAGGC
GGACAAACCGCATGCTCATTTCCATGGTGCTCATCTTCGCTGTCTCGTGGATGCCTCTCAACTT
GTACAATCTGGTGGCTGACTTCTACATTCCGGCCTCCAAGTGGCCCTACTCCAATGCCTTTTTC
TTTCTGTCGCACGCGATTGCCATGAGCTCAACGTGTTACAATCCTTTCTTGTACGCCTGGCTCA
ATGAAAACTTTCGAAAGGAGTTCAAGCTGGTTCTACCGGGGTTCATTTCGCCACATCTTGCCGC
AGAAAGGGCGGCCGACAACGCCACCAAACTGGAACGCACCTTGAACGGTCGTGACACGAACACC
GTTCAAGAAACAATAACGACTCGGAACAAGTCTCGTAGTGACCTGTTCAGTGAACGTAGATCCA
GCCAAGCTACGTGTGCTCGCTACGTTGCATCGTCTGGTCCAGAGGTCGTCCTCCTTGAGCTACC
CATTGACACCGAGAGCCACGCGGTACTCCTCACCGCCATATAA

>KC439541 cDNA sequence of *Ixodes scapularis* NPYLR1B for expression in HEK293T cells
ATGAACGTACTTCTGCAGTTTCCGCTGGCGTCCAGCTTGGAGAATCTGAGCCTGGAGGCAGAAA
ATACGAGCAGCCAATTTCACAACGTGACAAGTTATCACGTCTCAGTAATCAATGATTACATAAC
GAGCATTCCCGCCGTGAAAGCTTTCTTCTACTGCATATACATCTTGATTTTTGTCGTGGGCATT
TGCGGTAACGTGCTCGTGTGTTATGTCGTATTCCGAAACAAGCCCATGCAGACCGTGACCAACT
TCTTCATTACGAACTTGGGGCTCTCGGACATTCTTCTTTGCACGCTGGCCGTTCCTTTCACGCC
ACTATATCAGTTTATGCGCAAGTGGGTGTTTGGCCGAGTACTGTGCCACCTGGTCCCCTACGCG
CAGGGCGTCAGCGTCTACATCTCCTCCTTCACCCTTATGGCCATCGCCATCGACCGATTCTTCG
TGATCATTTACCCGTTCAAGCCGCGGCTCCAAATAAAAGTATGTTTCATGATCATTGTTTGCAT
CTGGTTGACCGGAGCCTTGCTGACTCTGCCCTACGGCATCTTTATGCACCTTACTCCGGATCCA
GATGACGGTCGACGACACTACTGCGAAGAGAAGTGGCCAGGCGAAGAGAGTCGTCGCACCTTCA
GCTTTTCGACAAGCACTCTGCAGTTTGTTGTTCCTTTCGGTATAATCAGTTTTTGCTACATGCG
CGTCTGCTGCAAGCTTCGCGACCGCGCACGTGCCAAGCCTGGCGCCAAGTCGGTCAAGAAAGAG
GAGCTTGAGCGCAAACGGACTAGACGCACCAATCGCATGCTCATCTCGATGGTGGTCATCTTCG
GCGCCTCCTGGCTACCCCTCAACTTATACAATCTCGTCATGGACTTCTTTATTCAAGCTGCAAG
CTGGAAATACGCCAACGCGTTCTTCTTTCTCTCCCATGCCGTCGCCATGAGCTCGACCTGTTAC
AACCCTTTCTTATACACCTGGCTCAACGAGAACTTCCGCAAGGAGTTCAAGCTGGTGCTGCCCT
ACTTTAGCGCTAGTGCACCCGCTCGTCGCACCAACGGCTCCAAGCCAGACCGCACCTGTTGCAA
CGGGCGCGAAGAGGTCCAGGAGTCGTTCGTGCCTCTCGCCTCCCAGCCGCCATCGCAGCAGCCG
GCCACCGACGACACTTCAGCATCTCGCACCACGCCCCCAGCTCCTGCTGTTCACTACGTATGCG
AGACGGACACCGTCAGGTTACACGTGTCCAACGACTCCAAGGAGGAGCTGTGA

**Prepro-sNPF sequences used in alignments**

> XP_042142062.1 uncharacterized protein LOC8032040 [Ixodes scapularis]
MVSLRRRPASFSIACVFVVLLVAETFVAAYADFNGERDMRDLVELLLKNEQESQLSHTMERKGG
RSPSLRLRFGRRSDPAWSDTLHRFLTAGNAGGDSGHSAPAA

>KAG0414334.1 hypothetical protein HPB47_008506 [Ixodes persulcatus]
MVSLRRRPASFSIACVFVVLLVAETFVAAYADFNDMRDLVELLLKNEQESQLSHTMERKGGRSP
SLRLRFGRRSDPAWSDTLHRFLAAGNTAGDSGHSAPAA

>QYF10816.1 sNPF [Rhipicephalus microplus]
MPSPAITRCLVLLLLVQAALAFPDYKDIRDLYELMGKGEQEGSGHAKERKAGYTPSLRLRFGRR
SDPAWSEARIWDAQRTV

>XP_037502403.1 short neuropeptide F-like [Rhipicephalus sanguineus]
MPSPAATRCLVLLLLLQAALAFPDYKDLRDLYELMAKGEQESSGHAMERKAGYTPSLRLRFGRR
SDPAWNEARTWDGQRTA

>XP_037572565.1 short neuropeptide F-like [Dermacentor silvarum]
MPSPAAARCLVLLLLLQAALAFPDYKDLRDLYELMAKGEQESAGHAMERKAGYTPTLRLRFGRR
SDPAWNEARTWDAQRAA

>KAH6931153.1 hypothetical protein HPB50_022510 [Hyalomma asiaticum]
MPSPAATRCLVLLLLLQAALAFPDYKDLRDLYELMAKGEQESSGHAMERKAGYTPSLRLRYGRR
SDPAWSEARTWDAQRAA

>KAI2810540.1 hypothetical protein BLOT_001703 [Blomia tropicalis]
MSKSKTNIRTATLSTAFLVLVICQIVTSAPSMAYDYDNIRDLYEMLLRQESPNQFVHQMERKGG
RGPSLRLRFGRRSDPLWSKLSIPNGNGIAMINGIGGGGGNSMNDDEKVPYKK

>XP_027199553.1 uncharacterized protein LOC113793690 [Dermatophagoides pteronyssinus]
MKISKSQQQQQLNHQQPSSTMISSIMSYSSMMNYYPSSSSISSSNKTKNSSLISRTTTTNRTTQ
MAMAFLLLLLICAQIVTSAPAYDLEKLRDLYEIWLRQDSPTPFVHQMERKGGRSPTLRLRFGRRS
DPLWNDKMPSIKNNRDNINSIEDTDLERMPWKE

>KAF7489901.1 hypothetical protein SSS_2534 [Sarcoptes scabiei]
MLYSNQISIDNQSEEDKIFHKNQIYGKKFSNSFADLNRTEEEISKSNSIKMSSKTSPLISSSQS
QPSSSQFSSLSSSSSSSSSLLLSSSSSSIASNQSSSKFNCLKISSHSSQIVFVFLLLIITTQFV
SSAPAYDYENLRDLYEILLRQDNPSPFVHQMERKGGRSPTLRLRFGRRSDPLWTQTKLAEIQSN
PGASNDHLERNVGGGVAFDPERVEGSILNDERTIEKVPFRK

>GFY79546.1 hypothetical protein TNIN_201361 [Trichonephila inaurata madagascariensis]
MSSGNAIRVCSFLLVVALLTADMISAAPYNDYDNLRDLYELLIRNEAAAAAPASAYNHQMERKG
GRSPSLRLRFGRRADPLWHAENPSDAPSN

>GFW53720.1 hypothetical protein TNCV_3938141 [Trichonephila clavipes]
MQFCFADGALQQTYYDVHPYSGSVFFPDLRDLYELLIRNEAAAAAPASAYNHQMERKGGRSPSL
RLRFGRRADPLWHAENPSDAPSN

>GBM09770.1 hypothetical protein AVEN_101811-1 [Araneus ventricosus]
MHEGRSEKGALQQTYYDVHPYPGSVFFPDLRDLYELLIRNEAAAAAPASAYNHQMERKGGRSPS
LRLRFGRRADPLWHADNPSDSTSN

>XP_008198705.1 PREDICTED: short neuropeptide F [Tribolium castaneum]
MQRYSAMKCLCAVTCIMIVVATVTSAAPSYADYDNNIRDLWEILLQKEAMDDKFAPGGPHQMVR
KSGRSPSLRLRFGRRSDASMTPEAAFMMAQAVDHETN

>NP_724239.1 short neuropeptide F precursor [Drosophila melanogaster]
MFHLKRELSQGCALALICLVSLQMQQPAQAEVSSAQGTPLSNLYDNLLQREYAGPVVFPNHQVE
RKAQRSPSLRLRFGRSDPDMLNSIVEKRWFGDVNQKPIRSPSLRLRFGRRDPSLPQMRRTAYDD
LLERELTLNSQQQQQQLGTEPDSDLGADYDGLYERVVRKPQRLRWGRSVPQFEANNADNEQIER
SQWYNSLLNSDKMRRMLVALQQQYEIPENVASYANDEDTDTDLNNDTSEFQREVRKPMRLRWGR
STGKAPSEQKHTPEETSSIPPKTQN

#### Protein sequences of sNPF receptors (sNPFR)

>AGX85008 NPYLR1A Ixodes scapularis
MSVPVEAGLHIRRTFEHVPRLRQMNATFLVLNGSASGLNYSSVVDMDVFSGYDYMYITSIPAVRAFFYCIYVL
IFVTGICGNILVCFVVFHKNSMQTVTNIFIANLALSDILLCALAVPFTPLYHFMTTWAFGSLLCHLVPYAQGV
SVYISAFTLMAIAIDRFFVVIYPFRPRMRLSVCFTIIVSVWVSSALLTLPYGIFMGLIQDPQNGKRYYCEEEW
PSEVNRKTFSSLTTTLQFLVPLSIVTFCYVRVCCRLQDRVRAKPGARSLKEMERKRTRRTNRMLISMVLIFAV
SWMPLNLYNLVADFYIPASKWPYSNAFFFLSHAIAMSSTCYNPFLYAWLNENFRKEFKLVLPGFISPHLAAER
AADNATKLERTLNGRDTNTVQETITTRNKSRSDLFSERRSSQATCARYVASSGPEVVLLELPIDTESHAVLLT
AI

>CAL6237959 NPYLR1 Ixodes pacificus
MQTVTNFFIANLALSDILLCALAVPFTPLYHFMTTWAFGSLLCHLVPYAQGVSVYISAFTLMAIAIDRFFVVI
YPFRPRMRLSVCFTIIVSVWVTSALLTLPYGIFMGLIRDPQNGTRYYCEEEWPSEVNRKTFSSLTTTLQFLVP
LSIVTFCYVRVCCRLQDRVRAKPGARSLKEMERKRTRRTNRMLISMVLIFAVSWMPLNLYNLVADFYIPASKW
PYSNAFFFLSHAIAMSSTCYNPFLYAWLNENFRKEFKLVLPGFISPHLAAERAADNATKLERTLNGRDTNTVQ
ETITTRNKSQVP

>KAG0445100 Ixodes persulcatus
MSVPVEAGLHIRRTFEHVPRLRQMNANGTTFLVLNGSASGLNYSSVVDMDVFTGHDYMYITSIPAVRAFFYCI
YVLIFVTGICGNILVCFVVFNKNSMQTVTNFFIANLALSDILLCALAVPFTPLYHFMTTWAFGSLLCHLVPYA
QGVSVYISAFTLMAIAIDRFFVVIYPFRPRMRLSVCFTIIVSVWVTSALLTLPYGIFMGLIEDPQNGKRYYCE
EEWPSEVNRKTFSSLTTTLQFLVPLSIVTFCYVRVCCRLQDRVRAKPGARSLKEMERKRTRRTNRMLISMVLI
FAVSWMPLNLYNLVADFYIPASKWPYSNAFFFLSHAIAMSSTCYNPFLYAWLNENFRKEFKLVLPGFILPHLA
AERAADNATKLERTLNGRDTNTVQETITTRNKSRSDLFSERRSSQATCARYVASSNPEVVLLELPIDTESHEV
LLTAI

>CAL6195172 Ixodes hexagonus
ENTTRQLHNVTTYHVSVINDYITSIPAVKAFFYCIYILIFVVGICGNVLVCYVVFRNKSMQTVTNFFITNLGL
SDILLCTLAVPFTPLYQFMRKWVFGRVLCHLVPYAQGVSVYISSFTLMAIAIDRFFVIIYPFKPRLQIKVCFM
IIISIWLTGALLTLPYGIFMHLTPDPDDGRRHYCEEKWPDEESRRTFSFSTSTLQFVVPFGIISFCYMRVCCK
LRDRARAKPGAKSVKKEELERKRTRRTNRMLISMVVIFGASWLPLNLYNLVMDFFIQAASWKYANAFFFLSHA
VAMSSTCYNPFLYTWLNENFRKEFKLVLPCF

>AGX85009 NPYLR1B Ixodes scapularis
MNVLLQFPLASSLENLSLEAENTSSQFHNVTSYHVSVINDYITSIPAVKAFFYCIYILIFVVGICGNVLVCYV
VFRNKPMQTVTNFFITNLGLSDILLCTLAVPFTPLYQFMRKWVFGRVLCHLVPYAQGVSVYISSFTLMAIAID
RFFVIIYPFKPRLQIKVCFMIIVCIWLTGALLTLPYGIFMHLTPDPDDGRRHYCEEKWPGEESRRTFSFSTST
LQFVVPFGIISFCYMRVCCKLRDRARAKPGAKSVKKEELERKRTRRTNRMLISMVVIFGASWLPLNLYNLVMD
FFIQAASWKYANAFFFLSHAVAMSSTCYNPFLYTWLNENFRKEFKLVLPYFSASAPARRTNGSKPDRTCCNGR
EEVQESFVPLASQPPSQQPATDDTSASRTTPPAPAVHYVCETDTVRLHVSNDSKEEL

>KAG0445099 Ixodes persulcatus
MALPVVPGLNMNVLLQFPLASSLENLSLETENTSRQFHNVTTYHVSVINDYITSIPAVKAFFYCIYILIFVVG
ICGNVLVCYVVFRNKSMQTVTNFFITNLGLSDILLCTLAVPFTPLYQFMRKWVFGRVLCHLVPYAQGVSVYIS
SFTLMAIAIDRFFVIIYPFKPRLQIKVCFMIIVCIWLTGALLTLPYGIFMHLTPDPDDGRRHYCEEKWPGEES
RRTFSFSTSTLQFVVPFGIISFCYMRVCCKLRDRARAKPGAKSVKKEELERKRTRRTNRMLISMVVIFGASWL
PLNLYNLVMDFFIQAASWKYANAFFFLSHAVAMSSTCYNPFLYTWLNENFRKEFKLVLPCFSASAPARRTNSS
KPDRTCCNGREEVQESFVPHASQPPSQQPATDDTSASRTTPPAPAVHYVCETDTVRLHVSNDSKEEL

>XP_040075532 RYaR1 Ixodes scapularis
MDSTNGPSAPPTATSNWTSQPASTESAACDLPPPVPEGMQALMYIMYIAVSVAAIGGNGIVCYIVLAYQRMRT
VTNMFIMNLAIGDILMASLCIPFTFVSNLLLGYWPFGGVMCVVVTYAQCVTVFISAYTLIAISVDRYTAIVYP
LRPRMTKLRSKIIIGVVWLVALVTPLPTALVTQLVPHPCANQTYYCLEQWGTPEQTTYYSMALMILQYFFPLL

ALIFTYTRIAVVVWGKETPGEAQDERDQRMAASKRKMIKMMIMVVTVFMLCWFPLNAYILLSDLNPDINSYEY
IRYIYFVIHWLAMSHASYNPFIYCWMNAKFREGFGNLTRRCWPPVCWPGRLRRQTLRKESNEGAALRRVNTYT
TYVSVRAAGGGSSLKFNGNRGLKEVNGKIGDYEDSRV

>XP_037568438 Dermacentor silvarum
MGNVSHPVDYVSGVPAVRAFFYCVYGLVFAVGVSGNALVCFVVARQRAMHTVTNLFIANLALSDILLCALAVP
FTPLHQFVGAWPLGAALCRLVPYAQGVSVYVSSFTLTAIAVDRFFVIMHPFRGRLRLPVCGALIGLVWLAGAL
LTLPYGLFIGLTADGGAFCEERWPSEHSRRAFSLCTSALQFGLPFAVISFCYMRVCCKLRERARAKPGAKSMQ
KEQLERRRTRRTNRMLVSMVAIFGACWLPLNLYNLAIDFSVRAASWQFANAFFFLAHAIAMSSTCYNPFLYTW
LNDSFRKEFKAVLPCFASRRAEPPAVRYVCSGDAVKL
>XP_037282410 Rhipicephalus microplus
MGNVSHHVDYVSSVPAVRAFFYCVYGLVFAVGVSGNALVCFVVARQRAMHTVTNLFIANLALSDILLCTLAVP
FTPLHQFVGAWPLGSALCRMVPYAQGVSVYVSSFTLTAIAVDRFFVIMHPFRGRLRLPVCGALIVLIWVAGAL
LTLPYGLFIGLTADGGSFCEERWPSEESRRAFSLCTSALQFGVPFTVISFCYMRVCCKLRERARAKPGAKSMQ
KEQLERRRTRRTNRMLVSMVVIFGACWLPLNLYNLAIDFSLRAASWQFANAFFFLAHAIAVSSTCYNPFLYTW
LNDSFRKEFKAVLPCFSSRRAELPPVRYVCSGDAVKL

>XP_037504759 Rhipicephalus sanguineus
MGNVSHHVDYVSGVPAVRAFFYCVYGLVFAVGVSGNALVCFVVARQRAMHTVTNLFIANLALSDILLCTLAVP
FTPLHQFVGAWPLGSALCRMVPYAQGVSVYVSSFTLTAIAVDRFFVIMHPFRGRLRLPVCGALIGLVWVAGAL
LTLPYGLFIGLTADGGAFCEERWPSERSRRAFSLCTSALQFGVPFTVISFCYMRVCCKLRERARAKPGAKSMQ
KEQLERRRTRRTNRMLVSMVAIFGACWLPLNLYNLAIDFSVRAASWQFANAFFFLAHAIAMSSTCYNPFLYTW
LNDSFRKEFKAVLPCFWSRRAELPPVRYVCSGDAVKL

>XP_037273498 Rhipicephalus microplus
MAGSKNMPNGCLEPGYCDLFLCIILYFALVMLAAFLAAYYSLPINDTSSYGGGRSGAAAALFGLLYAVVFALG
VSGNALVCVAVLRRHTMRTVTNLLVANLALSDILLCALAVPFTPLYLFLGRWPFGAALCHLVPYAQSVSVYVS
SLTLTAIAVHRFRAVVHPLRPRLLRPGSCGILCVALWVASALLTLPYAAFVRLVRHGRTAYCEEVWPSLRGRQ
LFGALTTAAQFVLPLGVVGGCYVRVGQRLRGRRRRSAHRTQLMLLAMVLAFALAWLPLNACNLLADFHSASLG
WPMGDVLFLAAHAAAMSSTCYDPLLYAWFNDNFRKQFVRMLHPDPQQNTVLETLRSDDPRCKVSEHQHSLSSQ
ALPDPEMEESRGIAATFV

>XP_037525186 Rhipicephalus sanguineus
MDAQSINDTASYGGGRSGAAAALFGLLYAVVFALGVSGNVLVCVAVLRRHTMRTVTNLLVANLALSDILLCAL
AVPFTPLYLFLGQWPFGTALCHLVPYAQGVSVYVSSLTLTAIAVHRFRAVVHPLRPRLLRPGSCGILCVALWV
ASALLTLPYAAFVRLVRHGRAAYCEEVWPSARGRQLFGALTTAAQFVLPLGVVGGCYVRVGRRLRGRRRRSAH
RTQLMLLAMVLAFALAWLPLNACNLLADFHSASLGWPMGDVLFLAAHAAAMSSTCYDPLLYAWFNDNFRRQFV
RMLHPDQQQNTVLETLRSDDPRCKVSEHHRCPSSQELHPEMEGSRGIAATFV

>Q9VW75 sNPFR DROME
MANLSWLSTITTTSSSISTSQLPLVSTTNWSLTSPGTTSAILADVAASDEDRSGGIIHNQFVQIFFYVLYATV
FVLGVFGNVLVCYVVLRNRAMQTVTNIFITNLALSDILLCVLAVPFTPLYTFMGRWAFGRSLCHLVSFAQGCS
IYISTLTLTSIAIDRYFVIIYPFHPRMKLSTCIGIIVSIWVIALLATVPYGMYMKMTNELVNGTQTGNETLVE
ATLMLNGSFVAQGSGFIEAPDSTSATQAYMQVMTAGSTGPEMPYVRVYCEENWPSEQYRKVFGAITTTLQFVL
PFFIISICYVWISVKLNQRARAKPGSKSSRREEADRDRKKRTNRMLIAMVAVFGLSWLPINVVNIFDDFDDKS
NEWRFYILFFFVAHSIAMSSTCYNPFLYAWLNENFRKEFKHVLPCFNPSNNNIINITRGYNRSDRNTCGPRLH
HGKGDGGMGGGSLDADDQDENGITQETCLPKEKLLIIPREPTYGNGTGAVSPILSGRGINAALVHGGDHQMHQ
LQPSHHQQVELTRRIRRRTDETDGDYLDSGDEQTVEVRFSETPFVSTDNTTGISILETSTSHCQDSDVMVELG
EAIGAGGGAELGRRIN

>NP_000900 NPY1R Homo sapiens
DDCHLPLAMIFTLALAYGAVIILGVSGNLALIIIILKQKEMRNVTNILIVNLSFSDLLVAIMCLPFTFVYTLM
DHWVFGEAMCKLNPFVQCVSITVSIFSLVLIAVERHQLIINPRGWRPNNRHAYVGIAVIWVLAVASSLPFLIY
QVMTDEPFQNVTLDAYKDKYVCFDQFPSDSHRLSYTTLLLVLQYFGPLCFIFICYFKIYIRLKRRNNMMDKMR

DNKYRSSETKRINIMLLSIVVAFAVCWLPLTIFNTVFDWNHQIIATCNHNLLFLLCHLTAMISTCVNPIFYGF
LNKNFQRDLQFFF

>NP_001362399 NPY2R Homo sapiens
IDSTKLIEVQVVLILAYCSIILLGVIGNSLVIHVVIKFKSMRTVTNFFIANLAVADLLVNTLCLPFTLTYTLM
GEWKMGPVLCHLVPYAQGLAVQVSTITLTVIALDRHRCIVYHLESKISKRISFLIIGLAWGISALLASPLAIF
REYSLIEIIPDFEIVACTEKWPGEEKSIYGTVYSLSSLLILYVLPLGIISFSYTRIWSKLKNHVSPGAANDHY
HQRRQKTTKMLVCVVVVFAVSWLPLHAFQLAVDIDSQVLDLKEYKLIFTVFHIIAMCSTFANPLLYGWMNSNY
RKAFLSAF

>P49683 PRLHR Homo sapiens
QSLQLVHQLKGLIVLLYSVVVVVGLVGNCLLVLVIARVRRLHNVTNFLIGNLALSDVLMCTACVPLTLAYAFE
PRGWVFGGGLCHLVFFLQPVTVYVSVFTLTTIAVDRYVVLVHPLRRRISLRLSAYAVLAIWALSAVLALPAAV
HTYHVELKPHDVRLCEEFWGSQERQRQLYAWGLLLVTYLLPLLVILLSYVRVSVKLRNRVVPGCVTQSQADWD
RARRRRTFCLLVVIVVVFAVCWLPLHVFNLLRDLDPHAIDPYAFGLVQLLCHWLAMSSACYNPFIYAWLHDSF
REELRKLL

### Ixodid sNPFR/NPYLR Identity Matrix

| Species | Accesion Number | | 1 | 2 | 3 | 4 | 5 | 6 | 7 | 8 | 9 | 10 | 11 | 12 | 13 | 14 | 15 | 16 |
|---|---|---|---|---|---|---|---|---|---|---|---|---|---|---|---|---|---|---|
| *H. sapiens* NPY1R | NP_000900 | 1 | 100 | 28.14 | 32.53 | 31.74 | 31.8 | 32.16 | 31.48 | 31.42 | 33.98 | 31.08 | 31.21 | 31.21 | 31.21 | 30.95 | 31.29 | 30.95 |
| *I. scapularis* RYaR1 | XP_040075532 | 2 | 28.14 | 100 | 34.01 | 32.88 | 29.23 | 31 | 30.13 | 30.56 | 34.81 | 30.59 | 29.43 | 33.77 | 29.43 | 31.97 | 31.03 | 31.97 |
| *H. sapiens* PRLHR | P49683 | 3 | 32.53 | 34.01 | 100 | 36.64 | 40.71 | 40.71 | 36.58 | 37.33 | 40.87 | 38.01 | 37.76 | 37.76 | 37.76 | 40.27 | 40.61 | 39.93 |
| *H. sapiens* NPY2R | NP_001362399 | 4 | 31.74 | 32.88 | 36.64 | 100 | 36.04 | 36.04 | 37.71 | 38.23 | 39.53 | 38.57 | 40.96 | 40.96 | 41.3 | 37.11 | 36.77 | 36.77 |
| *R. microplus* | XP_037273498 | 5 | 31.8 | 29.23 | 40.71 | 36.04 | 100 | 93.31 | 42.44 | 45.04 | 51.85 | 44.66 | 41.86 | 45.85 | 41.55 | 53.9 | 52.54 | 53.22 |
| *R. sanguineus* | XP_037525186 | 6 | 32.16 | 31 | 40.71 | 36.04 | 93.31 | 100 | 45.03 | 47.2 | 51.85 | 47.2 | 44.86 | 47.12 | 44.86 | 53.22 | 52.2 | 52.88 |
| *D. melanogaster* | Q9VW75 | 7 | 31.48 | 30.13 | 36.58 | 37.71 | 42.44 | 45.03 | 100 | 47.8 | 55.92 | 47.47 | 48.56 | 58.31 | 48.46 | 51.98 | 51.67 | 51.67 |
| *I. scapularis* sNPFR | AGX85008 | 8 | 31.42 | 30.56 | 37.33 | 38.23 | 45.04 | 47.2 | 47.8 | 100 | 97.7 | 97.95 | 60.64 | 71.65 | 60.62 | 60.86 | 61.16 | 61.16 |
| *I. pacificus* | CAL6237959 | 9 | 33.98 | 34.81 | 40.87 | 39.53 | 51.85 | 51.85 | 55.92 | 97.7 | 100 | 98.03 | 69.9 | 76.34 | 69.9 | 65.15 | 65.15 | 65.53 |
| *I. persulcatus* | KAG0445100 | 10 | 31.08 | 30.59 | 38.01 | 38.57 | 44.66 | 47.2 | 47.47 | 97.95 | 98.03 | 100 | 60.92 | 72.27 | 60.9 | 60.86 | 61.16 | 61.16 |
| *I. scapularis* NPYLR1B | AGX85009 | 11 | 31.21 | 29.43 | 37.76 | 40.96 | 41.86 | 44.86 | 48.56 | 60.64 | 69.9 | 60.92 | 100 | 97.21 | 98.34 | 67.78 | 69 | 67.48 |
| *I. hexagonus* | CAL6195172 | 12 | 31.21 | 33.77 | 37.76 | 40.96 | 45.85 | 47.12 | 58.31 | 71.65 | 76.34 | 72.27 | 97.21 | 100 | 98.45 | 69.81 | 70.78 | 69.81 |
| *I. persulcatus* | KAG0445099 | 13 | 31.21 | 29.43 | 37.76 | 41.3 | 41.55 | 44.86 | 48.46 | 60.62 | 69.9 | 60.9 | 98.34 | 98.45 | 100 | 68.09 | 69.3 | 67.78 |
| *D. silvarum* | XP_037568438 | 14 | 30.95 | 31.97 | 40.27 | 37.11 | 53.9 | 53.22 | 51.98 | 60.86 | 65.15 | 60.86 | 67.78 | 69.81 | 68.09 | 100 | 94.53 | 96.66 |
| *R. microplus* | XP_037282410 | 15 | 31.29 | 31.03 | 40.61 | 36.77 | 52.54 | 52.2 | 51.67 | 61.16 | 65.15 | 61.16 | 69 | 70.78 | 69.3 | 94.53 | 100 | 97.26 |
| *R. sanguineus* | XP_037504759 | 16 | 30.95 | 31.97 | 39.93 | 36.77 | 53.22 | 52.88 | 51.67 | 61.16 | 65.53 | 61.16 | 67.48 | 69.81 | 67.78 | 96.66 | 97.26 | 100 |

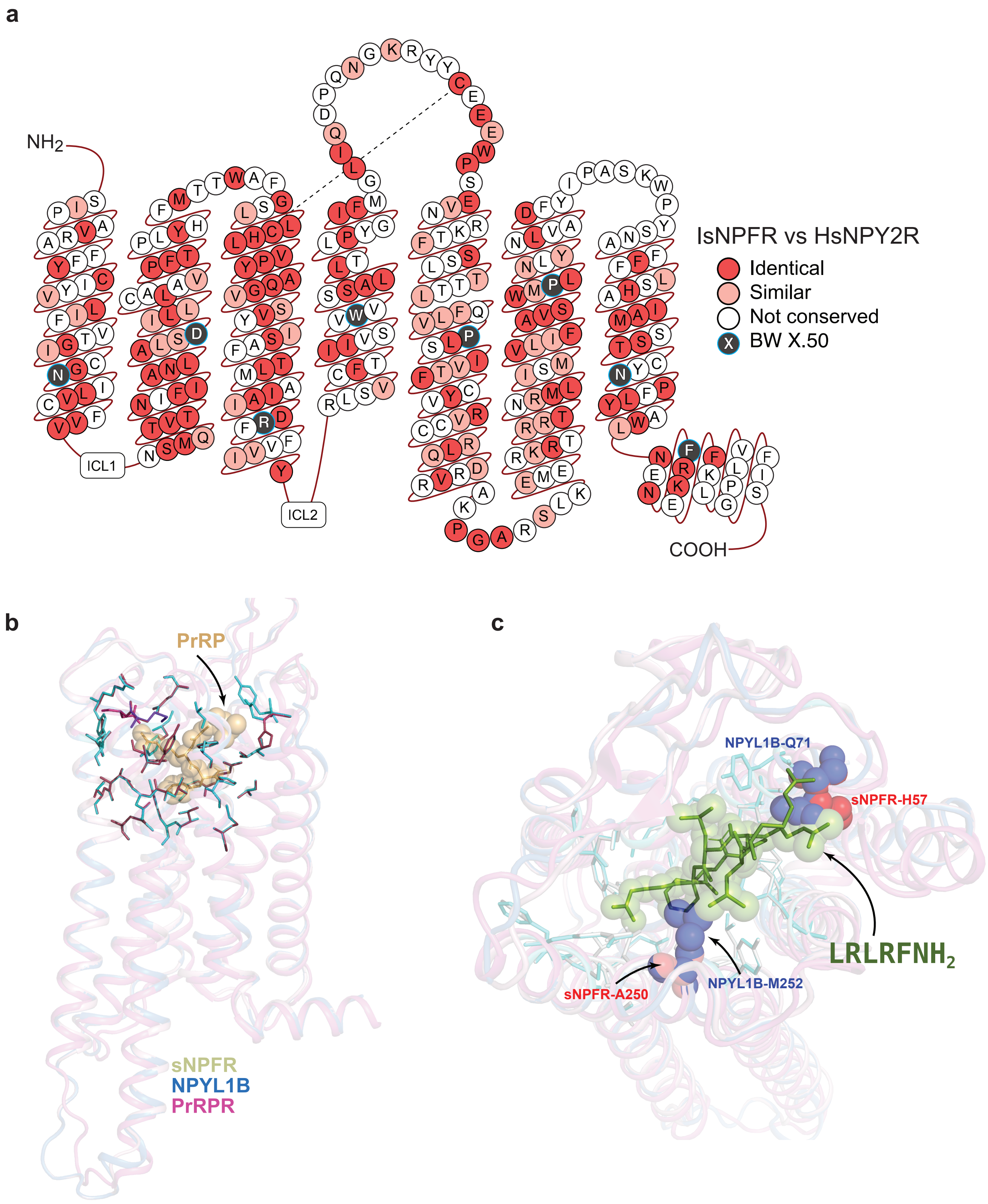
a
$NH_2$
ICL1
ICL2
COOH
IsNPFR vs HsNPY2R
Identical
Similar
Not conserved
BW X.50
b
PrRP
sNPFR
NPYL1B
PrRPR
c
NPYL1B-Q71
sNPFR-H57
$LRLRFNH_2$
sNPFR-A250
NPYL1B-M252

#### Alignment of *I. scapiularis* genomic sequences encoding prepro-SK with cDNA sequences from *I. scapularis* and *I. pacificus*

```
ppSK Isca PalLabHiFi 684  TTTCAAAAACCGTAGTAAAAAAAAAAA-----------AGTCCGGCTCATCGAGGAACGTTACAAATTCTCCCTTTCACATGTTGAATTAAGATATACATGTATGTAGGC
ppSK ISE6            670  TTTCAAAAACCGTAGTAAAAAAAAAAAAAAAAAAACGAGTTCGGCTCATCGAGGAACGTTACAAATTCTCCCTTTCACATGTTGAATTAAGATATACATGTATGTAGTT
ppSK Iperc genomic   653  CCTCAAGAACCGTAGTAAAAAAACAAA-----------AGTCGGGCTCATCGAGGAACGTTGCAAATTTTCCCTTTCACACGTTGAATTATGATATACATGTATGTAGGT

ppSK Isca PalLabHiFi 783  GCAGCGTAGATTAGAGTTATGTTGTTTTATCGAGAC--GCATACATTTCAAGCACCAAAAGCATCGTTCCTTCCTTCGACGACCAAATCTTGTAAGAACTACTTCCCGCA
ppSK ISE6            780  GCAGCTTAGATTAGATTTATATTGTTTTATCGAGACGGGCATACATTTCAAGCACCAAAAGCATCGTTCCTTCCTTCGACGACCAAATCTTGTAAGAACTACTTCCCGCA
ppSK Iperc genomic   752  GCAGCTTAGATTAGATTTACGTTATTTTATTGAGAC--CAATACATTTCGAGCACCAAAAGCATCGTTCCTTCCTTCGATGACCAAATCTTGTAAGAACTACTTCCCGCA

ppSK Isca PalLabHiFi 891  GCGCACGAATAATGCCCAAAGCTTCCTTTCTTCTT------------------------------TTTCCAAATAAACAGTTCCCCCTTTTCTTCCTCCGCGCAGGCCAA
ppSK ISE6            890  GCGCACGAATAATGCACAACGCTTCCTTTTTTCTT------------------------------TCTCCAAATAAACAGCTCCCCCTTTTCTTCCTCCGCGCAGGCCAA
ppSK Iperc genomic   860  GCGCACGAATAATGCTCAAAGCTTCCTCTTTTCTTGATGTGGCAAAGAGTATAGATGGCGTTACCTCTCCAAGTAAACAGCGCCGACTTTTCTTCCCCCACGCAGGCCAA

IsSK_RNA_PCR         1    -CGGGGGTAAAGAAGGCCATCAGCTAGCAACATGAGAGCGTCTTCGTGGTTCCTGCTTTGTCTTCTTGCAGCACTCGTTTACGGTTCGTGGTCGAGCCCAGCGTCCATGC
Ipac_ppSK            1    ------------ATGGCCG--ATCT-----CATATGACCGTAGTCGTCATCCCG--------------------CCGTTTACGGTTCGTGGTCGAGCCCAACGTCCATGC
ppSK Isca PalLabHiFi 971  GC-GGGGTACAGAAGGCCATCAGCTAGCAACATGAGAGCGTCTTCGTGGTTCCTGCTTTGTCTTCTGGCAGCACTCGTTTACGGTTCGTGGTCGAGCCCAGCGTCCATGC
ppSK ISE6            970  GCGGGGGTAAAGAAGGCCATCAGCTAGCAACATGAGAGCGTCTTCGTGGTTCCTGCTTTGTCTTCTTGCAGCACTCGTTTACGGTTCGTGGTCGAGCCCAGCGTCCATGC
ppSK Iperc genomic   970  GCGGGGGTACAGCAGGCCAGCAGCTAGCAACATGAGAGCGTCTTCGTGGTTCCTGATCTGCCTTCTTGCAGCACTCGTTTACGGTTCGTGGTCCAGCCCAACGTCCATGC

IsSK_RNA_PCR         110  AGCAGCGCCACCGCATGGCGATGGGCAAGTGGCTCAAGAGCGTCCTTCCCGGGGCGCCATCCGGCGGCGACGCGGGGAGTCGGAACAGCGGCGACATCGACACGGACATG
Ipac_ppSK            72   AGCAGCGCCACCGCATGGCGATGGGCAAGTGGCTCAAGAGCGTTCTTCCCGGGGCGCCATCCGGCGGCGACGCGGGGAGTCGGAACAGCGGCGACATCGACACGGACATG
ppSK Isca PalLabHiFi 1080 AGCAGCGCCACCGCATGGCGATGGGCAAGTGGCTCAAGAGCGTCCTCCCCGGGGCGCCATCTGGCGGCGACGCGGGGAGTCGGAACAGCGGCGACATCGACACGGACATG
ppSK ISE6            1080 AGCAGCGCCACCGCATGGCGATGGGCAAGTGGCTCAAGAGCGTCCTTCCCGGGGCGCCATCCGGCGGCGACGCGGGGAGTCGGAACAGCGGCGACATCGACACGGACATG
ppSK Iperc genomic   1080 AGCAGCGCCACCGCATGGCGATGGGCAAGTGGCTCAAGAGCGTCCTTCCCGGGGCGCCATCCGGCGGCGACGCGGGGAGTCGGAACAGCGGCGACATCGACACGGACATG

IsSK_RNA_PCR         220  ATCGATCCCGTCATACTTGCCAATGGCTTCGCCAAAAGGCAGGATGACGACTACGGTCATATGAGATTCGGCCGTTCCGACGACTACGGACACATGAGGTTCGGCCGCAA
Ipac_ppSK            182  ATCGATCCCGTCATACTTGCAAATGGCTTCGCCAAACGGCGGGATGACGACTACGGTCATATGAGATTCGGCCGTTCCGACGACTACGGACACATGAGGTTCGGCCGGAA
ppSK Isca PalLabHiFi 1190 ATCGATCCCGTCATACTTGCTAATGGCTTCGCCAAAAGGCAGGATGACGACTACGGTCATATGAGATTCGGCCGTTCCGACGACTACGGACACATGAGGTTCGGCCGCAA
ppSK ISE6            1190 ATCGATCCCGTCATACTTGCCAATGGCTTCGCCAAAAGGCAGGATGACGACTACGGTCATATGAGATTCGGCCGTTCCGACGACTACGGACACATGAGGTTCGGCCGCAA
ppSK Iperc genomic   1190 ATCGATCCCGTCATACTTGCCAATGGCTTCGCCAAACGGCAGGATGACGACTACGGTCATATGAGATTCGGCCGTTCCGACGACTACGGACACATGAGGTTCGGCCGGAA

IsSK_RNA_PCR         330  GTGAACCGACCGGAGGCGAGCGCACGTCGAAACGACGTGGTGACCGCCGTATACCTCCGCCACATGCGTTGGCGGACGAACTTCGACAAATAAAGATTTCTCCGGCATGG
Ipac_ppSK            292  GTGA----------------------------------------------------------------------------------------------------------
ppSK Isca PalLabHiFi 1300 GTGAACCGACGGGAGACGAGCGCACGTCGAAACGACTTTGTGACCGCCGTATACCTCCGCCACATGCGTTGGCGGACGAACTTCGAAAAATAAAGATTTCTCCGGCATGG
ppSK ISE6            1300 GTGAACCGACCGGAGGCGAGCGCACGTCGAAACGACGTGGTGACCGCCGTATACCTCCGCCACATGCGTTGGCGGACGAACTTCGACAAATAAAGATTTCTCCGGCATGG
ppSK Iperc genomic   1300 GTGAACCGACGGGAGACGAGCGCACGTCGAAACGACTTGGTGACCGCCGTATACCTCCGCCACGTGCGTTGGCGGACGAACTTCGACAAATAAAGATTTCTCCGGCGTGG

IsSK_RNA_PCR         440  TTCTGGTATATACCCTGCGTCTTAACTTTTTTTTT---------GCCAAGATTGGTTTACACAC------------------------------------------GCAT
ppSK Isca PalLabHiFi 1410 TTCTGGTATATACCCTGCGTCTTAATTTTTTTTTTTTTTTATTCGCCAAGATTGGTTTACACAC------------------------------------------GCAT
ppSK ISE6            1410 TTCTGGTATATACCCTGCGTCTTAACTTTTTTTTT---------GCCAAGATTGGTTTACACAC------------------------------------------GCAT
ppSK Iperc genomic   1410 TTCTGGTATATCCCCCGAGCTTGATTTTTCTTTTTGTGTTATTTTTTAAGACTGGTTTACACACTTCCGTTTGACTAAAACCGATTGCGCCAGAACCACTTTCTGAGCAT

IsSK_RNA_PCR         499  CTTGTGA-GGCTTTCTAGCTTTAAGGGTTCTGAGGCTGGAGCGTCGGATGCGCTCGAAAGATTGGCGGGAAAGGGAGGGGGGGGGGGGGGGGGGGGGAGTGTGTCCCCCC
ppSK Isca PalLabHiFi 1478 CTTGTGACGGCTTTCTACCTTAAGGGGTTCTGAGGCTGGAGCGTCGGATGTGCTCGAAAGATTGGCGGAAAA-------------GAAGGGGGGGAAAGTGTGTCCCCCC
ppSK ISE6            1469 CTTGTGA-GGCTTTCTAGCTTTAAGGGTTCTGAGGCTGGAGCGTCGGATGCGCTCGAAAGATTGGCGGGAAAGGGAGGGGGGGGGGGGGGGGGGGGGAGTGTGTCCCCCC
ppSK Iperc genomic   1520 CTTGTGACTGCTTTCTACCTTTAGGGGTTCTGAGGCAGAGGCGTCGGATGTGCTCGAAAGATTGGCGAGAAA-------GGGGGGGGGGGGGGGGTTAGTGTGTCCCCC-

IsSK_RNA_PCR         608  CCCCCCCCCCCCCCCCCCCCCGCCCCCTTCAACCTTTCTTTCTTCCGACCCATTCCCCTGCTCCGCGCCACTGCTTTGGGATAAACCCAATACAGAACAAGCAATGTGAA
ppSK Isca PalLabHiFi 1575 CCCCCCGCCCCCCCCCCCC---CCCCCCTCAACCTTTC-TTCTTCCGACCCATTCGCCTGCTCCGCGCCACTGCTTTGGGATAAACCCAATACAGAACAAGCAATGTGAA
ppSK ISE6            1578 CCCCCCCCCCCCCCCCCCCCCGCCCCCTTCAACCTTTCTTTCTTCCGACCCATTCCCCTGCTCCGCGCCACTGCTTTGGGATAAACCCAATACAGAACAAGCAATGTGAA
ppSK Iperc genomic   1621 ----------------------CCCCCCTCTATCTTTCTTTCTTCCGACCCATTCCCCTGCTCCGCGTCACTGCTTTGGAATAAACCCAATACAGAACAGACAATGGAAA

IsSK_RNA_PCR         718  CCTAACATTCTAAATGAACGTAGAACGATGTTTGAACAGCAGTAGAGGCTTTATTACACGACGTTTCGGGAAGAGACTAGAAGGGAACACTCTTTTCGAAACCTCGTGTA
ppSK Isca PalLabHiFi 1681 CCTAACATTCTAAATGAACGTAGAAAAATGTTTGAACAACAATAGAGGCTTTATAACACGACGTTCCGGGAAGAGACTAGAAG------TCTCTTTTTAAACCTCGTGTA
ppSK ISE6            1688 CCTAACATTCTAAATGAACGTAGAACGATGTTTGAACAGCAGTAGAGGCTTTATTACACGACGTTTCGGGAAGAGACTAGAAGGGAACACTCTTTTCGAAACCTCGTGTA
ppSK Iperc genomic   1710 CCTACCATTCTAAATG----------------------------------------------------------------------------------------------

IsSK_RNA_PCR         828  ATAAAGCCTCTACTGCTTTTCAAACATTTTTCTACGTTTACGACTTTTTTGTGAAGAATTGTTCATTAATACAGTTATTCTGAATGAAAAAAAAAAGTTTCCTCTCCTTG
ppSK Isca PalLabHiFi 1785 ATAAAGCCTCTACTGCTGTTCAAACATTTTTCTACGTTTACGACTTTCTGGTGAAGACTTGTTCATTAATACAGTTATTCTAAATGAAAAAAAAAAGTTTCCTCTCCTTG
ppSK ISE6            1798 ATAAAGCCTCTACTGCTTTTCAAACATTTTTCTACGTTTACGACTTTTTTGTGAAGAATTGTTCATTAATACAGTTATTCTGAATGAAAAAAAAAAGTTTCCTCTCCTTG
ppSK Iperc genomic   1725 ---------------------------------------------------------------------------------GGAAAAAAAAAACAAGTTTCCTCTCCTTG

IsSK_RNA_PCR         938  AACGCTGTCCTTTACGACGGAACCCTGGCGAT------------------------------------------------------------------------------
ppSK Isca PalLabHiFi 1895 AACGCTGTCCTTTACCACGGAACCCTGGCGATATGGTGGATTTCTTTTTCAAAAGTAACACAGAACAAACCTTGCCTTATTTGAAACTTGGAATCGTCAAACTATTTCAT
ppSK ISE6            1908 AACGCTGTCCTTTACGACGGAACCCTGGCGATATGGTGGATTTCTTTTTCAAAAGTAACACAGAACAAACCTTGCCTTATTTGAAACTTGGAATCGTCAAACTATTTCAT
ppSK Iperc genomic   1755 AACGCTGTCATTTACCACGGAACTCTGTCGATATGGTGGATTTATTTTTCAAAAGTAACACAGGA-AAACCTTGCCTCATTTGAAATTCGAAATTGTCAAACTATTTCAT
```

ppSK Isca PalLabHiFi : genome assembly from *I. scapularis* whole tick
ppSK ISE6 : genome assembly from *I. scapularis* ISE6 cell line
ppSK Iperc genomic : shotgun sequence from *I. persulcatatus* JABSTQ010000000.1
IsSK_RNA_PCR : RT-PCR product sequencing from adult *I. scapularis*(this work)
Ipac_ppSK : RT-PCR product sequencing from adult *I. pacificus* (this work)

**Sulfakinin peptide sequences used in alignments**

```
> Ixodes_scapularis sulfkinin (this work)
QDDDYGHMRFGRSDDYGHMRFGRK

> Ixodes pacificus sulfakinin (this work)
DDDYGHMRFGRSDDYGHMRFGRK

>KAG0410768.1:181-204 hypothetical protein HPB47_012117 [Ixodes persulcatus]
QDDDYGHMRFGRSDDYGHMRFGRK

>ACC99604.1:92-115 preprosulfakinin [Dermacentor variabilis]
QEDDYGHMRFGRSDDYGHMRFGRK

>XP_037563120.1:92-115 uncharacterized protein LOC119442349 [Dermacentor silvarum]
QEDDYGHMRFGRSDDYGHMRFGRK

>KAH8041160.1:330-353 hypothetical protein HPB51_013820 [Rhipicephalus microplus]
QEDDYGHMRFGRSDDYGHMRFGRK

>XP_037510364.1:94-117 uncharacterized protein LOC119387129 [Rhipicephalus sanguineus]
QEDDYGHMRFGRSDDYGHMRFGRK

>QYF10817.1:95-118 sulfakinin [Rhipicephalus microplus]
QEDDYGHMRFGRSDDYGHMRFGRK

>KAG5886106.1:83-109 hypothetical protein JTB14_031705 [Gonioctena quinquepunctata]
SDDYGHMRFGRKGEEVSDDYGHMRFGR

>KAG5886106.1:100-111 hypothetical protein JTB14_031705 [Gonioctena quinquepunctata]
DDYGHMRFGRSN

>XP_023029393.1:89-116 drosulfakinins isoform X2 [Leptinotarsa decemlineata]
DDYGHMRFGRKGEEISDDYGHMRFGRSD

>KAH8369195.1:112-140 hypothetical protein KR009_003854 [Drosophila setifemur]
RFDDYGHMRFGKRGGDDQSDDYGHMRFGR
```

>XP_022246095.1:103-130 uncharacterized protein LOC111086652 [Limulus polyphemus]
RFDDYGHMRFGRSGPAKKFDDYGHMRFG

>XP_013782337.1:94-127 drosulfakinins-like [Limulus polyphemus]
MRAIKRQFDDYGHMRFGRSNSMKKYDDYGHMRFG

>ALG35949.1:17-40 sulfakinin precursor, partial [Thermobia domestica]
DDYGHMRFGKREFDDYGHMRFGRS

>XP_043274514.1:67-90 uncharacterized protein LOC122410445 [Venturia canescens]
DDYGHMRFGKRDRFDDYGHMRFGR

>QHB80571.1:80-105 sulfakinin [Carabus violaceus]
QSDDYGHMRFGKREFDDYGHMRFGRS

>XP_030371026.1:118-147 drosulfakinins [Scaptodrosophila lebanonensis]
RFDDYGHMRFGKRGGDDQFDDYGHMRFGRK

>ALG35946.1:80-106 sulfakinin precursor, partial [Periplaneta americana]
QSDDYGHMRFGKREQFDDYGHMRFGRS

>CAL48349.1:104-130 preprosulfakinin [Gryllus bimaculatus]
QSDDYGHMRFGKREPFDDYGHMRFGRS

>XP_023701497.1:98-124 callisulfakinin [Cryptotermes secundus]
QSDDYGHMRFGKREQFDDYGHMRFGRS

>XP_044743583.1:97-121 drosulfakinins [Chrysoperla carnea]
DDYGHMRFGKREDRFDDYGHMRFGR

>XP_021939912.1:97-123 callisulfakinin [Zootermopsis nevadensis]
QSDDYGHMRFGKREQFDDYGHMRFGRS

>CAG9863328.1:79-107 unnamed protein product [Phyllotreta striolata]
SDDYGHLRFGRRNGEELSDDYGHMRFGRS

>CAG9837077.1:83-110 unnamed protein product [Diabrotica balteata]
SDDYGHLRFGRRNGEDISDDYGHMRFGR

>XP_028154559.1:83-110 callisulfakinin [Diabrotica virgifera virgifera]
SDDYGHLRFGRRNGEDISDDYGHMRFGR

>CAY61866.1:1-29 hypothetical protein, partial [Tityus discrepans]
DDDYGHLRFGRNAKGMAKKYDDYGHMRFG

>CAD7398328.1:213-240 unnamed protein product [Timema poppensis]
QSDDYGHMRFGKRGEQFDDYGHMRFGRS

>CAD7569749.1:147-174 unnamed protein product [Timema californicum]
QSDDYGHMRFGKRGEQFDDYGHMRFGRS

>CAD7260165.1:144-171 unnamed protein product [Timema shepardi]
QSDDYGHMRFGKRGEQFDDYGHMRFGRS

>CAD7394294.1:100-127 unnamed protein product [Timema cristinae]
QSDDYGHMRFGKRGEQFDDYGHMRFGRS

>CAD7425306.1:100-127 unnamed protein product [Timema monikensis]
QSDDYGHMRFGKRGEQFDDYGHMRFGRS

>KAG8190275.1:91-119 hypothetical protein JTE90_025789 [Oedothorax gibbosus]
DDYGHMRFGRSSATSREKKYDDYGHMRFG

>CAD7228614.1:98-122 unnamed protein product [Cyprideis torosa]
DDYGHMRFGKRADFDDYGHMRFGRS

>RZF40461.1:43-72 hypothetical protein LSTR_LSTR017186, partial [Laodelphax striatellus]
QSDDYGHMRFGKRGEVDDKFDDYGHMRFGR

>XP_023288532.1:56-81 callisulfakinin [Orussus abietinus]
DDYGHMRFGKRDQFDDYGHMRFGRGH

>XP_012138452.1:66-90 PREDICTED: uncharacterized protein LOC105662204 [Megachile rotundata]
DDYGHMRFGKREQFDDYGHMRFGRS

>XP_029034143.1:66-90 drosulfakinins [Osmia bicornis bicornis]
DDYGHMRFGKREQFDDYGHMRFGRS

```
>XP_015429701.1:66-89 PREDICTED: uncharacterized protein
LOC107186369 [Dufourea novaeangliae]
DDYGHMRFGKREQFDDYGHMRFGR

>XP_017763111.1:66-90 PREDICTED: callisulfakinin [Eufriesea
mexicana]
DDYGHMRFGKREQFDDYGHMRFGRS

>XP_043262104.1:66-89 uncharacterized protein LOC122402959
[Colletes gigas]
DDYGHMRFGKREQFDDYGHMRFGR

>KOC62825.1:66-89 Drosulfakinin [Habropoda laboriosa]
DDYGHMRFGKREHFDDYGHMRFGR

>XP_012255947.2:73-97 callisulfakinin [Athalia rosae]
SDDYGHMRFGKRDQFDDYGHMRFGR

>OAD53182.1:152-176 Callisulfakinin [Eufriesea mexicana]
DDYGHMRFGKREQFDDYGHMRFGRS

>XP_046734083.1:75-99 uncharacterized protein LOC124404187
[Diprion similis]
SDDYGHMRFGKREQFDDYGHMRFGR

>XP_015521780.1:75-99 uncharacterized protein LOC107225735
[Neodiprion lecontei]
SDDYGHMRFGKREQFDDYGHMRFGR

>XP_017792637.1:137-160 PREDICTED: uncharacterized protein
LOC108574539 [Habropoda laboriosa]
DDYGHMRFGKREHFDDYGHMRFGR

>AFW19802.1:106-135 sulfakinin [Nilaparvata lugens]
QSDDYGHMRFGKRGEADDKFDDYGHMRFGR

>KAH9402798.1:108-137 hypothetical protein TYRP_015555
[Tyrophagus putrescentiae]
DDYGHLRFGRSGTRIKYNDPDYGHMRFGRK

>XP_022203745.2:104-133 uncharacterized protein LOC111060391
[Nilaparvata lugens]
QSDDYGHMRFGKRGEADDKFDDYGHMRFGR

>RZF41044.1:103-132 hypothetical protein LSTR_LSTR002676
[Laodelphax striatellus]
QSDDYGHMRFGKRGEVDDKFDDYGHMRFGR
```

>XP_017879515.2:96-120 uncharacterized protein LOC108624620 [Ceratina calcarata]
DDYGHMRFGKREHFDDYGHMRFGRS

>XP_033330641.1:84-107 uncharacterized protein LOC117222819 [Megalopta genalis]
DDYGHMRFGKREQFDDYGHMRFGR

>XP_044752147.1:73-101 drosulfakinins [Coccinella septempunctata]
QIDDYGHMRFGKRGEDQFDDYGHMRFGRS

>XP_001993726.1:101-127 drosulfakinins [Drosophila grimshawi]
DDYGHMRFGKRNGDDQFDDYGHMRFGR

>SPP88329.1:131-159 blast:Drosulfakinins [Drosophila guanche]
RFDDYGHMRFGKRGGDDQFDDYGHMRFGR

>TMW49025.1:74-103 hypothetical protein DOY81_005913 [Sarcophaga bullata]
RFDDYGHMRFGKRGGEEQFDDYGHMRFGRS

>XP_011188135.1:113-142 callisulfakinin [Zeugodacus cucurbitae]
RFDDYGHMRFGKRGGEEQFDDYGHMRFGRS

>KNC25916.1:114-143 hypothetical protein FF38_10329 [Lucilia cuprina]
RFDDYGHMRFGKRGGEEQFDDYGHMRFGRS

>XP_034665366.1:116-144 drosulfakinins [Drosophila subobscura]
RFDDYGHMRFGKRGGDDQFDDYGHMRFGR

>KAH8304929.1:116-144 hypothetical protein KR018_005051 [Drosophila ironensis]
RFDDYGHMRFGKRGGDDNYDDYGHMRFGR

>XP_037807372.1:114-143 callisulfakinin [Lucilia sericata]
RFDDYGHMRFGKRGGEEQFDDYGHMRFGRS

>XP_034138343.1:116-144 drosulfakinins [Drosophila guanche]
RFDDYGHMRFGKRGGDDQFDDYGHMRFGR

>KAH8330731.1:115-143 hypothetical protein KR067_006906 [Drosophila pandora]
RFDDYGHMRFGKRGGDDQFDDYGHMRFGR

>KAH8250135.1:115-143 hypothetical protein KR026_005742 [Drosophila bipectinata]
RFDDYGHMRFGKRGGDDQFDDYGHMRFGR

>KAH8335432.1:115-143 hypothetical protein KR074_001933 [Drosophila pseudoananassae]
RFDDYGHMRFGKRGGDDQFDDYGHMRFGR

>XP_001953285.1:115-143 drosulfakinins [Drosophila ananassae]
RFDDYGHMRFGKRGGDDQFDDYGHMRFGR

>KAH8311455.1:114-142 hypothetical protein KR044_006452 [Drosophila immigrans]
RFDDYGHMRFGKRGGDEQFDDYGHMRFGR

>XP_002000913.1:114-142 drosulfakinins [Drosophila mojavensis]
RFDDYGHMRFGKRGNDEQFDDYGHMRFGR

>XP_030238368.1:114-142 drosulfakinins [Drosophila navojoa]
RFDDYGHMRFGKRGNDEQFDDYGHMRFGR

>XP_017133525.1:113-141 drosulfakinins [Drosophila elegans]
RFDDYGHMRFGKRGGDDQFDDYGHMRFGR

>KAH8407295.1:113-141 hypothetical protein KR215_001129 [Drosophila sulfurigaster]
RFDDYGHMRFGKRGGDEQFDDYGHMRFGR

>XP_001978781.1:113-141 drosulfakinins [Drosophila erecta]
RFDDYGHMRFGKRGGDDQFDDYGHMRFGR

>XP_016985391.1:113-141 drosulfakinins [Drosophila rhopaloa]
RFDDYGHMRFGKRGGDDQFDDYGHMRFGR

>XP_002102055.1:113-141 drosulfakinins [Drosophila simulans]
RFDDYGHMRFGKRGGDDQFDDYGHMRFGR

>XP_030559213.1:113-141 drosulfakinins [Drosophila novamexicana]
RFDDYGHMRFGKRGGDDQFDDYGHMRFGR

>XP_034114793.1:113-141 drosulfakinins [Drosophila albomicans]
RFDDYGHMRFGKRGGDEQFDDYGHMRFGR

>ACC99373.1:113-141 drosulfakinin [Drosophila simulans]
RFDDYGHMRFGKRGGDDQFDDYGHMRFGR

```
>XP_017052006.1:113-141 drosulfakinins-like [Drosophila
ficusphila]
RFDDYGHMRFGKRGGDDQFDDYGHMRFGR

>XP_017855511.1:113-141 PREDICTED: drosulfakinins [Drosophila
arizonae]
RFDDYGHMRFGKRGNDEQFDDYGHMRFGR

>XP_002038249.1:113-141 drosulfakinins [Drosophila sechellia]
RFDDYGHMRFGKRGGDDQFDDYGHMRFGR

>KAH8381277.1:113-141 hypothetical protein KR093_001714
[Drosophila rubida]
RFDDYGHMRFGKRGGDEQFDDYGHMRFGR

>NP_524845.2:113-141 drosulfakinin [Drosophila melanogaster]
RFDDYGHMRFGKRGGDDQFDDYGHMRFGR

>XP_034485331.1:113-141 drosulfakinins [Drosophila innubila]
RFDDYGHMRFGKRGGDEQFDDYGHMRFGR

>XP_002053743.1:113-141 drosulfakinins [Drosophila virilis]
RFDDYGHMRFGKRGGDDQFDDYGHMRFGR

>XP_017056881.1:113-141 drosulfakinins [Drosophila ficusphila]
RFDDYGHMRFGKRGGDDQFDDYGHMRFGR

>KAH8344502.1:113-141 hypothetical protein KR084_012843
[Drosophila pseudotakahashii]
RFDDYGHMRFGKRGGDDQFDDYGHMRFGR

>Q7M3V5.1:112-139 [Calliphora vomitoria]
DDYGHMRFGKRGGEEQFDDYGHMRFGRS

>XP_017015431.1:112-140 drosulfakinins [Drosophila takahashii]
RFDDYGHMRFGKRGGDDQFDDYGHMRFGR

>XP_014097487.1:110-139 callisulfakinin [Bactrocera oleae]
RFDDYGHMRFGKRGGEEQFDDYGHMRFGRS

>XP_004529342.1:110-139 callisulfakinin [Ceratitis capitata]
RFDDYGHMRFGKRGGEDQFDDYGHMRFGRS

>XP_001359636.3:111-139 drosulfakinins [Drosophila
pseudoobscura]
RFDDYGHMRFGKRGGDDQFDDYGHMRFGR
```

>KAH8283209.1:111-139 hypothetical protein KR054_011942 [Drosophila jambulina]
RFDDYGHMRFGKRGGDDQFDDYGHMRFGR

>Q295C6.2:111-139 [Drosophila pseudoobscura pseudoobscura]
RFDDYGHMRFGKRGGDDQFDDYGHMRFGR

>KAH8420883.1:111-139 hypothetical protein KR222_008311 [Zaprionus bogoriensis]
RFDDYGHMRFGKRGGDEQFDDYGHMRFGR

>KAH8264670.1:113-139 hypothetical protein KR038_011950 [Drosophila bunnanda]
RFDDYGHMRFGKRGGDDQFDDYGHMRFGR

>XP_017143141.1:111-139 drosulfakinins [Drosophila miranda]
RFDDYGHMRFGKRGGDDQFDDYGHMRFGR

>XP_017020401.1:111-139 drosulfakinins [Drosophila kikkawai]
RFDDYGHMRFGKRGGDDQFDDYGHMRFGR

>XP_020813064.1:111-139 drosulfakinins [Drosophila serrata]
RFDDYGHMRFGKRGGDDQFDDYGHMRFGR

>XP_039492415.1:110-138 drosulfakinins [Drosophila santomea]
RFDDYGHMRFGKRGGDDQFDDYGHMRFGR

>XP_043655408.1:110-138 drosulfakinins [Drosophila teissieri]
RFDDYGHMRFGKRGGDDQFDDYGHMRFGR

>XP_039970427.1:108-137 drosulfakinins [Bactrocera tryoni]
RFDDYGHMRFGKRGGEEQFDDYGHMRFGRS

>XP_018802062.1:108-137 PREDICTED: drosulfakinins [Bactrocera latifrons]
RFDDYGHMRFGKRGGEEQFDDYGHMRFGRS

>XP_011206252.1:108-137 drosulfakinins [Bactrocera dorsalis]
RFDDYGHMRFGKRGGEEQFDDYGHMRFGRS

>XP_037729864.1:109-137 drosulfakinins [Drosophila subpulchrella]
RFDDYGHMRFGKRGGDDQFDDYGHMRFGR

>XP_016950713.1:109-137 drosulfakinins [Drosophila biarmipes]
RFDDYGHMRFGKRGGDDQFDDYGHMRFGR

>XP_002095906.1:109-137 drosulfakinins [Drosophila yakuba]
RFDDYGHMRFGKRGGDDQFDDYGHMRFGR

>XP_016932271.1:109-137 drosulfakinins [Drosophila suzukii]
RFDDYGHMRFGKRGGDDQFDDYGHMRFGR

>KAH8255046.1:108-136 hypothetical protein KR032_008242 [Drosophila birchii]
RFDDYGHMRFGKRGGDDQFDDYGHMRFGR

>XP_005188371.2:106-135 PREDICTED: callisulfakinin [Musca domestica]
RFDDYGHMRFGKRGGEEQFDDYGHMRFGRS

>XP_017484043.1:104-133 PREDICTED: drosulfakinins [Rhagoletis zephyria]
RFDDYGHMRFGKRGGEEQFDDYGHMRFGRS

>XP_036318820.1:103-132 callisulfakinin-like [Rhagoletis pomonella]
RFDDYGHMRFGKRGDEEQFDDYGHMRFGRS

>XP_017846232.1:103-131 drosulfakinins [Drosophila busckii]
RFDDYGHMRFGKRGGEDQFDDYGHMRFGR

>AAB03703.1:100-128 drosulfakinin precursor [Drosophila melanogaster]
RFDDYGHMRFGKRGGDDQFDDYGHMRFGR

>CAB3373177.1:94-123 Hypothetical predicted protein [Cloeon dipterum]
DDYGHMRFGKRTQQAAGGFDDYGHMRFGRS

>XP_013116087.1:83-112 PREDICTED: callisulfakinin [Stomoxys calcitrans]
RFDDYGHMRFGKRGGEEQFDDYGHMRFGRS

>KOX81045.1:23-46 hypothetical protein WN51_09970, partial [Melipona quadrifasciata]
DDYGHMRFGKREQFEDYGHMRFGR

>XP_046677204.1:57-84 drosulfakinins [Homalodisca vitripennis]
DDYGHMRFGKRGDPEDKFDDYGHMRFGR

>XP_002424661.1:63-90 drosulfakinins precursor, putative [Pediculus humanus corporis]
QDQFDDYGHMRFGKRGEDLDDYGHMRFG

>XP_041775538.1:166-195 uncharacterized protein LOC121595558 [Anopheles merus]
RFDDYGHMRFGKRGGEGDQFDDYGHMRFGR

>XP_040236726.1:167-194 uncharacterized protein LOC120958178 [Anopheles coluzzii]
DDYGHMRFGKRGGEGDQFDDYGHMRFGR

>XP_001238248.1:165-194 AGAP009275-PA [Anopheles gambiae str. PEST]
RFDDYGHMRFGKRGGEGDQFDDYGHMRFGR

>XP_040171403.1:164-193 uncharacterized protein LOC120904939 [Anopheles arabiensis]
RFDDYGHMRFGKRGGEGDQFDDYGHMRFGR

>AAR03495.1:164-193 sulfakinin preproprotein [Anopheles gambiae]
RFDDYGHMRFGKRGGEGDQFDDYGHMRFGR

>KFB42310.1:158-187 sulfakinin [Anopheles sinensis]
RFDDYGHMRFGKRGGEGDQFDDYGHMRFGR

>AAW82713.1:156-185 sulfakinin [Anopheles maculatus]
RFDDYGHMRFGKRGGEGDQFDDYGHMRFGR

>XP_035909976.1:156-185 uncharacterized protein LOC118511243 [Anopheles stephensi]
RFDDYGHMRFGKRGGEGDQFDDYGHMRFGR

>EDS43001.1:116-145 sulfakinin [Culex quinquefasciatus]
RFDDYGHMRFGKRGGEGDQFDDYGHMRFGR

>XP_039431740.1:116-145 drosulfakinins [Culex pipiens pallens]
RFDDYGHMRFGKRGGEGDQFDDYGHMRFGR

>XP_002070454.3:112-141 drosulfakinins [Drosophila willistoni]
RFDDYGHMRFGKRGGGEDQFDDYGHMRFGR

>CAG9798060.1:84-113 unnamed protein product [Chironomus riparius]
RFDDYGHMRFGKRGVGEDGFDDYGHMRFGR

>CAD7421984.1:100-127 unnamed protein product [Timema poppensis]
QSDDYGHMRFGKRGEQLDDYGHMRFGHS

>CAD7207579.1:100-127 unnamed protein product [Timema douglasi]

QSDDYGHMRFGKRGEQLDDYGHMRFGHS

>XP_043511406.1:66-89 drosulfakinins [Frieseomelitta varia]
DDYGHMRFGKREQFEDYGHMRFGR

>EFX80896.1:181-203 putative sulfakinin-like peptide [Daphnia pulex]
DDYGHMRYGKRDFDDYGHMRFGR

>CAH0102591.1:142-164 unnamed protein product [Daphnia galeata]
DDYGHMRYGKRDFDDYGHMRFGR

>XP_046635393.1:141-163 drosulfakinins-like [Daphnia pulicaria]
DDYGHMRYGKRDFDDYGHMRFGR

>XP_032778356.2:140-162 uncharacterized protein LOC116917097 [Daphnia magna]
DDYGHMRYGKRDFDDYGHMRFGR

>KXJ72000.1:119-148 hypothetical protein RP20_CCG019163 [Aedes albopictus]
RFDDYGHMRFGKRGGGGEGEQFDDYGHMRFGR

>XP_021703537.1:116-147 drosulfakinins [Aedes aegypti]
RFDDYGHMRFGKRGGGGEGEQFDDYGHMRFGR

>CAD1474980.1:120-143 unnamed protein product, partial [Heterotrigona itama]
DDYGHMRFGKREQFEDYGHMRFGR

>ODN04808.1:75-99 Drosulfakinin [Orchesella cincta]
DDYGHMRFGKRNPKDFDDYGHMRFG

>KAF8764284.1:78-107 hypothetical protein HNY73_022373 [Argiope bruennichi]
EEDDYGHMRFGRSNANMSADYDHVHMRFGR

>XP_022919257.1:90-112 drosulfakinins [Onthophagus taurus]
DDYGHLRFGKREFDDYGHMRFGR

>GFX43891.1:87-114 uncharacterized protein TNCV_4111701 [Trichonephila clavipes]
DDYGHMRFGRNAGMAQKKFDDYGHMRYG

>GIY02054.1:87-114 uncharacterized protein CDAR_229461 [Caerostris darwini]
DDYGHMRFGRNVGGPHKKFDDYGHMRYG

>GBL89615.1:87-114 hypothetical protein AVEN_104585-1 [Araneus ventricosus]
DDYGHMRFGRNAGVPHKKYDDYGHMRYG

>GFQ77535.1:87-114 uncharacterized protein TNCT_388621 [Trichonephila clavata]
DDYGHMRFGRNAGMTQKKFDDYGHMRYG

>GFY72129.1:87-114 uncharacterized protein TNIN_87961 [Trichonephila inaurata madagascariensis]
DDYGHMRFGRNAGMAQKKFDDYGHMRYG

>KAF8764970.1:87-114 Callisulfakinin like protein [Argiope bruennichi]
DDYGHMRFGRNAGVPHKKFDDYGHMRYG

>GFU25137.1:87-114 uncharacterized protein NPIL_394041 [Nephila pilipes]
DDYGHMRFGRNAGMAQKKFDDYGHMRYG

>XP_015601843.1:65-88 uncharacterized protein LOC107270914 [Cephus cinctus]
DDYGHMRFGKRDQFDDYGHLRFGR

>XP_043467793.1:69-92 drosulfakinins [Leptopilina heterotoma]
DDYGHMRFGKRDQFEDYGHLRFGR

>XP_031848655.1:131-154 uncharacterized protein LOC116434061 [Nomia melanderi]
DDYGHMRFGKREEYDDYGHLRFGR

>XP_031618783.1:109-136 drosulfakinins [Contarinia nasturtii]
RYDDYGHMRFGKRGDSDQFDDYGHMRFG

>XP_037955757.1:101-128 drosulfakinins [Teleopsis dalmanni]
RFDDYGHMRFGKRGGDEQFDDYGHMRFG

>KAF7280388.1:98-123 hypothetical protein GWI33_006119 [Rhynchophorus ferrugineus]
SDDYGHLRFGKRDEQFDDYGHMRFGR

>XP_037024525.1:91-118 drosulfakinins [Bradysia coprophila]
RYDDYGHMRFGKRGGEDQFDDYGHMRFG

>QGA72577.1:84-109 drosulfakinins [Rhynchophorus ferrugineus]
SDDYGHLRFGKRDEQFDDYGHMRFGR

>QXU63184.1:86-111 sulfakinin [Dendroctonus armandi]
SDDYGHLRFGKREEQVDDYGHMRFGR

>CAG9767660.1:86-111 unnamed protein product [Ceutorhynchus assimilis]
SDDYGHLRFGKRDEQFDDYGHMRFGR

>XP_019762103.2:85-110 drosulfakinins [Dendroctonus ponderosae]
SDDYGHLRFGKREEQFDDYGHMRFGR

>KAI4465698.1:63-88 sulfakinin family [Holotrichia oblita]
DDYGHLRFGKRGDEQFDDYGHMRFGR

>BAH11170.1:82-108 preprosulfakinin [Psacothea hilaris hilaris]
SDDYGHLRFGKRGEESFDDYGHMRFGR

>XP_018561061.1:82-108 drosulfakinins [Anoplophora glabripennis]
SDDYGHLRFGKRGEESFDDYGHMRFGR

>XP_008194373.1:83-109 PREDICTED: drosulfakinins [Tribolium castaneum]
SDDYGHLRFGKRGEEPFDDYGHMRFGR

>XP_044268639.1:84-111 drosulfakinins [Tribolium madens]
SDDYGHLRFGKRGEEPFDDYGHMRFGRS

>XP_023238955.1:93-120 uncharacterized protein LOC111637646 [Centruroides sculpturatus]
DDYGHLRFGRNAKGMAKKYDDYGHMRFG

>RZB38975.1:84-110 hypothetical protein BDFB_002382 [Asbolus verrucosus]
SDDYGHLRFGKRGEEAFDDYGHMRFGR

>CAH1368946.1:85-112 unnamed protein product [Tenebrio molitor]
SDDYGHLRFGKRGEEPFDDYGHMRFGRS

>KAG5676024.1:88-118 hypothetical protein PVAND_005879 [Polypedilum vanderplanki]
RYDDYGHMRFGKRAGPGDEGFDDYGHLRFGR

>KAF6214438.1:137-164 hypothetical protein GE061_009181 [Apolygus lucorum]
NDYGHMRFGKRSGGEEKFDDYGYMRFGR

>ALQ28597.1:68-96 sulfakinin, partial [Scylla paramamosain]

DDYGHMRFGKRGSGNDDYQDDYGHLRFGR

>XP_034230796.1:97-124 uncharacterized protein LOC117639338 [Thrips palmi]
SDDYGHMRFGKRGGENDFAEYGHMRFGR

>XP_045133011.1:85-113 uncharacterized protein LOC123517134 [Portunus trituberculatus]
DDYGHMRFGKRGSASDDYQDDYGHLRFGR

>CRL00088.1:516-546 CLUMA_CG013370, isoform A [Clunio marinus]
RFDDYGHLRFGKRGGGEGDGFDDYGHMRFGR

>XP_006557714.2:80-103 uncharacterized protein LOC102654729 [Apis mellifera]
DDYGHLRFGKREQFEDYGHMRFGR

>XP_006618228.1:81-105 uncharacterized protein LOC102676428 [Apis dorsata]
DDYGHLRFGKREQFEDYGHMRFGRS

>XP_012346753.2:81-105 uncharacterized protein LOC105736501, partial [Apis florea]
DDYGHLRFGKREQFEDYGHMRFGRS

>KAE8744395.1:79-107 Sulfakinin [Frankliniella occidentalis]
SDDYGHMRFGKRGGGDNDFPEYGHMRFGR

>XP_016922928.2:88-112 uncharacterized protein LOC108004484 [Apis cerana]
DDYGHLRFGKREQFEDYGHMRFGRS

>XP_026282628.1:93-121 drosulfakinins [Frankliniella occidentalis]
SDDYGHMRFGKRGGGDNDFPEYGHMRFGR

>XP_022256192.1:114-141 uncharacterized protein LOC111088983 [Limulus polyphemus]
RFDDYGHLRFGRSGPAEKTDDYGYMRFG

>XP_043800897.1:137-161 uncharacterized protein LOC122719285 [Apis laboriosa]
DDYGHLRFGKREQFEDYGHMRFGRS

>PBC25365.1:143-16S Callisulfakinin [Apis cerana cerana]
DDYGHLRFGKREQFEDYGHMRFGRS

>XP_019873919.1:80-108 PREDICTED: uncharacterized protein LOC109602040 [Aethina tumida]
SEDYGHLRFGKRDEQFDDYGHMRFGRGGD

>XP_045604047.1:77-102 uncharacterized protein LOC123761871 [Procambarus clarkii]
DEYGHMRFGKRGGDYDDYGHLRFGRS

>AWK57545.1:79-104 sulfakinin [Cherax quadricarinatus]
DEYGHMRFGKRGGDYDDYGHLRFGRS

>PSN53545.1:98-124 hypothetical protein C0J52_09179 [Blattella germanica]
QSEDYGHFRFGKREQFDDYGHMRFGRS

>KAG7170016.1:76-102 sulfakinin-like [Homarus americanus]
DEYGHMRFGKRGGGEYDDYGHLRFGRS

>QBX89077.1:79-105 sulfakinin [Nephrops norvegicus]
DEYGHMRFGKRGGVDYDDYGHLRFGRS

>XP_042220086.1:79-105 uncharacterized protein LOC121864942 [Homarus americanus]
DEYGHMRFGKRGGGEYDDYGHLRFGRS

>CAD7646181.1:32-66 unnamed protein product [Medioppia subpectinata]
DDYGHLRFGRNDPSVPQLDARVKYNDPDYGHMRFG

>QTE34438.1:74-99 sulfakinin [Cataglyphis nodus]
DDYGHMRFGKRSNGDDEEYGHSRFGR

>XP_029665461.1:74-99 uncharacterized protein LOC115236882 [Formica exsecta]
DDYGHMRFGKRSNGDDEEYGHSRFGR

>XP_012540957.2:100-125 uncharacterized protein LOC105839297 [Monomorium pharaonis]
DDYGHMRFGKRSNGDDEEYGHSRFGR

>RXG52591.1:46-70 Drosulfakinin [Armadillidium vulgare]
DDYGHLRFGKRKDFDDYGHLRFGRS

>CAG7817686.1:79-105 unnamed protein product [Allacma fusca]
DDYGHMRFGKRNPPGKEFDDYGHLRFG

>XP_035712620.1:85-111 drosulfakinins isoform X1 [Folsomia candida]
DDYGHMRFGKRNQPSKDFDDYGHLRFG

>CAD7241566.1:87-111 unnamed protein product [Darwinula stevensoni]
DDYGHMRFGKRADFDEYGHSRFGRS

>KAF5289009.1:57-82 hypothetical protein FQA39_LY03888 [Lamprigera yunnana]
DDYGHLRFGKRGEDFDDYGHLRFGRS

>KAF5286789.1:66-90 hypothetical protein FQR65_LT02207 [Abscondita terminalis]
DDYGHLRFGKRGEDFDDYGHLRFGR

>XP_031344997.1:102-126 uncharacterized protein LOC116172042 [Photinus pyralis]
DDYGHLRFGKRGEDFDDYGHLRFGR

>KAF2881883.1:118-143 hypothetical protein ILUMI_24288 [Ignelater luminosus]
DDYGHLRFGKRGEDFDDYGHLRFGRS

>XP_029162716.1:74-99 uncharacterized protein LOC114934234 [Nylanderia fulva]
DDYGHMRFGKRSNGNDEEYGHSRFGR

>CAD7659273.1:91-128 unnamed protein product [Oppiella nova]
DDYGHLRFGRNDPNMQSADGLNSRVKYNDPDYGHMRFG

>XP_027208253.1:79-111 uncharacterized protein LOC113801964 [Penaeus vannamei]
DEYGHMRFGKRAGGSGGVGGEYDDYGHLRFGRS

>XP_042870020.1:79-111 uncharacterized protein LOC122251873 [Penaeus japonicus]
DEYGHMRFGKRAGGSGGVGGEYDDYGHLRFGRS

>XP_025267330.1:78-102 uncharacterized protein LOC105251829 isoform X1 [Camponotus floridanus]
DDYGHMRFGKRSNNEDEEYGHLRFG

>XP_022242867.1:106-131 uncharacterized protein LOC111085996 [Limulus polyphemus]
DDYGHMRFGRNGRMKKHDEYGHMYFG

>RWS29032.1:52-79 hypothetical protein B4U80_00988 [Leptotrombidium deliense]
DDYGHLRFGRNEPRIKMSDPDYGHLRFG

>XP_012232058.1:74-99 PREDICTED: uncharacterized protein LOC105677777 isoform X2 [Linepithema humile]
DDYGHMRFGKRSNGEDEEYGHPRFGR

>XP_018394343.1:74-99 PREDICTED: uncharacterized protein LOC108773120 [Cyphomyrmex costatus]
DDYGHMRFGKRSNGDDEEYGHLRLGR

>XP_012232049.1:78-103 PREDICTED: uncharacterized protein LOC105677777 isoform X1 [Linepithema humile]
DDYGHMRFGKRSNGEDEEYGHPRFGR

>XP_011633090.2:78-103 uncharacterized protein LOC105424513 [Pogonomyrmex barbatus]
DDYGHMRFGKRTNSNDEEYGHPRFGR

>XP_017780371.1:86-109 PREDICTED: uncharacterized protein LOC108565434 [Nicrophorus vespilloides]
DDYGHLRFGKREDFDDYGHLRYGR

>ACS45388.1:62-90 sulfakinin precursor [Rhodnius prolixus]
NEYGHMRFGKRGGSDEKFDDYGYMRFGRS

>XP_014274494.1:48-76 drosulfakinins [Halyomorpha halys]
NDYGHMRFGKRGGVTEDKFEDYGYMRFGR

>BAV78829.1:48-76 sulfakinin [Plautia stali]
NDYGHMRFGKRGGVTEDKFEDYGYMRFGR

>AZK31368.1:48-76 neuropeptide precursor [Nezara viridula]
NDYGHMRFGKRGGVTEDKFEDYGYMRFGR

>KAH0944647.1:62-85 hypothetical protein HN011_005536 [Eciton burchellii]
DDYGHMRFGKRSRSDHAGYGHLRF

>GBM16231.1:81-96 hypothetical protein AVEN_195347-1 [Araneus ventricosus]
QDDDYGHMRFGRSNAY

>KMQ94624.1:74-99 sulfakinin [Lasius niger]
DDYGHMRFGKRSNGNDEEYGHSRFAR

>XP_011686925.1:74-98 PREDICTED: uncharacterized protein LOC105449408 [Wasmannia auropunctata]
DDYGHMRFGKRSNGDDEEYGHPRFG

>XP_018316699.1:74-99 PREDICTED: uncharacterized protein LOC108731103 [Trachymyrmex zeteki]
DDYGHMRFGKRSNGDDEEYGHSRLGR

>XP_047001071.1:70-106 drosulfakinins-like [Schistocerca americana]
SDDYGHMRFGKRQPAAPAAAPVPVAPRFDDYGHFRFG

>KAA0193774.1:48-70 hypothetical protein HAZT_HAZT006110 [Hyalella azteca]
DYGHLRFGKRAEFGDYGHLRFGR

>UGX04242.1:67-105 sulfakinin [Schistocerca gregaria]
SDDYGHMRFGKRQPAAAPAAAAAPVPVAPRFDDYGHFRFG

>CAH2066757.1:47-75 unnamed protein product, partial [Iphiclides podalirius]
DYALMRSRDTRADDSFDDYGHMRFGRSDD

>XP_014251620.1:66-94 drosulfakinins [Cimex lectularius]
NDYGHMRFGKREGTDEKFEDYGYLRFGRS

>XP_025987062.1:73-97 drosulfakinins [Solenopsis invicta]
DDYGHMRFGKRSNGDDEEFGHPRFG

>EGI60712.1:74-99 hypothetical protein G5I_11077 [Acromyrmex echinatior]
DDYGHMRFGKRFNGDDEEYGHSRLGR

>KAG5334666.1:74-99 DSK protein, partial [Acromyrmex heyeri]
DDYGHMRFGKRFNGDDEEYGHSRLGR

>KAG5311275.1:74-99 DSK protein, partial [Acromyrmex insinuator]
DDYGHMRFGKRFNGDDEEYGHSRLGR

>XP_018058795.1:74-99 PREDICTED: uncharacterized protein LOC108694052 [Atta colombica]
DDYGHMRFGKRFNGDDEEYGHSRLGR

>KAG5312046.1:74-99 DSK protein, partial [Pseudoatta argentina]
DDYGHMRFGKRFNGDDEEYGHSRLGR

>XP_018350001.1:74-99 PREDICTED: uncharacterized protein LOC108753149 [Trachymyrmex septentrionalis]
DDYGHMRFGKRFNGDDEEYGHSRLGR

>XP_011062258.1:74-99 PREDICTED: uncharacterized protein LOC105150704 [Acromyrmex echinatior]
DDYGHMRFGKRFNGDDEEYGHSRLGR

>KAG8200871.1:79-94 hypothetical protein JTE90_015776 [Oedothorax gibbosus]
SEDDYGHMRFGRSDSY

>XP_011863280.1:74-97 PREDICTED: drosulfakinins [Vollenhovia emeryi]
DDYGHMRFGKRSNGDEEEYGHPRF

>XP_018374161.1:74-99 PREDICTED: drosulfakinins [Trachymyrmex cornetzi]
DDYGHMRFGKRFNGDDEEYAYLRFGR

>XP_022912824.1:138-155 FMRFamide neuropeptides [Onthophagus taurus]
NDNHMRFGRTDNYMRFGR

>EZA62204.1:73-94 hypothetical protein X777_02830 [Ooceraea biroi]
DDYGHMRFGRRSLGDDIGYGHL

>XP_026831426.1:105-126 uncharacterized protein LOC105279135 isoform X1 [Ooceraea biroi]
DDYGHMRFGRRSLGDDIGYGHL

>GFY55800.1:86-98 uncharacterized protein TNIN_280661 [Trichonephila inaurata madagascariensis]
DDDYGHMRFGRSN

>GFR15512.1:86-98 uncharacterized protein TNCT_578721 [Trichonephila clavata]
DDDYGHMRFGRSN

>GFS72434.1:120-132 uncharacterized protein TNCV_2115921 [Trichonephila clavipes]
DDDYGHMRFGRSN

>GFT96555.1:97-108 uncharacterized protein NPIL_445641 [Nephila pilipes]
DDDYGHMRFGRS

>KAG8113877.1:54-66 hypothetical protein SFRUCORN_001037 [Spodoptera frugiperda]
DDYGHLRFGRSDD

>KPJ15107.1:34-70 hypothetical protein RR48_09134 [Papilio machaon]
DDDEYRARRLYREYGMRRAARADDSFDDYGHLRFGRS

>KPI93793.1:34-72 hypothetical protein RR46_12958 [Papilio xuthus]
DDDEYRARRLYREYGMRRAARADDSFDDYGHLRFGRSDD

>XP_028159220.1:61-73 uncharacterized protein LOC114352007 [Ostrinia furnacalis]
DDYGHLRFGRSDD

>AXY04298.1:62-74 sulfakinin [Spodoptera exigua]
DDYGHLRFGRSDD

>XP_021184067.1:62-74 uncharacterized protein LOC110371929 [Helicoverpa armigera]
DDYGHLRFGRSDD

>XP_028036696.1:46-74 drosulfakinins [Bombyx mandarina]
DYGLIRSRVIRGDDTFDDYGHLRFGRSDD

>CAB3231046.1:62-74 unnamed protein product [Arctia plantaginis]
DDYGHLRFGRSDD

>BAG49564.1:46-74 sulfakinin precursor [Bombyx mori]
DYGLIRSRVIRGDDTFDDYGHLRFGRSDD

>XP_031763859.1:47-75 drosulfakinins-like [Galleria mellonella]
DYSLIRGRVARADDSFDDYGHLRFGRSDD
>CAH1644921.1:72-84 unnamed protein product [Spodoptera littoralis]
DDYGHLRFGRSDD

>XP_047041638.1:74-86 uncharacterized protein LOC124645785 [Helicoverpa zea]
DDYGHLRFGRSDD

>CAD0196156.1:79-91 unnamed protein product [Chrysodeixis includens]
DDYGHLRFGRSDD

```
>OTF78416.1:100-125 hypothetical protein BLA29_005187
[Euroglyphus maynei]
HLRFGKRFNARAKYSDLDYGHMRFGR

>KAF2366247.1:172-197 hypothetical protein FHG87_002983
[Trinorchestia longiramus]
DYGHLRFGKRAEFSDYGHLGFGRSQD

>XP_020299794.1:76-101 uncharacterized protein LOC109863729
[Pseudomyrmex gracilis]
DTEYGHWRFGKRSNDDEEYGHSRFGR
```

#### Amino acid sequences of sulfakinin receptors used in alignments

>Sulfakinin receptor [Ixodes scapularis] (this work)
METCNLSTEGNGSDARPAYSWWRSDQAVLVAPYTVILLLAVLGNGLVIVTLAVNKRMRTVTNLFLLNLAVSDL
LLGVFCMPFTLAGVLLREFVFGELMCRLIPYLQAVSVCVSAWTLMAMSVERYFAICYPLRSRTWQTLRHAQRT
IGAVWVASLLLMLPIALLSQLQPVKDSGKMKCREDWREPLYERLFTLFLDALLLVLPLLGMTATYSRIAATLR
SAMHHAPSDVAWHNGSAFSAVHPADQQPASRLSHQPSTIRWHQRDQDRSLATKQRIIRMLFAVVVEFFVCWTP
LYVLNTVSLFQPEAVYYGLGYRGISFLQLLAYASSCCNPITYCFMNRTFRVSFLGLCRHCCRTKGKLGTSRHS
TKQSFVYVVGQEAQ

>CAL6254493.1 sulfakinin receptor[Ixodes persulcatus]
METWNLSTEGNGSDARPAYAWWRSDQAVLVAPYTVILLLAVLGNGLVIVTLAVNKRMRTVTNLFLLNLAVSDL
LLGVFCMPFTLAGVLLREFVFGELMCRLIPYLQAVSVCVSAWTLMAMSVERYFAICYPLRSRTWQTLRHAQRT
IAAVWMASLLLMLPIALLSQLQPVKDSGKMKCREDWREPLYERLFTLFLDALLLVLPLLGMTATYSRIAATLR
SAMHHAPSDAAWHNGSALGTVHPADQQPASRLSHQPSTIRWHQRDQDRSLATKQRIIRMLFAVVVEFFVCWTP
LYVLNTVSLFRPEAVYYGLGYRGISFLQLLAYASSCCNPITYCFMNRTFRVSFLGLCRHCCRTKGNLGTSRHS
TKQSFVYVVGQETQ

>CAL6193878.1 sulfakinin receptor [Ixodes hexagonus]
YSWWRSDQAVLVAPYTVILLLAVLGNGLVIVTLAVNKRMRTVTNLFLLNLAVSDLLLGVFCMPFTLAGVLLRE
FVFGELMCRLIPYLQAVSVCVSAWTLMAMSVERYFAICYPLRSRTWQTLRHAQRTIAAVWVASLLLMLPIALL
SHLQPVKDSGKMKCREDWRDPLYERLFTLFLDALLLVLPLLGMIATYSRIAATLRSAMHHAPTDVACQNGSGL
AAVHPADRQPTPRLSHQPSTIRWHQRDHDRSLATKQRIIRMLFAVVVEFFVCWTPLYVLNTVSLFQPEAVYYG
LGYRGISFLQLLAYASSCCNPITYCFMNRTFRVSFLGLCRHCCRTKDRLSSSRHSVKQSYVYVVGQDTP

>CAL6256592.1 sulfakinin [Ixodes pacificus]
GSAIGAVHPADQQPASRLSHQPSTIRWHQRDQDRSLATKQRIIRMLFAVVVEFFVCWTPLYVLNTVSLFQPEA
VYYGLGYRGISFLQLLAYASSCCNPITYCFMNRTFRVSFLGLCRHCCRTKGKLGTSRHSTKQSFVYVVGQETQ

>XP_037270673.1 SKR type A-like [Rhipicephalus microplus]
MSGPLQREQQAVGKGKGSMAPGISGRPPNDSKPSTTASLICLHTLRSGCSAGHYERXNLSEAVLVAPYAIILL
LAVLGNGLVILTLAVNQRMRTVTNLFLLNLAVSDLLLGVFCMPFTLAGVLLREFVFGQLLCNLIPYLQAVSVC
VSDWTLVAMSVERYYAICEPLRSRSWQTPRHAQRTVATVWTVSFLLMLPIALLSELRPIRDSRKMKCREDWGD
LLYERLFTLFLDVLLLVLPLLIMVTTYARIAATLRSAMQQQQQQSEEATGSPKHNGSPLAAVHPAEREASPS
LRMRRVATVRWHQRDSDRSLATKQRIIRMLFVVVVEFFVCWTPLYVLNTVSLFRPEAVYYGLGYRGICFLQLL
AYASSCCNPITYCFMNRTFRVSFLSLCHRRKPKQGPGSVRSLTKKDGVYFLDREKN

>XP_037277149.1 SKR type B receptor-like isoform X1 [Rhipicephalus microplus]
MCRGAESTNASGDALSNETAAAEALASAAAACWTGISFETAFRILLYAIIFVFAVVGNSLVIITLVQRRRMRT
VTNVFLLNLAVSDLLLGVFCMPTTLVGSVLRNFVLGPVMCKLIPYFQAVSVSVSAWTLVSISLERFFAIVRPL
ESRRWQTRSHAYKVILLVWLCSMVTMLPIAVLSQLVPLRGERKKCREVWPDATSERVFNVYLDVTLFLLPLII
MSFVYSCICATLCHGMKLDNKSTLCCCSLVRRALARRCRGCAAPTPETEQHPSNEELMSTAAVHEEYHTNPNM
PQLFAPPDVEARLQRTPRRQFIFRGTYRKSRASKRRVIRMLSVLVLEFFVCWTPLYVLHTWTVFDAHAAYSRV
PAGAFAAVHLLAYVSSCCNPITYCFMHDKYRQAFRNVLACSSAASRRRRRWSSTKSHDRFVSLNSICSNGKNA
SGKGSLRSTLTSRLSSLKEEKPTNDINEVADQKHAAVCKLPATSLADGESSSSCNMSTYCGRRAGNASDTES

>XP_037289624.1 SKR1C partial [Rhipicephalus microplus]
MVNRLISSLPLLNLQWPLFNKTTSSPFTEPFAIPTVTLPDDAVFTEFPQGSVQNFSTPQSESTGAVDGLPDAT
DLIVRVGLYVVIFVFAVVGNVLVLVTLVQNKRMRTVTNVFLVNLAVSDLLLGVLCMPFTLIGSLLRNFVFGEL
MCKFIPYLQAVSVSVSVWTLVSISMERFFAIVRPLQSRRWQTVSHAYRVIAVIWVLSLVHVAPIAVLSQLIPT
RTAGHQKCRELWPNDEAEKGFNLYLDSVLLVLPLLIMIFVYAIIVRTLCVGIRLENRSVAL

>XP_037505025.1 SKR type A-like SKR1A [Rhipicephalus sanguineus]

MESHNYSLDNSSSSNEEQPTATSWWHTEEAVLVAPYAIILLLAVLGNGLVILTLAVNQRMRTVTNLFLLNLAV
SDLLLGVFCMPFTLAGVILREFVFGQLMCKLIPYLQAVSVCVSDWTLVAVSVERYYAICEPLRSRAWQTPRHA
QRTVATVWTVSLLLMLPIALLSELRPIRDSSKMKCREDWGDLLYERLFTLFLDVLLLVLPLLIMVATYARIAA
TLRSAMQQQQTEAATGSPKQNGSPLAAVHPAQREASTSLRVHRGATIRWHQRDPDRSLATKQRIIRMLFVVVV
EFFVCWTPLYVLNTVSLFRPEAVYYGLGYRGICFLQLLAYASSCCNPITYCFMNRTFRVSFLALCRRRQRVQG
PGSVRRLTKQDFVYVLDREKT

>XP_037505292.1 SKR1B [Rhipicephalus sanguineus]
MVYVIAQQHHHTTSSQASPPVSVSVSVWTLVSISIERFFAIVRPLQSRRWQTVSHAYRVIAVIWALSLLHVAP
IAVLSQLIPTRTAGHQKCRELWPSDEAEKGFNLYLDSVLLVLPLLIMIVVYAIIVRTLCVGIRLENRSVALGI
PMKSASRGNENTNSSCSGAGSPTAREETTSAGTTTSTKKEALGRSSNGVRCQTLLVRGCNPEKTQASVVRVIR
MLFVVVVEFFVCWTPLYVVHTWSLYDPDAVYDWLGSAGVSVVHLLAYASSCSNPITYCFMHQKFRQGFLAAVG
CRRGGRPCCGSEGSCFAQQGSVRGSTYVSGTALVYQDAHKKGTGPFFESKSYT

>XP_065290468.1 SK type B receptor-like isoform X1 [Dermacentor
albipictus]
MDTYNYSFANSSSSKQEKPTATIWWQSEEAVLVAPYTIILLLAVLGNGLVILTLAINQRMRTVTNLFLLNLAV
SDLLLGVFCMPFTLVGVLLREFVFGQLMCKLIPYLQAVSVCVSDWTLVAVSVERYYAICQPLRSRAWQTPRHA
QRTVAVVWVVSLFLMLPIALLSELRPTRDSAKMKCREDWGDALYERLFTLFLDVLLLVLPLVVMVATYARIAV
TLHSAMQQQQTTATTGSPLKNGSPFAAVHPSQREPGTSLRLHRGATIRWHPRDPDRSLATKQRIIRMLFVVVV
EFFVCWTPLYVLNTVSLFQPEAVYYGLGYRGICFLQLLAYASSCCNPITYCFMNRTFRVSFLALCRRSCSQRK
QGPGNVRSLDKQDFVYVLGREVT

>XP_065288676.1 SKR type A-like [Dermacentor albipictus]
MVNRLMSSLPLLDLRQPLYNKTPSPPFTELTVPTVVLPGVALFTEFPDGVENSTSHPDSISAVDGLPDAADLI
VRVCLYAVIFVFAVVGNVLVLVTLVQNKRMRTVTNVFLVNLAVSDLLLGVFCMPFTLIGSILRNFVFGELMCK
FIPYLQAVSVSVSVWTLVSISLERFFAIVRPLQSRRWQTVSHAYRVIAVVWALSLVHVAPIAVLSQLIPTRTA
GRQKCRELWHSDEAEKGFNLYLDSVLLVLPLLIMIVVYAIITRTLCVGIRLENRSVALGIPMKSTVRENDNGS
SACSDSATPTGREEASFAMTTASTNKGSKFSRSANGVRCQTLLVRGCNTERAQASMVRVIRMLFVVVVEFFVC
WTPLYVVHTWSLYDPDTVYDWLGSAGVSVVHLLAYASSCSNPITYCFMHQKFRQGFLTAVGCRRGIRGRCGGA
ESCFSRQSSVRGATYASGTTLVYQDTKNKVPNGRGADFV

>XP_065283059.1 SKR type A-like isoform X1 [Dermacentor albipictus]
MCRGAESTNSSGDALSNETAAAEALASAAAGCWTGISFETAFRILLYAIIFVCAVVGNSLVIITLVQRRRMRT
VTNVFLLNLAVSDLLLGVFCMPTTLVGSVLRNFVLGAAMCKLIPYLQAVSVSVSAWTLVSISLERFFAIVRPL
ESRRWQTRSHAYKVILMVWLCSMVTMLPIAVLSQLVPLRGERKKCREVWPNVTSERVFNVYLDVTLFLLPLVI
MSFVYSCIGATLCHGMKLDNKSTLCCCSLVHRALARRCRGCAAPAPETEQHPSKDEELMSTAVVHEEYHTNPN
MPQLMAPPDVEARLQKMPRKQFIFRGNYRKSRASKRRVIRMLSVLVLEFFVCWTPLYVLHTWTVFDAHAAYSR
VPAGAFAAVHLLAYVSSCCNPITYCFMHDKYRQAFRNVLACSSASGRRRRRWSSTKSHDRFVSLNSICSNGKN
ASGKGSLRSTFTSRLSSLKEERATNDFNEAADQNHAVCKGPVTSLADGESSSSCNMSTYCGRRLDVANASDTE
S

>XP_054929511.1 SK type B receptor-like [Dermacentor andersoni]
MDTYNYSFDNSSSSKQEKPTAAIWWQSEEAVLVAPYTIILLLAVLGNGLVILTLAINQRMRTVTNLFLLNLAV
SDLLLGVFCMPFTLVGVLLREFVFGQLMCKLIPYLQAVSVCVSDWTLVAVSVERYYAICQPLRSRAWQTPRHA
QRTVAVVWAVSLLLMLPIALLSELRPTRDSAKMKCREDWGEALYERLFTLFLDVLLLVLPLVVMVATYARIAV
TLHSAMQQQQTTATTGSPLKNGSPFAAVHPSQPEPGTSLRLHRGATIRWHPRDPDRSLATKQRIIRMLFVVVV
EFFVCWTPLYVLNTVSLFQPEAVYYGLGYRGICFLQLLAYASSCCNPITYCFMNRTFRVSFLALCRRSCSQRK
QGPGNVRSLEKQDFVYVLGREVT

>XP_050039647.1 SKR type A-like [Dermacentor andersoni]
MVNRLMSSLPLLDLRQPLYNKTPSPPFTELAVPTVVLPDVAVFTEFPDGVENSTSHPDSISAVDGLPDAADLI
VRVCLYAVIFVFAVVGNVLVLVTLVQNKRMRTVTNVFLVNLAVSDLLLGVFCMPFTLIGSILRNFVFGELMCK
FIPYLQAVSVSVSVWTLVSISLERFFAIVRPLQSRRWQTVSHAYRVIAVVWALSLVHVAPIAVLSQLIPTRTA

GRQKCRELWHSDEAEKGFNLYLDSVLLVLPLLIMIVVYAIITRTLCVGIRLENRSVALGIPMKSAVRENDNGS
SACSDSATPTGREEASFAMTTASTNKGSKFSRSANGVRCQTLLVRGCNPERAQASMVRVIRMLFVVVVEFFVC
WTPLYVVHTWSLYDPDTVYDWLGSAGVSVVHLLAYASSCSNPITYCFMHQKFRQGFLTAVGCRRGIRGRCGGA
GSCFARQSSVRGATYASGTTLVYQDTKNKVPNGRGADFV

>XP_050040288.1 SKR type A-like isoform X1 [Dermacentor andersoni]
MCRGAELTNSSGDALSNETAAAEALASAAAGCWAGISFETAFRILLYAIIFVCAVVGNSLVIITLVQRRRMRT
VTNVFLLNLAVSDLLLGVFCMPTTLVGSVLRNFVLGAAMCKLIPYLQAVSVSVSAWTLVSISLERFFAIVRPL
ESRRWQTRSHAYKVILMVWLCSMVTMLPIAVLSQLVPLRGERKKCREVWPNVTSERVFNVYLDVTLFLLPLVI
MSFVYSCIGATLCHGMKIDNKSTLCCCSLVHRALARRCRGCAAPAPETEQHPSKDEELMSTAVVHEEYHTNPN
MPQLMAPPDVEARLQKMPRRQFIFRGNYRKSRASKRRVIRMLSVLVLEFFVCWTPLYVLHTWTVFDAHAAYSR
VPAGAFAAVHLLAYVSSCCNPITYCFMHDKYRQAFRNVLACSSASSRRRRRWSSTKSHDRFVSLNSICSNGKN
ASGKGSLRSTFTSRLSSLKEERATNDFNEVAGQNHAVCKGPVTSLADGESSSSCNMSTYCGRRLDVANPSDTE
S

>XP_049521298.1 SK type B receptor [Dermacentor silvarum]
MDSYNYSFDNSSSGNKTQPTAPTWWHSEEAVLVAPYTIILLLAVLGNGLVILTLAVNQRMRTVTNLFLLNLAV
SDLLLGVFCMPFTLAGVLLREFVFGQLMCKLIPYLQAVSVCVSDWTLVAVSVERYYAICEPLRSRAWQTPRHA
QRTVAAVWVVSLLLMLPIALLSELRPIRDSAKMKCREDWGELLYERLFTLFLDVLLLVLPLLIMVATYARIAA
TLRSAMQQQQTEAATGSPVKNGSPFAAVHPSQPEPGTSLRLHRGATIRWHQRDPDRSLATKQRIIRMLFVVVV
EFFVCWTPLYVLNTVSLFRPEAVYYGLGYRGICFLQLLAYASSCCNPITYCFMNRTFRVSFLALCRRSCSHRK
QGAGRVRSLNKQDFVYVLSREVT

>XP_049521496.1 SKR type A isoform X1 [Dermacentor silvarum]
MCRGAESANSSGDALSNETAAAEALAAAAAGCWTGISFETAFRILLYAIIFVCAVVGNSLVIITLVQRRRMRT
VTNVFLLNLAVSDLLLGVFCMPTTLVGSILRNFVLGGAMCKLIPYLQAVSVSVSAWTLVSISLERFFAIVRPL
ESRRWQTRSHAYKVILLVWLCSMVTMLPIAVLSQLVPLRGERKKCREVWPNVTSERVFNVYLDVTLFLLPLVI
MSFVYSCICATLCHGMKLDNKSTLCCCSLIHRALARRCRGCAAPTPETEQHPSKDEELMSTAVVHEEYHTNPN
MPQLIAPPDVEARLQKMPRRQFIFRGNYRKSRASKRRVIRMLSVLVLEFFVCWTPLYVLHTWTVFDAHAAYSR
VPAGAFAAVHLLAYVSSCCNPITYCFMHDKYRQAFRNVLACSSASSRRRRRWSSTKSHDRFVSLNSICSNGKN
VSGRGSLRSTFTSRLSSVKEERATNDVNEASDQKHDIRHNVKSPPATHDLCFERKTVLRKWTACAGIPAPVLT
FPVASLSWP

>XP_037567660.1 SKR [Dermacentor silvarum]
MAAVSVSVSVWTLVSISLERFFAIVRPLQSRRWQTISHAYRVIAVVWALSLVHVAPIAVLSQLIPTRTAGRQK
CRELWPSDEAEKVFNLYLDSVLLVLPLLIMIVVYAVITRTLCVGIRLENRSVALGIPMKSAAREMDKGSSSCS
GTATPTGREDASFAMTTASTNKGTTFGRSANGVRCHTVLVRGCNPERAQASMVRVIRMLFVVVVEFFVCWTPL
YVVHTWSLYDPDTVYDWLGSAGVSVVHLLAYASSCSNPITYCFMHQKFRKGFLAAVDCRRGGRGCCGGAASCF
NRQRSVRGSTYVSGTTFVYEDTQKKVPNGRGADFV

>KAH6935953.1 SKR [Hyalomma asiaticum]
MESHNYSLDNSSSKNNEEPSSWWQTEEAVLVAPYAIILLLAVLGNGLVILTLAVNQRMRTVTNLFLLNLAVSD
LLLGVFCMPFTLAGVLLREFVFGQLICSCFSTFRCNLCAYECAAVSVCVSDWTLVAVSVERYYAICQPLRSRA
WQTPRHAQRTVATVWTVSLIIMLPIALLSELKPIRDTTKMKCREDWGDLLYERLFTLFLDVLLLVLPLLIMVT
TYARIAATLHSAMQQQQTETATGSPNKNGSPFASVHPSQPEPGTSLRLHRGATIRWHQRDPDRSLATKQRIIR
MLFVVVVEFFVCWTPLYVLNTVSLFRPEAVYYGLGYRGICFLQLLAYASSCCNPITYCFMNRTFRVSFLALCR
RGSAQRKSGPGSVRSFTKQDFVYVLGRERM

>XP_022671206.1 SKR type A-like isoform X1 SKR [Varroa destructor]
MNESGSMLMMDVIGGAGIGAKDDSIIDFTGSLGVSVSAGVGERVSSSGGSSLAAEAVAMASGGTLGDLATDAE
TALDVMVTGSASMGNGVSVILVSSVSARQTSIFDSSALLILSYGAIFATGLVGNLLVILTLTRNQRGKVATNV
FLLNLAVSDLLLGVFCMPVTLSGVLLRNFVFGEIMCKVIPYLQGVTVAVNAWTLVMISIERYYAVCEPLKSRG
WYASSHAWHCVFVIWVFALTAMLPVFLLNRLQPTGKGRQKCREAWPEPTVETAFTVMLDLLLLVIPLAVMAFA
YINITVTLRKGIRDNERSSNEIKIMGPRGDIGVMSHKSNEGMIEREPRLIVKWCRVAAPEESSESLPGHGVKI

PPAEPTATQPLPTVATHDPDSSPVVDAKSPTAQCPVSASSTHREEIIRYLRTNNHEQRMAVRRKVIRMMFVVV
LEFFVCWTPIYVVNTIASFSKDLLTPLGGVGISLLHLLSYVSSCCNPITYCFMNKNFRREFKRSIRCCDPKYL
SKCSSFRSPRSRGRDIRSTPSRRGARIELPRQRRRAICASRRINNSLTDL

>XP_018494335.2 SKR type A SKR1A [Galendromus occidentalis]
MAAVNHNRSFTSALIDPSSAVSTPEVVTEWLGVLTTYLLILSYISIFILGLIGNVLVIVTLTRNKRFKVPTNA
FLLNLAVSDLLVGVFCMPFTLSGIVFREFIFGDLMCKAVPYVQAVAVAVSAWTLVMISFERYFAVCQPLRCRT
WQSSWHAVKLIIVIWIVALILMLPLGCLNRLIEMERFKGKYKCREKWPSPALETAFNLALDAVLFAVPLAIMG
TAYVQISWTLKKGIKDHERCVNAIPITRTSGLMIKDRSCDDEGGGKQNGKKVADKEPMLIVKWCRMTENNSDY
EEDNVDEAPAEREVNGGDSLRMSRTVSTHHRRELLIGHLRSNNLEQRMAVKRKVIRMMFVVVLEFFICWTPLY
TINTISSISPEYLSPLGSFGPPLFLLLSYISSCCNPITYCFMNNNFRREFRRAVGCNSKRGYLGSQAARTRSP
RARLRDVQRSSPHEEDLSRNLSRRELRIDESLIITREAAVDERKIHQISSSQESGNVAFGRLPGTTRTR

>XP_028966986.1 SKR type A SKR1B [Galendromus occidentalis]
MSLLSTMRNVSHLCFVAPLDAAGIYEIRDLESYTCQQLSVTSDDGDYGSIEFNMNGTEVEGDAASTLNAEVSV
ILVQSRDSWQFNYSALLILSYGAIFATGLVGNSLVILTLTRNQRGKVATNVFLLNLAISDLLLGVFCMPFTLA
GFLLRNFIFGQLMCKVIPFLQGVTVAVNAWTLVMISIERYFAVCEPLKSRGFYASTHAWHSVVIIWIFALTSM
TPIFYVNRLQPAGYGRQKCREDWPNPTVESAFTVLLDLLLLVIPLTVMAFAYINITVTLRKGIRENEKGSNEI
KLMGPRGIGVVNNKTPENVIEREPRLIVKWCKVQNEDSSDSSFGHVDRPKAANPASSDSSSHSTHDNFSCNDF
RAPVQFTNYTPTPREEMIRYLRTNNNEQRMAVRRKVIRMVFVVVLEFFVCWTPIYAINTIASFSKDLLTPLGG
VGISLFHLLSYVSSCCNPVTYCFMNKNFCREFKRSISCCDPKYLKFWILRSRERGPEISLTGIRAQTRIEIRD
NRRAFCSQRNHTDL

>XP_028967749.1 SKR1C [Galendromus occidentalis]_truncated
MVMDAANSSDPLVQIQGPSRLSIPWTTGIIVAAYGSIFVLGLIGNSLVILTLTRNKRMKVATNVFLLNLAVSD
LLLGVFCMPFTLSGALLRNFIFGELMCKLIPFLQGVTVAVSVWTLVVISVERYLAVCSPLSQHGSSFASRHAL
LITLIIWIVALATMSPTFLLSELRPVGETGRQKCREMWTDKYKEYAFNIGLDLMLFALPMIAMSFAYGNICWM
LKRGLTEEKPIHMNHMSEIVIAGTSGCAVLSDKTSQEVIDREPRLIVKWSRVEENSEPLATTELTNQKWPTTA
LNGSEVSSSAKEMIRFMRTNTYEHRMAVRNRVIRMMFVVVIEFFICWTPIYVINTVNSISPAHLSPLGSRGVS
LFHLLAYCSSSCCNPITYCFMNRNFRREFRRAVRCNEF

>OQR80232.1 SKR type A-like SKR partial [Tropilaelaps mercedesae]
MPVTLSGVLLRSFVFGELMCKVIPYLQGVTVAVNAWTLVMISIERYYAVCEPLKSRGWYASSHAWHCVFVIWA
FALTAMLPIFLLNRLQPTGKGRQKCREAWPDPTVETVFTVMLDLLLLVIPLAVMAFAYINITITLRKGIRHNE
RAANEIKIMGPRGGIGVVSHKTDESMIEREPRLIVKWRRVVPPEDSSDSSLGHGLKLPPTQPTATQPLPTAAV
HGPDTSPAVKSPTSQCPVGATSVHREEIIRYLRTNNHEQRMAVRRKVIRMMFVVVLEFFVCWTPIYVINTIAS
FSKDLLTPLGGVGISLFHLLSYISSCCNPITYCFMNKNFR

>NP_001097023 Drosulfakinin receptor at 17D3 [Drosophila melanogaster]
MFNYEEGDADQAAMAAAAAYRALLDYYANAPSAAGHIVSLNVAPYNGTGNGGTVSLAGNATSSYGDDDRDGYM
DTEPSDLVTELAFSLGTSSSPSPSSTPASSSSTSTGMPVWLIPSYSMILLFAVLGNLLVISTLVQNRRMRTIT
NVFLLNLAISDMLLGVLCMPVTLVGTLLRNFIFGEFLCKLFQFSQAASVAVSSWTLVAISCERYYAICHPLRS
RSWQTISHAYKIIGFIWLGGILCMTPIAVFSQLIPTSRPGYCKCREFWPDQGYELFYNILLDFLLLVLPLLVL
CVAYILITRTLYVGMAKDSGRILQQSLPVSATTAGGSAPNPGTSSSSNCILVLTATAVYNENSNNNNGNSEGS
AGGGSTNMATTTLTTRPTAPTVITTTTTTTVTLAKTSSPSIRVHDAALRRSNEAKTLESKKRVVKMLFVLVLE
FFICWTPLYVINTMVMLIGPVVYEYVDYTAISFLQLLAYSSSCCNPITYCFMNASFRRAFVDTFKGLPWRRGA
GASGGVGGAAGGGLSASQAGAGPGAYASANTNISLNPGLAMGMGTWRSRSRHEFLNAVVTTNSAAAAVNSPQL

> NP_001097021 Drosulfakinin receptor at 17D1 [Drosophila melanogaster]
MLPRLCADACRQCFAKIARRDTHRGTRTPYGCADTQSRPKPNFLLREVDEVCCTAASASPRLLVLFRDHKRAS
FFGLTIDAFYHYLRQALPLAKEAAIHLNASNEISAVGDGVTITGTPGDLLNYSGLELDLGLDLDLNLDMDLAT
TPSSSTLAPAVTVRTPGNRSVVRVSADVPIWVVPCYSAILLCAVVGNLLVVLTLVQNRRMRTITNVFLLNLAI
SDILLGVFCMPVTLVGTLLRHFIFGELLCKLIQFAQAASVAVSSWTLVAISCERYYAICHPLRSRTWQTINHA
NKIIAIIWLGSLVCMTPIAAFSQLMPTSRPGLRKCREQWPADSLNYERAYNLFLDLALLVLPLLALSFTYLFI

TRTLYVSMRNERAMNFGSSGPEVTTSSSAAVAEAGSQRRANGSHCQSLDTIVPHQHNPHQQHHHHSQYYYDYG
HCGSKRRLISGGGPCEGRRHLYCMRSASVKSLRHQQINGGGGTLSGTGAGNGECCSRVHRMRQQMQLQQQGYV
SDNESRRKSLSQPSLRITEAGLRRSNETKSLESKKRVVKMLFVLVLEFFICWTPLYVINTMTMLLGPTVYEYV
GYTSISFLQLLAYSSSCCNPITYCFMNASFRRAFVDTFKGMRVCERLCAPCCFWRRRSKNETNLSVAGNSIAL
ANSVMSSHTILESPRL

>P32238 Cholecystokinin receptor type A [Homo sapiens]
MDVVDSLLVNGSNITPPCELGLENETLFCLDQPRPSKEWQPAVQILLYSLIFLLSVLGNTLVITVLIRNKRMR
TVTNIFLLSLAVSDLMLCLFCMPFNLIPNLLKDFIFGSAVCKTTTYFMGTSVSVSTFNLVAISLERYGAICKP
LQSRVWQTKSHALKVIAATWCLSFTIMTPYPIYSNLVPFTKNNNQTANMCRFLLPNDVMQQSWHTFLLLILFL
IPGIVMMVAYGLISLELYQGIKFEASQKKSAKERKPSTTSSGKYEDSDGCYLQKTRPPRKLELRQLSTGSSSR
ANRIRSNSSAANLMAKKRVIRMLIVIVVLFFLCWMPIFSANAWRAYDTASAERRLSGTPISFILLLSYTSSCV
NPIIYCFMNKRFRLGFMATFPCCPNPGPPGARGEVGEEEEGGTTGASLSRFSYSHMSASVPPQ

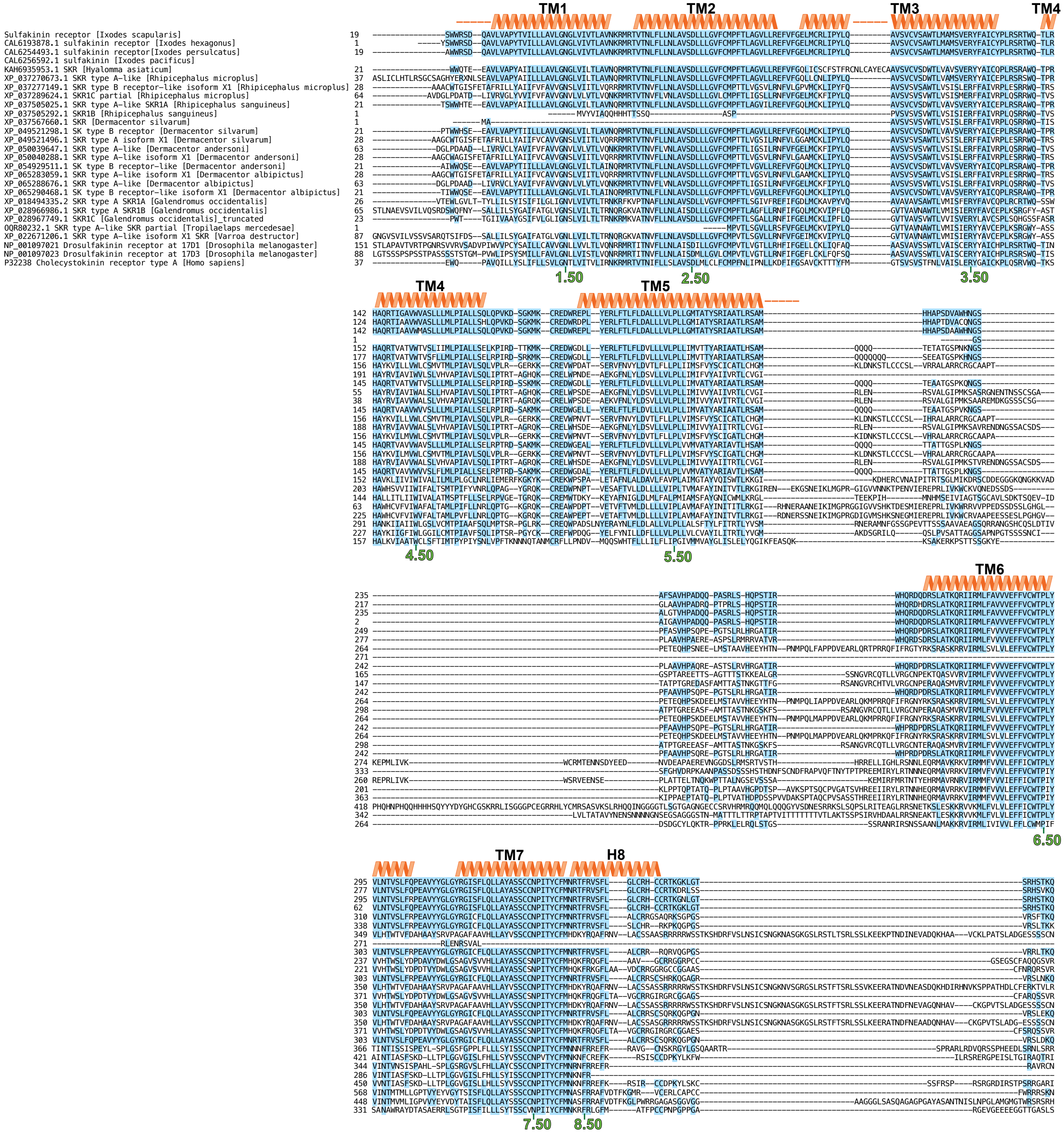
TM1
TM2
TM3
TM4
TM5
TM6
TM7
H8
Sulfakinin receptor [Ixodes scapularis]
CAL6193878.1 sulfakinin receptor [Ixodes hexagonus]
CAL6254493.1 sulfakinin receptor[Ixodes persulcatus]
CAL6256592.1 sulfakinin [Ixodes pacificus]
KAH6935953.1 SKR [Hyalomma asiaticum]
XP_037270673.1 SKR type A-like [Rhipicephalus microplus]
XP_037277149.1 SKR type B receptor-like isoform X1 [Rhipicephalus microplus]
XP_037289624.1 SKR1C partial [Rhipicephalus microplus]
XP_037505025.1 SKR type A-like SKR1A [Rhipicephalus sanguineus]
XP_037505292.1 SKR1B [Rhipicephalus sanguineus]
XP_037567660.1 SKR [Dermacentor silvarum]
XP_049521298.1 SK type B receptor [Dermacentor silvarum]
XP_049521496.1 SKR type A isoform X1 [Dermacentor silvarum]
XP_050039647.1 SKR type A-like [Dermacentor andersoni]
XP_050040288.1 SKR type A-like isoform X1 [Dermacentor andersoni]
XP_054929511.1 SK type B receptor-like [Dermacentor andersoni]
XP_065283059.1 SKR type A-like isoform X1 [Dermacentor albipictus]
XP_065288676.1 SKR type A-like [Dermacentor albipictus]
XP_065290468.1 SK type B receptor-like isoform X1 [Dermacentor albipictus]
XP_018494335.2 SKR type A SKR1A [Galendromus occidentalis]
XP_028966986.1 SKR type A SKR1B [Galendromus occidentalis]
XP_028967749.1 SKR1C [Galendromus occidentalis]_truncated
OQR80232.1 SKR type A-like SKR partial [Tropilaelaps mercedesae]
XP_022671206.1 SKR type A-like isoform X1 SKR [Varroa destructor]
NP_001097021 Drosulfakinin receptor at 17D1 [Drosophila melanogaster]
NP_001097023 Drosulfakinin receptor at 17D3 [Drosophila melanogaster]
P32238 Cholecystokinin receptor type A [Homo sapiens]
1.50
2.50
3.50
4.50
5.50
6.50
7.50
8.50

### Percent Identity Matrix Sulfakinin Receptors

| Species | Accesion Number | XP_042145535 | CAL6254493 | CAL6193878 | CAL6256592 | XP_037270673 | XP_037277149 | XP_037289624 | XP_037505025 | XP_037505292 | XP_065290468 | XP_065288676 | XP_065283059 | XP_054929511 | XP_050039647 | XP_050040288 | XP_049521298 | XP_049521496 | XP_037567660 | KAH6935953 | XP_022671206 | XP_018494335 | XP_028966986 | XP_028967749 | OQR80232 | NP_001097023 | NP_001097021 | P32238 | P32239 |
|---|---|---|---|---|---|---|---|---|---|---|---|---|---|---|---|---|---|---|---|---|---|---|---|---|---|---|---|---|---|
| *I. scapularis* | XP_042145535 | 100 | 97.1 | 93.6 | 98.0 | 72.6 | 46.9 | 56.2 | 74.6 | 44.7 | 73.9 | 51.6 | 47.2 | 74.2 | 51.8 | 47.2 | 76.1 | 47.2 | 46.9 | 71.9 | 45.7 | 43.5 | 42.1 | 45.2 | 49.6 | 44.6 | 48.4 | 36.4 | 36.3 |
| *I. persulcatus* | CAL6254493 | 97.1 | 100 | 93.1 | 97.3 | 73.1 | 47.2 | 56.2 | 74.9 | 45.1 | 73.1 | 51.8 | 47.2 | 73.7 | 52.1 | 47.2 | 75.8 | 47.2 | 47.3 | 71.9 | 45.4 | 43.5 | 42.1 | 45.2 | 49.6 | 44.3 | 48.1 | 36.9 | 36.5 |
| *I. hexagonus* | CAL6193878 | 93.6 | 93.1 | 100 | 89.7 | 76.3 | 48.7 | 60.1 | 78.5 | 45.1 | 77.2 | 53.7 | 48.7 | 76.7 | 53.7 | 48.7 | 78.3 | 48.7 | 47.3 | 74.4 | 46.5 | 44.6 | 43.8 | 47.3 | 49.6 | 45.6 | 50.6 | 37.3 | 37.2 |
| *I. pacificus* | CAL6256592 | 98.0 | 97.3 | 89.7 | 100 | 71.5 | 38.0 |  | 75.0 | 46.6 | 75.3 | 45.8 | 38.0 | 75.3 | 46.6 | 38.0 | 74.7 | 38.0 | 45.8 | 74.7 | 40.2 | 36.5 | 33.9 | 44.9 | 46.2 | 34.7 | 42.0 | 30.7 | 28.6 |
| *R. microplus* | XP_037270673 | 72.6 | 73.1 | 76.3 | 71.5 | 100 | 45.6 | 49.8 | 86.7 | 44.1 | 80.2 | 47.8 | 45.6 | 80.0 | 48.0 | 45.6 | 83.3 | 45.9 | 45.8 | 81.9 | 42.7 | 42.6 | 40.5 | 45.7 | 47.9 | 43.5 | 46.8 | 36.4 | 36.0 |
| *R. microplus* | XP_037277149 | 46.9 | 47.2 | 48.7 | 38.0 | 45.6 | 100 | 60.9 | 47.4 | 42.9 | 46.6 | 50.7 | 92.9 | 46.8 | 50.5 | 92.7 | 47.1 | 89.0 | 47.0 | 45.7 | 38.3 | 35.9 | 37.4 | 41.6 | 42.1 | 41.8 | 42.5 | 36.2 | 36.2 |
| *R. microplus* | XP_037289624 | 56.2 | 56.2 | 60.1 |  | 49.8 | 60.9 | 100 | 53.6 | 83.3 | 54.1 | 86.6 | 61.4 | 53.6 | 87.7 | 60.9 | 53.6 | 61.4 | 90.6 | 50.5 | 44.5 | 46.5 | 42.4 | 46.4 | 53.7 | 46.8 | 48.2 | 41.5 | 41.4 |
| *R. sanguineus* | XP_037505025 | 74.6 | 74.9 | 78.5 | 75.0 | 86.7 | 47.4 | 53.6 | 100 | 46.0 | 88.6 | 51.5 | 47.4 | 88.3 | 52.1 | 47.1 | 91.7 | 47.7 | 48.3 | 89.6 | 45.8 | 44.6 | 44.3 | 46.6 | 50.4 | 45.5 | 48.4 | 37.8 | 37.0 |
| *R. sanguineus* | XP_037505292 | 44.7 | 45.1 | 45.1 | 46.6 | 44.1 | 42.9 | 83.3 | 46.0 | 100 | 45.4 | 77.9 | 43.2 | 46.1 | 79.1 | 42.9 | 45.7 | 43.5 | 81.7 | 46.4 | 37.5 | 36.1 | 34.5 | 37.5 | 42.1 | 38.0 | 37.3 | 33.7 | 33.2 |
| *D. albipictus* | XP_065290468 | 73.9 | 73.1 | 77.2 | 75.3 | 80.2 | 46.6 | 54.1 | 88.6 | 45.4 | 100 | 50.7 | 46.6 | 98.2 | 51.3 | 46.6 | 91.5 | 46.8 | 47.9 | 86.3 | 46.4 | 44.3 | 43.4 | 46.9 | 50.8 | 45.2 | 48.7 | 37.0 | 36.8 |
| *D. albipictus* | XP_065288676 | 51.6 | 51.8 | 53.7 | 45.8 | 47.8 | 50.7 | 86.6 | 51.5 | 77.9 | 50.7 | 100 | 50.7 | 51.0 | 98.5 | 50.5 | 50.4 | 51.4 | 89.6 | 49.3 | 39.8 | 39.4 | 38.1 | 44.9 | 47.0 | 40.1 | 41.7 | 35.7 | 36.5 |
| *D. albipictus* | XP_065283059 | 47.2 | 47.2 | 48.7 | 38.0 | 45.6 | 92.9 | 61.4 | 47.4 | 43.2 | 46.6 | 50.7 | 100 | 47.1 | 50.5 | 98.4 | 47.4 | 91.6 | 47.0 | 45.7 | 38.9 | 36.2 | 38.0 | 40.5 | 42.1 | 41.6 | 42.6 | 36.2 | 36.2 |
| *D. andersoni* | XP_054929511 | 74.2 | 73.7 | 76.7 | 75.3 | 80.0 | 46.8 | 53.6 | 88.3 | 46.1 | 98.2 | 51.0 | 47.1 | 100 | 51.5 | 47.1 | 92.3 | 47.4 | 48.3 | 86.5 | 46.4 | 44.6 | 44.0 | 46.9 | 50.8 | 45.7 | 48.4 | 37.3 | 36.5 |
| *D. andersoni* | XP_050039647 | 51.8 | 52.1 | 53.7 | 46.6 | 48.0 | 50.5 | 87.7 | 52.1 | 79.1 | 51.3 | 98.5 | 50.5 | 51.5 | 100 | 50.2 | 51.0 | 51.2 | 90.2 | 49.9 | 40.0 | 39.4 | 38.1 | 44.9 | 47.0 | 40.1 | 41.7 | 35.7 | 36.5 |
| *D. andersoni* | XP_050040288 | 47.2 | 47.2 | 48.7 | 38.0 | 45.6 | 92.7 | 60.9 | 47.1 | 42.9 | 46.6 | 50.5 | 98.4 | 47.1 | 50.2 | 100 | 47.4 | 91.0 | 46.6 | 45.4 | 38.9 | 36.0 | 37.8 | 40.5 | 42.1 | 41.3 | 42.6 | 36.2 | 36.2 |
| *D. silvarum* | XP_049521298 | 76.1 | 75.8 | 78.3 | 74.7 | 83.3 | 47.1 | 53.6 | 91.7 | 45.7 | 91.5 | 50.4 | 47.4 | 92.3 | 51.0 | 47.4 | 100 | 47.7 | 47.9 | 88.1 | 46.7 | 44.9 | 44.0 | 46.6 | 50.0 | 45.0 | 48.2 | 37.9 | 36.8 |
| *D. silvarum* | XP_049521496 | 47.2 | 47.2 | 48.7 | 38.0 | 45.9 | 89.0 | 61.4 | 47.7 | 43.5 | 46.8 | 51.4 | 91.6 | 47.4 | 51.2 | 91.0 | 47.7 | 100 | 47.6 | 45.7 | 39.0 | 36.6 | 38.2 | 40.8 | 42.5 | 41.6 | 42.6 | 36.2 | 36.2 |
| *D. silvarum* | XP_037567660 | 46.9 | 47.3 | 47.3 | 45.8 | 45.8 | 47.0 | 90.6 | 48.3 | 81.7 | 47.9 | 89.6 | 47.0 | 48.3 | 90.2 | 46.6 | 47.9 | 47.6 | 100 | 49.1 | 40.0 | 37.9 | 36.4 | 39.5 | 44.0 | 40.7 | 40.5 | 33.9 | 34.4 |
| *H. asiaticum* | KAH6935953 | 71.9 | 71.9 | 74.4 | 74.7 | 81.9 | 45.7 | 50.5 | 89.6 | 46.4 | 86.3 | 49.3 | 45.7 | 86.5 | 49.9 | 45.4 | 88.1 | 45.7 | 49.1 | 100 | 43.9 | 43.1 | 42.0 | 45.3 | 46.9 | 45.0 | 47.6 | 38.6 | 37.3 |
| *V. destructor* | XP_022671206 | 45.7 | 45.4 | 46.5 | 40.2 | 42.7 | 38.3 | 44.5 | 45.8 | 37.5 | 46.4 | 39.8 | 38.9 | 46.4 | 40.0 | 38.9 | 46.7 | 39.0 | 40.0 | 43.9 | 100 | 48.5 | 65.3 | 56.1 | 88.6 | 35.7 | 34.5 | 33.4 | 33.6 |
| *G. occidentalis* | XP_018494335 | 43.5 | 43.5 | 44.6 | 36.5 | 42.6 | 35.9 | 46.5 | 44.6 | 36.1 | 44.3 | 39.4 | 36.2 | 44.6 | 39.4 | 36.0 | 44.9 | 36.6 | 37.9 | 43.1 | 48.5 | 100 | 47.9 | 53.3 | 53.4 | 35.5 | 33.5 | 33.9 | 34.3 |
| *G. occidentalis* | XP_028966986 | 42.1 | 42.1 | 43.8 | 33.9 | 40.5 | 37.4 | 42.4 | 44.3 | 34.5 | 43.4 | 38.1 | 38.0 | 44.0 | 38.1 | 37.8 | 44.0 | 38.2 | 36.4 | 42.0 | 65.3 | 47.9 | 100 | 55.5 | 74.2 | 35.8 | 34.2 | 33.0 | 33.3 |
| *G. occidentalis* | XP_028967749 | 45.2 | 45.2 | 47.3 | 44.9 | 45.7 | 41.6 | 46.4 | 46.6 | 37.5 | 46.9 | 44.9 | 40.5 | 46.9 | 44.9 | 40.5 | 46.6 | 40.8 | 39.5 | 45.3 | 56.1 | 53.3 | 55.5 | 100 | 56.6 | 40.3 | 39.3 | 35.9 | 36.8 |
| *T. mercedesae* | OQR80232 | 49.6 | 49.6 | 49.6 | 46.2 | 47.9 | 42.1 | 53.7 | 50.4 | 42.1 | 50.8 | 47.0 | 42.1 | 50.8 | 47.0 | 42.1 | 50.0 | 42.5 | 44.0 | 46.9 | 88.6 | 53.4 | 74.2 | 56.6 | 100 | 40.9 | 36.5 | 34.8 | 35.7 |
| *D. melanogaster 17D3* | NP_001097023 | 44.6 | 44.3 | 45.6 | 34.7 | 43.5 | 41.8 | 46.8 | 45.5 | 38.0 | 45.2 | 40.1 | 41.6 | 45.7 | 40.1 | 41.3 | 45.0 | 41.6 | 40.7 | 45.0 | 35.7 | 35.5 | 35.8 | 40.3 | 40.9 | 100 | 53.8 | 36.3 | 35.0 |
| *D. melanogaster 17D1* | NP_001097021 | 48.4 | 48.1 | 50.6 | 42.0 | 46.8 | 42.5 | 48.2 | 48.4 | 37.3 | 48.7 | 41.7 | 42.6 | 48.4 | 41.7 | 42.6 | 48.2 | 42.6 | 40.5 | 47.6 | 34.5 | 33.5 | 34.2 | 39.3 | 36.5 | 53.8 | 100 | 35.7 | 35.9 |
| *H. sapiens CCK1R* | P32238 | 36.4 | 36.9 | 37.3 | 30.7 | 36.4 | 36.2 | 41.5 | 37.8 | 33.7 | 37.0 | 35.7 | 36.2 | 37.3 | 35.7 | 36.2 | 37.9 | 36.2 | 33.9 | 38.6 | 33.4 | 33.9 | 33.0 | 35.9 | 34.8 | 36.3 | 35.7 | 100 | 50.5 |
| *H. sapiens CCK2R* | P32239 | 36.3 | 36.5 | 37.2 | 28.6 | 36.0 | 36.2 | 41.4 | 37.0 | 33.2 | 36.8 | 36.5 | 36.2 | 36.5 | 36.5 | 36.2 | 36.8 | 36.2 | 34.4 | 37.3 | 33.6 | 34.3 | 33.3 | 36.8 | 35.7 | 35.0 | 35.9 | 50.5 | 100 |

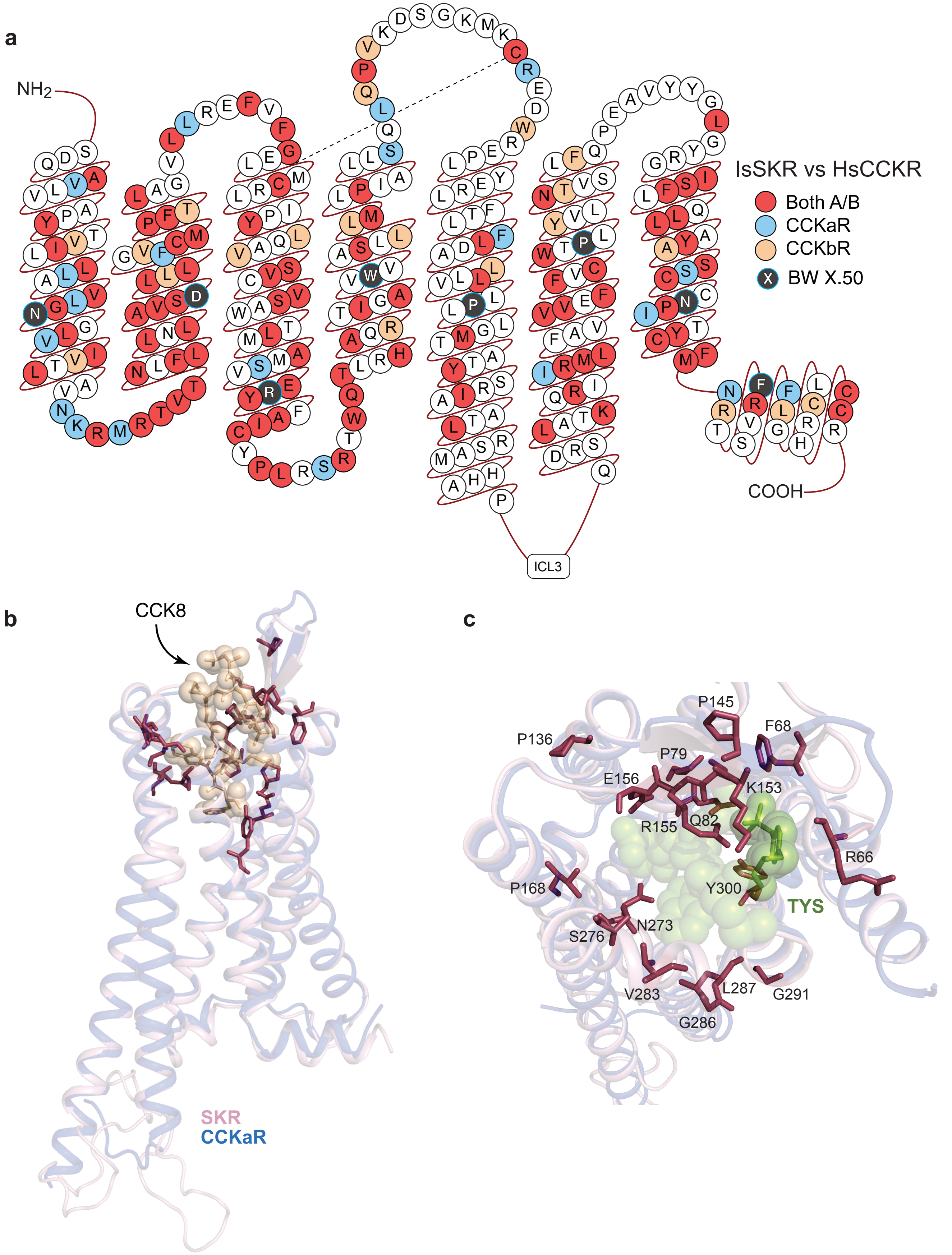
a
NH2
COOH
ICL3
IsSKR vs HsCCKR
Both A/B
CCKaR
CCKbR
BW X.50
b
CCK8
SKR
CCKaR
c
P145
F68
P136
P79
E156
K153
Q82
R155
R66
P168
Y300
TYS
N273
S276
L287
V283
G291
G286

**Prepro-CCHa peptide sequences used in alignments**

>XP_029834161.2 uncharacterized protein LOC8039983 isoform X1 [Ixodes scapularis]
MCFKPAMPFLLIMLALLVACTQCAAFGDSMEEGLRDKKIFTILRRNNSCKMYGHSCLGGHGKRS
EEGTVSSDVMQMLRKSAAEDDAGAMLAQARLDRPALLEKTLLQILRNRLQ

>XP_037279305.1 uncharacterized protein LOC119172349 [Rhipicephalus microplus]
MCFQSSLSVLLLCLALLVASAYSAAFGDSVEEGVRDKKIITLLRRNNSCKLYGHSCLGGHGKRS
DDGSAPGGAISPDVMQMLRKTSDDEALGPFPPQARFDRVALLEKALIQLLRNRMMV

>XP_037517515.1 uncharacterized protein LOC119394298 isoform X3 [Rhipicephalus sanguineus]
MCLQSSLSIWLLCFAVLVASAYSAAFGDSVEEGVRDKKIITLLRRNNSCKLYGHSCLGGHGKRS
DDGSAPGGAISPDVMQMLRKTSDDESLGPFPPQTRFDRVGLLEKALIQLLRNKMMV

>XP_037572707.1 uncharacterized protein LOC119455382 isoform X3 [Dermacentor silvarum]
MTRGFGDCLKGTAERRLEDIRQSAAKSASSLSMCLQSSMSVWLLCLAVLVASAYSAAFGDSVEE
GVRDKKIITLLRRNNSCKLYGHSCLGGHGKRSDDGSAPGGGVSPDVMQMLRKTTDDEALPPFPA
QARFDRVALLEKALIQLLRNRMMV

>XP_003748328.1 uncharacterized protein LOC100901943 [Galendromus occidentalis]
MKASLRLAMRSSVLLVLVAFAFVLSVAEQVSAADAATSFGNDVNRDFNTKRIIALLRRSSRQPN
ALADSCRLYGHSCLGGHGKRSSPVTAEEVDDDGPMYRIDYDWLQSRI

>XP_022706958.1 uncharacterized protein LOC111270805 [Varroa jacobsoni]
MRPFASNTFIVLFIAAVFIISFMGETTAADSGATYGNEAPRDFNTKRIIALLRRSSRQPNALAD
SCRLYGHSCLGGHGKRSSLNGAEEAEDEHAAYRFDDEWLRNRV

>OQR68378.1 hypothetical protein BIW11_12951 [Tropilaelaps mercedesae]
MRPFALKIFLVFFVAAVCIMSLIGQITAADPGASYGNDASRDFNTKRIIALLRRNSRQPNALAD
SCRFYGHSCLGGHGKRSSPNGAEEVDDEKATLRFDDDWLPNRL

>GBM21433.1 hypothetical protein AVEN_161049-1 [Araneus ventricosus]
MPTFSSGFPAETVEQISLSYICSPCFAWQMMFEHPVLSFNAFVFSCRTAGRAGDMRIYKCLQGF
PRRRLDRPSTQTVGFGAFPKMTMADNRLLLLIAAIILIIANFETVQAGGSCQSYGHSCLGGHGK
RNSEPPYALHLLLQQALRPRNEFPLTDYRTADRDAYDFLRGFSDSIKALEDKAN

>KAF8785304.1 hypothetical protein HNY73_010864 [Argiope bruennichi]

MVNRLVVFAAAIILIVANFENVVAGGSCQSYGHSCLGGHGKRNGEPPFGLHLLLQQALRARNEL
PGNAVAEMNYPAVADRTAYDFMRGFTNGVKALEEKAK

>KFM81698.1 hypothetical protein X975_10367, partial
[Stegodyphus mimosarum]
MTFNRVSPLTVAAILTFAYFETVSAGGSCQSYGHSCLGGHGKRNNEQLSAVQLLLQQALKGRDQ
YPLYDYRLPERDTYDLIRSFTDLKGLEDKTINKNE

>XP_035217372.1 neuropeptide CCHamide-2-like isoform X1
[Stegodyphus dumicola]
MTSNRVCPLALAAILTFAYFETVSAGGSCQSYGHSCLGGHGKRNNEQLSALQLLLQQALKGRDQ
YPLYDYRLPERDTYDILRSFTDLKVLEDKTVNKNE

>XP_042897003.1 uncharacterized protein LOC110283395
[Parasteatoda tepidariorum]
MSFLCRIMLFTAATFLIITCFGLTPAGGSCQSYGHSCLGGHGKRSSAGSVPLPLLWQQILRARE
PVPFVDYRIPERDAFDILKGFTDNIKALEEKVDGQN

>XP_035214177.1 uncharacterized protein LOC118187970
[Stegodyphus dumicola]
MSSTTIVSFALVLVILGIHVMHACPLSTGLCKQYGHSCLGGHGKRAEVQPSRLMKHIFREMSHL
PMKEADMLPQDFAESADFWENIFNNKADAFENDEQKELRANLFD

>GBM84837.1 hypothetical protein AVEN_95666-1 [Araneus
ventricosus]
MSTPTVVAFAFGMVILGIHVLHACPLSTSGKCKQYGHACLGGHGKRSDTSNQGLMDYILRKMAR
MQDENADTPTPRRDEWQSIFEKNYEKKLDDWDSYPEDGLRSTIYNAFN

>XP_022238232.1 uncharacterized protein LOC111085154 [Limulus
polyphemus]
MTEGRQKLWVRLALIYIIVCEVGGGCINYGHSCLGGHGKRGINGIFHETGDLKKLISQLLQNDR
TSFKYPGFNSIGRAEGRLPLNNLYSSAKNKQQYDSPPEYNFASSSSEDRDYPKDDHHRLNPRYP
GEDSGADDRFYSTDSRLYNLIISAVRDVN

>NP_001097784.1 CCHamide-1 [Drosophila melanogaster]
MWYSKCSWTLVVLVALFALVTGSCLEYGHSCWGAHGKRSGGKAVIDAKQHPLPNSYGLDSVVEQ
LYNNNNNNQNNQDDDNNDDDSNRNTNANSANNIPLAAPAIISRRESEDRRIGGLKWAQLMRQHR
YQLRQLQDQQQQGRGRGGQGQYDAAAESWRKLQQALQAQIDADNENYSGYELTK

>NP_001189216.1 CCHamide-2, isoform B [Drosophila melanogaster]
MKSTISLLLVVICTVVLAAQQSQAKKGCQAYGHVCYGGHGKRSLSPGSGSGTGVGGGMGEAASG
GQEPDYVRPNGLLPMMAPNEQVPLEGDFNDYPARQVLYKIMKSWFNRPRRPASRLGELDYPLAN
SAELNGVN

>NP_001280509.1 hypothetical protein precursor [Tribolium
castaneum]

MNCWSTQVVLLAFVMAFVLAAAEAKRGCATFGHSCYGGMGKRTENNNEELLQDVQSEENPAFVF
TGPRSENQQKLTPEQYDNISRVIRQWDTIVQESSGNAPRL

**CCHa receptor (CCHaR) amino acid sequences used in alignment**

```
>XP_042147864.1 CCHamide-1 Receptor CAK [Ixodes scapularis]
MSSLVVNATLATLRANRDPSASRAGINTSQNSHVNLTTGYGDSEEFVPYEQRLETYVVPTLFAL
IFIVGLLGNGTLILVFIRNRTMRSVPNIYIMSLSIGDFIVIAGTVPFISTIYVLDSWPYGLFLC
KLSEFLRDVSIGVTVLTLTVLSIDRYVAIAMPLLNHKGRRHTRRTVTILLAISVWIVAILMAIP
GTHYSFVMQVQATPNLHYSVCYPFPPEMWPWYPKLMVLLKFLIQYAIPLVIIGTFYCLMARQLI
RTSRAHLSQTSCGGVAHLKQMKARVKVAKIALAFVLLFAVCFFPNHVFMMWYYFAPNAPSQYNS
FWHVWKIMGYVMTFVNSCLNPVALYLVSGVFRNHFKHYLFCGRHPNVHARHNSYSFRTIHSSSM
SKCTASTKI

>CAL6223886.1 CCHamide-1 Receptor [Ixodes persulcatus]
MSSLVANATLATLRANRDPSASRADINTSQTSHVNLTTGYGDSEEFVPYEQRLETYVVPTLFAL
IFIVGLLGNGTLILVFIRNRTMRSVPNIYIMSLSIGDFIVIAGTVPFISTIYVLDSWPYGLFLC
KLSEFLRDVSIGVTVLTLTILSIDRYVAIAMPLLNHKGRRHTRRTVTIFLAISVWIVAILMAIP
GTHYSFVMQVQATPNLHYSVCYPFPPEMWPWYPKLMVLLKFLIQYAIPLVIIGTFYCLMARQLI
RTSRAHLSQTSCGGVAHLKQMKARVKVAKIALAFVLLFAVCFFPNHVFMMWYYFAPNAPSQYNS
FWHVWKIMGYVMTFVNSCLNPVALYLVSGVFRNHFKHYLFCGRHPNVHARHNSYSFRTIHSSSM
SKCTASTKI

>CAL6258661.1 CCHaR-1 [Ixodes pacificus]
MSSLVVNATLATLRANRDPSASRAGINTSQTSHVNLTTGYGDSEEFVPYEQRLETYVVPTLFAL
IFIVGLLGNGTLILVFIRNRTMRSVPNIYIMSLSIGDFIVIAGTVPFISTIYVLDSWPYGLFLC
KLSEFLRDVSIGVTVLTLTILSIDRYVAIAMPLLNHKGRRHTRRTVTILLAISVWIVAILMAIP
GTHYSFVMQVQATPNLHYSVCYPFPPEMWPWYPKLMVLLKFLIQYAIPLVIIGTFYCLMARQLI
RTSRAHLSQTSCGGVAHLKQMKARVKVAKIALAFVLLFAVCFFPNHVFMMWYYFAPNAPSQYNS
FWHVWKIMGYVMTFVNSCLNPVALYLVSGVFRNHFKHYLFCGRHPNVHARHNSYSFRTIHSSSM
SKCTASTKI

>CAL6176764.1 CCHaR-1 [Ixodes hexagonus]
MSSLIANTTLAALRASRDPSTSRVSANTSHTSHVNLTTGYGDSEEFVPYEQRLETYVVPTLFAF
IFLVGLLGNGTLILVFLRNRTMRSVPNIYIMSLSIGDFIVIAGTVPFISTIYVLDSWPYGLFLC
KLSEFLRDVSIGVTVLTLTILSIDRYVAIAMPLLNHKGRRHTRRTITVLLTISVWIVAVLMAIP
GTHYSFVMQVQATPDLHYSVCYPFPPEMWPWYPKLMVLLKFLIQYAIPLVIIGTFYCLMARQLI
RTSRAHLSQTSCGGIAHLKQMKARVKVAKIALAFVVLFAVCFFPNHVFMMWYYFAPDAPSHYNS
FWHVWKIMGYVMTFVNSCLNPVALYLVSGVFRNHFKHYLFCGRHPNVRARQNSYSFRTIHSSSM
SKCTASTKI

>JXMZ01044463.1 CCHaR-1 shotgun[Ixodes ricinus]
MSSLVVNATLATLRANRDPSASRAGINTSQTSHVNLTTGYGDSEEFVPYEQRLETYVVPTLFAL
IFIVGLLGNGTLILVFIRNRTMRSVPNIYIMSLSIGDFIVIAGTVPFISTIYVLDSWPYGLFLC
KLSEFLRDVSIGVTVLTLTILSIDRYVAIAMPLLNHKGRRHTRRTVTIVLAISVWIVAILMAIP
GTHYSFVMQVQATPNLHYSVCYPFPPEMWPWYPKLMVLLKFLIQYAIPLVIIGTFYCLMARQLI
RTSRAHLSQTSCGGVAHLKQMKARVKVAKIALAFVLLFAVCFFPNHVFMMWYYFAPNAPSQYNS
FWHVWKIMGYVMTFVNSCLNPVALYLVSGVFRNHFKHYLFCGRHPNVHA

>XP_037578809.1 CCHamide-1 receptor-like [Dermacentor silvarum]
```

MADVPESYDMLSTSTLGPLVANVTSVTLRVSRDQSVSRSYSNTTPTSHLNLTTGHGDSEEFVPY
EQRLETYVVPTLFAFIFLVGLLGNGTLIVVFLRNRTMRSVPNIYIMSLSLGDFIVIAGTVPFIS
TIYILDSWPYGLFLCKLSEFLRDVSISVTVLTLTVLSIDRYVAIAMPLLNHKGRRHTRRTITIM
LTVAVWMIAILLAIPGAHFSFVMEVEATPELRYSVCYPFPPEMWPWYPKLMVLMKFLVQYAIPL
MIIGTFYCLMARQLIRTSRTHLTQPGCGGVAQLKQMKARVKVAKIALAFVVLFAVCFFPNHVFM
MWYYFAPDAPSHYNSFWHVWKIMGYVMTFVNSCLNPIALYLVSGVFRNHFKHYLFCSRRAASGV
HSRNNSYSFRTIHSSSMSKCTASTKI

>XP_037518906.1 CCHamide-1 receptor-like [Rhipicephalus sanguineus]
MADVPESYDMLSTSTLGPMVANVTSMTLRVTRDQSVSRIYSNSTPASHLNLTTGHGDSEEFVPY
EQRLETYVVPTLFAFIFLVGLLGNGTLILVFIRNRTMRSVPNIYIMSLSLGDFIVIAGTVPFIS
TIYILDSWPYGLFLCKLSEFLRDVSISVTVLTLTVLSIDRYVAIAMPLLNHKGRRHTRRTITIM
LTVAVWMVAVLLAIPGAHFSFVMEVEATPELKYSVCYPFPPEMWPWYPKLMVLMKFLVQYLLPL
AIIGTFYGLMARQLIRTSRANLTQPGCGGVAQLKQMKARVKVAKIALAFVILFAVCFFPNHVFM
MWYYFAPDAPSHYNSFWHVWKIMGYVMTFVNSCLNPIALYLVSGVFRNHFKHYLFCSRRAASGA
HSRNNSYSFRTIHSSSMSKCTASTKI

>XP_037280954.1 CCHamide-1 receptor-like [Rhipicephalus microplus]
MADVPESYDMLSSTLGPLVANATSMTLRVTRDQSVSRIYNYSMPTSHLNLTTGHGDSEEFVPYE
QRLETYVVPTLFAFIFLVGLLGNGTLILVFIRNRTMRSVPNIYIMSLSVGDFIVIAGTVPFIST
IYILDSWPYGLFLCKLSEFLRDVSISVTVLTLTVLSIDRYVAIAMPLLNHKGRRHTRRTITIML
TVAVWMVAVLLAIPGAHFSFVMEVEATPELRYSVCYPFPPEMWPWYPKLMVLMKFLVQYMLPLA
IIGTFYGLMARQLIRTSRANLTQPGCGGVAQLKQMKARVKVAKIALAFVILFAVCFFPNHVFMM
WYYFAPDAPSHYNSFWHVWKIMGYVMTFVNSCLNPIALYLVSGVFRNHFKHYLFCSRRAASGIH
SRNNSYSFRTIHSSSMSKCTASTKI

>XP_022651431.1 CCHamide-1 receptor-like [Varroa destructor]
MRMDMDELWNLTSSELLGDLPFLGQLLSATGNGSDTILPADNISFTLNDEEELSYIPLGKHLET
YLAPPVFTLIFGVGLIGNGTLMLIFLRNKAMRSVPNIYIMSLSLGDLFVITGTVPFICAIYVLD
SWPFGLFLCKLSEFMRDLSIGVTVLTLTMLSIDRYIAIALPLYKRKGGRHDRRVTIVITCLVWF
IASLMAIPGAYFSYLGEAHLPGRPPIVYCHPFPQHMQPWYPRLMVMMKFLVQYVIPLIVIATFY
ILMSRSLIKTAKSTLCDQKNAAAKKQQKARVKVAKISLCFVLIFAVCFFPNHVVMIWLYFHPNA
HQNYNDFWHLFKLFGFVLTFVNSCLNPIALYFVSGVFRSYFKAYICCQPRTRTVVNNSMLSYRQ
SLHPSVQIDSTSLRASLISRHSGHGVYDHRKNSTRI

>XP_028968249.1 CCHamide-1 receptor-like [Galendromus occidentalis]
MAIETLNLIRLMDWRKIRDLCPNKFEAQGMAPLALMEDLSNMTSAEFLDELPSLNQLLANDSTN
ISFILTDGETYFSVSEEASFVPLGKRLETYLAPPVFTIIFLVGLIGNGTLMLIFXKNKAMRSVP
NIYIMSLSVGDLFVITGTVPFICAIYVLDSWPFGLFLCKLSEFMRDVSIGVTVLTLMMLSVDRY
IAIALPLYKRKGGRHDRRVTLFITGLVWLIASLMALPGAYFSYLGEAHIPGRKTIVYCHPFPQH
MHPWYPRLIVMLKFTMMYVIPLVIIATFYILMARSLIKTAKSALCDQKNAAAKKQQKARVKVAK
ISLCFVLIFAVCFFPNHVVMIWLYYHPTAHQDYNDFWHLFKLFGYVLTFVNSCLNPIALFFISG
AFRSYFKMYLFCQPRSKNIIQNSMLSYRHSLHPSVRADSTSPRASLISRHSVNDRRNSTRI

>XP_022693689.1 CCHamide-1 receptor-like isoform X2 [Varroa jacobsoni]
MREELSYIPLGKHLETYLAPPVFTLIFGVGLIGNGTLMLIFLRNKAMRSVPNIYIMSLSLGDLF
VITGTVPFICAIYVLDSWPFGLFLCKLSEFMRDLSIGVTVLTLTMLSIDRYIAIALPLYKRKGG
RHDRRVTIVITCLVWFIASLMAIPGAYFSYLGEAHLPGRPPIVYCHPFPQHMQPWYPRLMVMMK
FLVQYVIPLIVIATFYILMSRSLIKTAKSTLCDQKNAAAKKQQKARVKVAKISLCFVLIFAVCF
FPNHVVMIWLYFHPNAHQNYNDFWHLFKLFGFVLTFVNSCLNPIALYFVSGVFRSYFKAYICCQ
PRTRTVVNNSMLSYRQSLHPSVQIDSTSLRASLISRHSGHGVYDHRKNSTRI

>NP_611241.2 CCHamide-1 receptor [Drosophila melanogaster]
MIANLVSMETDLAMNIGLDTSGEAPTALPPMPNVTETLWDLAMVVSQSTQWPLLDTGSSENFSE
LVTTETPYVPYGRRPETYIVPILFALIFVVGVLGNGTLIVVFLSVRQMRNVPNTYILSLALADL
LVIITTVPLASTVYTVEYWPYGSFLCSLSEFMKDVSIGVSVFTLTALSGDRYFAIVDPLRKFHA
HGGGRRATRMTLATAVSIWLLAILCGLPALIGSNLKHLGINEKSIVICYPYPEEWGINYAKSMV
LLHFLVYYAIPLVVIAVFYVLIALHLMYSASVPGEIQGAVRQVRARRKVAVTVLAFVVIFGICF
LPYHVFFLWFYFWPTAQDDYNAFWHVLRIVAYCMSFANSCANPVALYFVSGAFRKHFNRYLFCR
GASGRRKKRGQHDTFCMHRDTSLTSTASKRFQSRHSCYQSTIRSCRLQETTITTLPNGGNQNGA
NISAVELALPVLQAPGHNEAHAPPSYGFLPLNEIVQQTRSSPAKFQESLLN

>NP_610199.2 CCHamide-2 receptor, isoform A [Drosophila melanogaster]
MYASLMDVGQTLAARLADSDGNGANDSGLLATGQGLEQEQEGLALDMGHNASADGGIVPYVPVL
DRPETYIVTVLYTLIFIVGVLGNGTLVIIFFRHRSMRNIPNTYILSLALADLLVILVCVPVATI
VYTQESWPFERNMCRISEFFKDISIGVSVFTLTALSGERYCAIVNPLRKLQTKPLTVFTAVMIW
ILAILLGMPSVLFSDIKSYPVFTATGNMTIEVCSPFRDPEYAKFMVAGKALVYYLLPLSIIGAL
YIMMAKRLHMSARNMPGEQQSMQSRTQARARLHVARMVVAFVVVFFICFFPYHVFELWYHFYPT
AEEDFDEFWNVLRIVGFCTSFLNSCVNPVALYCVSGVFRQHFNRYLCCICVKRQPHLRQHSTAT
GMMDNTSVMSMRRSTYVGGTAGNLRASLHRNSNHGVGGAGGGVGGGVGSGRVGSFHRQDSMPLQ
HGNAHGGGAGGGSSGLGAGGRTAAVSEKSFINRYESGVMRY

>AAH95542.1 Neuromedin B receptor [Homo sapiens]
MPSKSLSNLSVTTGANESGSVPEGWERDFLPASDGTTTELVIRCVIPSLYLLIITVGLLGNIML
VKIFITNSAMRSVPNIFISNLAAGDLLLLLTCVPVDASRYFFDEWMFGKVGCKLIPVIQLTSVG
VSVFTLTALSADRYRAIVNPMDMQTSGALLRTCVKAMGIWVVSVLLAVPEAVFSEVARISSLDN
SSFTACIPYPQTDELHPKIHSVLIFLVYFLIPLAIISIYYCHIAKTLIKSAHNLPGEYNEHTKK
QMETRKRLAKIVLVFVGCFIFCWFPNHILYMYRSFNYNEIDPSLGHMIVTLVARVLSFGNSCVN
PFALYLLSESFRRHFNSQLCCGRKSYQERGTSYLLSSSAVRMTSLKSNAKNMVTNSVLLNGHSM
KQEMAM

>AAB19411.1 ETB endothelin receptor [Homo sapiens]
MQPPPSLCGPALVALVLACGLSRIWGEERGFPPDRATPLLQTAEIMTPPTKTLWPKGSNASLAR
SLAPAEVPKGDRTAGSPPRTISPPPCQGPIEIKETFKYINTVVSCLVFVLGIIGNSTLLRIIYK
NKCMRNGPNILIASLALGDLLHIVIDIPINVYKLLAEDWPFGAEMCKLVPFIQKASVGITVLSL
CALSIDRYRAVASWSRIKGIGVPKWTAVEIVLIWVVSVVLAVPEAIGFDIITMDYKGSYLRICL
LHPVQKTAFMQFYKTAKDWWLFSFYFCLPLAITAFFYTLMTCEMLRKKSGMQIALNDHLKQRRE
VAKTVFCLVLVFALCWLPLHLSRILKLTLYNQNDPNRCELLSFLLVLDYIGINMASLNSCINPI
ALYLVSKRFKNCFKSCLCCWCQSFEEKQSLEEKQSCLKFKANDHGYDNFRSSNKYSSS

```
>NP_001718.1 bombesin receptor subtype-3 [Homo sapiens]
MAQRQPHSPNQTLISITNDTESSSSVVSNDNTNKGWSGDNSPGIEALCAIYITYAVIISVGILG
NAILIKVFFKTKSMQTVPNIFITSLAFGDLLLLLTCVPVDATHYLAEGWLFGRIGCKVLSFIRL
TSVGVSVFTLTILSADRYKAVVKPLERQPSNAILKTCVKAGCVWIVSMIFALPEAIFSNVYTFR
DPNKNMTFESCTSYPVSKKLLQEIHSLLCFLVFYIIPLSIISVYYSLIARTLYKSTLNIPTEEQ
SHARKQIESRKRIARTVLVLVALFALCWLPNHLLYLYHSFTSQTYVDPSAMHFIFTIFSRVLAF
SNSCVNPFALYWLSKSFQKHFKAQLFCCKAERPEPPVADTSLTTLAVMGTVPGTGSIQMSEISV
TSFTGCSVQAEDRF

>NP_005305.1 gastrin-releasing peptide receptor [Homo sapiens]
MALNDCFLLNLEVDHFMHCNISSHSADLPVNDDWSHPGILYVIPAVYGVIILIGLIGNITLIKI
FCTVKSMRNVPNLFISSLALGDLLLLITCAPVDASRYLADRWLFGRIGCKLIPFIQLTSVGVSV
FTLTALSADRYKAIVRPMDIQASHALMKICLKAAFIWIISMLLAIPEAVFSDLHPFHEESTNQT
FISCAPYPHSNELHPKIHSMASFLVFYVIPLSIISVYYYFIAKNLIQSAYNLPVEGNIHVKKQI
ESRKRLAKTVLVFVGLFAFCWLPNHVIYLYRSYHYSEVDTSMLHFVTSICARLLAFTNSCVNPF
ALYLLSKSFRKQFNTQLLCCQPGLIIRSHSTGRSTTCMTSLKSTNPSVATFSLINGNICHERYV
```

#### Percent Identity Matrix for CCHaR and vertebrate homologs

| Species | Accesion Number | XP_042147864 | CAL6223886 | CAL6258661 | CAL6176764 | JXMZ01044463 | XP_037578809 | XP_037518906 | XP_037280954 | XP_022651431 | XP_028968249 | XP_022693689 | NP_611241 | NP_610199 | AAH95542 | AAB19411 | NP_001718 | NP_005305 |
|---|---|---|---|---|---|---|---|---|---|---|---|---|---|---|---|---|---|---|
| *I. scapularis* | XP_042147864 | 100 | 98.7 | 99.5 | 93.4 | 99.2 | 85.2 | 84.2 | 84.5 | 55.2 | 54.7 | 59.4 | 46.6 | 39.4 | 35.8 | 29.4 | 33.2 | 33.5 |
| *I. persulcatus* | CAL6223886 | 98.7 | 100 | 99.2 | 93.9 | 99.2 | 85.5 | 84.2 | 84.7 | 55.2 | 55.0 | 59.4 | 46.6 | 39.7 | 35.8 | 29.4 | 33.5 | 33.2 |
| *I. pacificus* | CAL6258661 | 99.5 | 99.2 | 100 | 93.9 | 99.7 | 85.2 | 84.0 | 84.5 | 55.2 | 54.7 | 59.4 | 46.6 | 39.4 | 35.8 | 29.4 | 33.5 | 33.5 |
| *I. hexagonus* | CAL6176764 | 93.4 | 93.9 | 93.9 | 100 | 93.5 | 86.3 | 84.5 | 84.7 | 55.0 | 54.7 | 58.6 | 46.1 | 38.9 | 34.9 | 29.1 | 33.0 | 33.5 |
| *I. ricinus* | JXMZ01044463 | 99.2 | 99.2 | 99.7 | 93.5 | 100 | 84.6 | 83.2 | 83.7 | 56.8 | 56.0 | 61.5 | 48.3 | 40.7 | 36.9 | 30.8 | 35.2 | 33.7 |
| *D. silvarum* | XP_037578809 | 85.2 | 85.5 | 85.2 | 86.3 | 84.6 | 100 | 95.4 | 94.9 | 55.1 | 53.4 | 59.9 | 45.3 | 38.3 | 33.4 | 28.9 | 31.3 | 33.1 |
| *R. sanguineus* | XP_037518906 | 84.2 | 84.2 | 84.0 | 84.5 | 83.2 | 95.4 | 100 | 97.6 | 54.1 | 52.9 | 59.1 | 43.7 | 38.1 | 34.3 | 30.0 | 31.3 | 32.5 |
| *R. microplus* | XP_037280954 | 84.5 | 84.7 | 84.5 | 84.7 | 83.7 | 94.9 | 97.6 | 100 | 54.0 | 53.5 | 58.8 | 43.4 | 37.8 | 33.8 | 29.8 | 31.4 | 32.3 |
| *V. destructor* | XP_022651431 | 55.2 | 55.2 | 55.2 | 55.0 | 56.8 | 55.1 | 54.1 | 54.0 | 100 | 79.2 | 99.5 | 40.8 | 39.3 | 34.7 | 30.1 | 28.9 | 34.6 |
| *G. occidentalis* | XP_028968249 | 54.7 | 55.0 | 54.7 | 54.7 | 56.0 | 53.4 | 52.9 | 53.5 | 79.2 | 100 | 84.8 | 43.0 | 36.6 | 34.1 | 28.9 | 30.5 | 34.5 |
| *V. jacobsoni* | XP_022693689 | 59.4 | 59.4 | 59.4 | 58.6 | 61.5 | 59.9 | 59.1 | 58.8 | 99.5 | 84.8 | 100 | 43.3 | 40.7 | 34.9 | 31.2 | 29.9 | 35.3 |
| *D. melanogaster CCHaR-1* | NP_611241 | 46.6 | 46.6 | 46.6 | 46.1 | 48.3 | 45.3 | 43.7 | 43.4 | 40.8 | 43.0 | 43.3 | 100 | 46.5 | 34.0 | 28.7 | 33.4 | 35.1 |
| *D. melanogaster CCHaR-2* | NP_610199 | 39.4 | 39.7 | 39.4 | 38.9 | 40.7 | 38.3 | 38.1 | 37.8 | 39.3 | 36.6 | 40.7 | 46.5 | 100 | 36.0 | 28.1 | 32.1 | 34.6 |
| *H. sapiens BB1* | AAH95542 | 35.8 | 35.8 | 35.8 | 34.9 | 36.9 | 33.4 | 34.3 | 33.8 | 34.7 | 34.1 | 34.9 | 34.0 | 36.0 | 100 | 30.0 | 46.7 | 56.7 |
| *H. sapiens ETBR* | AAB19411 | 29.4 | 29.4 | 29.4 | 29.1 | 30.8 | 28.9 | 30.0 | 29.8 | 30.1 | 28.9 | 31.2 | 28.7 | 28.1 | 30.0 | 100 | 28.3 | 29.6 |
| *H. sapiens BRS3* | NP_001718 | 33.2 | 33.5 | 33.5 | 33.0 | 35.2 | 31.3 | 31.3 | 31.4 | 28.9 | 30.5 | 29.9 | 33.4 | 32.1 | 46.7 | 28.3 | 100 | 50.8 |
| *H. sapiens GRPR* | NP_005305 | 33.5 | 33.2 | 33.5 | 33.5 | 33.7 | 33.1 | 32.5 | 32.3 | 34.6 | 34.5 | 35.3 | 35.1 | 34.6 | 56.7 | 29.6 | 50.8 | 100 |

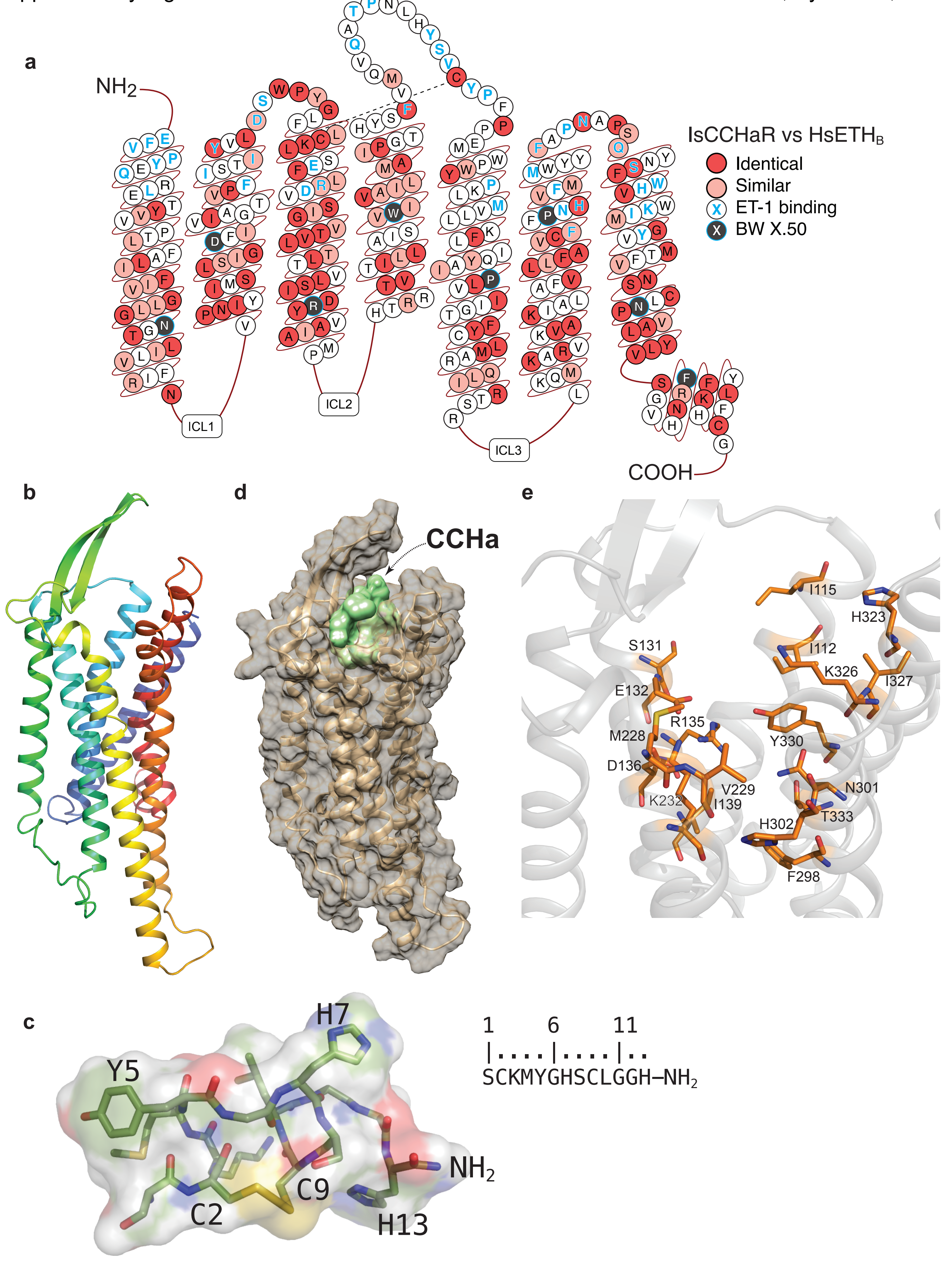
a
NH2
COOH
ICL1
ICL2
ICL3
IsCCHaR vs HsETHB
Identical
Similar
ET-1 binding
BW X.50
b
d
CCHa
e
I115
H323
S131
I112
K326
I327
E132
R135
M228
Y330
D136
V229
N301
K232
I139
H302
T333
F298
c
H7
Y5
NH2
C9
C2
H13
1 6 11
SCKMYGHSCLGGH–NH2

#### Amino sequences *Ixodes* GNA subunits used for alignments

>XP_029846564 GNAQ I. scapularis
MMACCISEEAKEQRRINQEIERQLRKDKRDARRELKLLLLGTGESGKSTFIKQMRIIHGHGYNEEDKKGFIKL
VYQNIFMAMQSMIKAMDMLKIQYTNPDNIEHANLVGTVDYETVTTFDGPYVVAIKKLWADGGIQECYDRRREY
QLTDSAKYYLSDVDRIAATDYLPTQQDILRVRVPTTGIIEYPFDLDSIIFRMVDVGGQRSERRKWIHCFENVT
SIIFLVALSEYDQILFESDNENRMEESKALFKTIITYPWFQNSSVILFLNKKDLLEEKIMYSHLVDYFPEYDG
PKKDAIHAREFILKMFVDLNPDSEKIIYSHFTCATDTENIRFVFAAVKDTILQLNLKEYNLV

>CAL6172950 I. hexagonus
MMACCISEEAKEQRRINQEIERQLRKDKRDARRELKLLLLGTGESGKSTFIKQMRIIHGHGYNEEDKKGFIKL
VYQNIFMAMQSMIKAMDMLKIQYTNPDNIEHANLVGTVDYETVTTFDGPYVVAIKKLWADGGIQECYDRRREY
QLTDSAKYYLSDVDRIAATDYLPTQQDILRVRVPTTGIIEYPFDLDSIIFRMVDVGGQRSERRKWIHCFENVT
SIIFLVALSEYDQILFESDNENRMEESKALFKTIITYPWFQNSSVILFLNKKDLLEEKIMYSHLVDYFPEYDG
PKQDHESARNFILKVYLDSNPDKKRMVYSHFTCATDTENIRFVFAAVKDTILQLNLKEYNLV

>XP_029846565 GNAQ I. scapularis
MMACCISEEAKEQRRINQEIERQLRKDKRDARRELKLLLLGTGESGKSTFIKQMRIIHGHGYNEEDKKGFIKL
VYQNIFMAMQSMIKAMDMLKIQYTNPDNIEHANLVGTVDYETVTTFDGPYVVAIKKLWADGGIQECYDRRREY
QLTDSAKYYLSDVDRIAATDYLPTQQDILRVRVPTTGIIEYPFDLDSIIFRMVDVGGQRSERRKWIHCFENVT
SIIFLVALSEYDQILFESDNENRMEESKALFKTIITYPWFQNSSVILFLNKKDLLEEKIMYSHLVDYFPEYDG
PKQDHITAREFLLRQFQSQKPGSDFMYSHFTCATDTENIRFVFAAVKDTILQLNLKEYNLV

>KAG0422235 I. persulcatus
MMACCISEEAKEQRRINQEIERQLRKDKRDARRELKLLLLGTGESGKSTFIKQMRIIHGHGYNEEDKKGFIKL
VYQNIFMAMQSMIKAMDMLKIQYTNPDNIEHANLVGTVDYETVTTFDGPYVVAIKKLWADGGIQECYDRRREY
QLTDSAKYYLSDVDRIAATDYLPTQQDILRVRVPTTGIIEYPFDLDSIIFRMVDVGGQRSERRKWIHCFENVT
SIIFLVALSEYDQILFESDNENRMEESKALFKTIITYPWFQNSSVILFLNKKDLLEEKIMYSHLVDYFPEYDG
PKKDAIHAREFILKMFVDLNPDSEKIIYSHFTCAT

>CAL6216232 I. pacificus
MMACCISEEAKEQRRINQEIERQLRKDKRDARRELKLLLLGTGESGKSTFIKQMRIIHGHGYNEEDKKGFIKL
VYQNIFMAMQSMIKAMDMLKIQYTNPDNIEHANLVGTVDYETVTTFDGPYVVAIKKLWADGGIQECYDRRREY
QLTDSAKYGLCGTLRALSRDLGPNESASFKMAVTSTGAARATWCFRSSVLREIAMVDVGGQRSERRKWIHCFE
NVTSIIFLVALSEYDQILFESDNENRMEESKALFKTIITYPWFQNSSVILFLNKKDLLEEKIMYSHLVDYFPE
YDGPKQDHESARNFILKVYLDSNPDTKRMVYSHFTCATDTENIRFVFAAVKDTILQLNLKEYNLV

>XP_002406461 GNAO I. scapularis
MGCAMSAEERAALARSKQIEKNLKEDGIQAAKDIKLLLLGAGESGKSTIVKQMKIIHDSGFTQEDFKQYKPVV
YSNTIQSMVAILRAMPNLGISFGNNEREADAKMVFDVVARMEDTEPFSEELLSAMKRLWTDSGVQECFGRSNE
YQLNDSAKYFLDDLDRLGKKEYMPTEQDILRTRVKTTGIVEVHFSFKNLNFKLFDVGGQRSERKKWIHCFEDV
TAIIFCVAMSEYDQVLHEDETTNRMQESLKLFDSICNNKWFTDTSIILFLNKKDLFEEKIKKSPLTICFPEYT
GAQEYGEAAAYIQAQFEAKNKSTTKEIYCHMTCATDTTNIQFVFDAVTDVIIANNLRGCGLY

>XP_029828399 GNAI I. scapularis
MGCAVSTAADKEAAERSKKIDRDLRADGERQAREVKLLLLGAGESGKSTIVKQMKIIHESGYTSDECKLYRPV
VHSNTIQSLLAIIRAMGQLKIDFKDPSRADDARQFFTVAGATSSQECEITPELASMMKRLWQDHGVQHCFSRS
REYQLNDSASYYLNALDRISQASYTPTQQDVLRTRVKTTGIVETHFVFKELHFKMFDVGGQRSERKKWIHCFE
GVTAIIFCVALSGYDLVLAEDEEMNRMIESMKLFDSICNNKWFVETSIILFLNKKDLFEEKIVRSPLTICFPE
YPGSNTYEEAAAYIQMKFESLNKRRDTKEIYTHFTCATDTNNIQFVFDAVTDVIIKNNLKDCGLF

>CAL6190929 I. hexagonus

MGACLSLDEEERRARLRSSQIDRQIAELARQERNVIKILLLGAGESGKSTIVKQMKIIHSEGFSDEELRSFRP
TVLDNLLGSMKYVLTGMGILRINLENPKNKMYAQTVLSCQCCFDENIAMLPFVGSALKNLWNDKGIRLAVARG
FEYELNDSALYLFENMDRICSEKYMPSPRDVLRARVRTNGIIETHFKIDDIVFRMFDVGGQRSERRKWIQCFD
DVKALLYVVALSGYDMTLQEDPNVNRLQESLKLFASICNNMFFTDTSLVLFLNKLDLFRDKILYSERHLRYYL
PDYKGPDYDVDSGALFIQHRFQSRNRDPSKLVYPHFTTATDTSNVQVVFQVAMDTVLRGNLKTATLL

>CAL6192420 I. hexagonus
MAPPVSCCLSEEAKLQKRVNEQIERQLSRERFEAGRESRFLLLGPEEAGKTTFLKQIRLIQEGDARADRQDHV
RSVYQALLGAMQRIVRAMDALGIAYSDPSNAENAMLLAAVRCESLEPPLTEPVSEALRRLWEDDGVRQCYARR
AEYGGLPPFLPDVQRVTRPGYVPSREDYLRVRTPSRGVLEYTFQLDEFPVRLVDVGEQRSEPKKWIHCFEDVS
GVIFVAPLSDYDDPDKMERARALFSTIVSSPWFAESHLLLLLNKKDLLEQKIARRPLGAYFPAYRGPPGDPKA
AREFLRRSFADLKPERPVYFFFTCAID

>CAL6189997 I. hexagonus
MAGLLANCACFYRLKYGPDELESLQRSRKIDKMIQKDKQVIRRQVKVLLLGAGESGKSTFLKQMRIIHGYNFD
EEVLSEFRTVVFQNVVKGMKVLVDARDKLCIPWGDAANSGHGQRILRYEPSLFLDWETFERFAPSVRILWQDS
AIRTAFDRRVEFQLGDGVRYFFDNLDRIAAKDYLPTNQDILHARKATKGITEFVIPVNGIPFRFVDVGGQRSQ
RQKWFRCFDSVTSILFLVSSSEFDQVLLEDRCTNRVTESRSIFDTIVNNRCFAEVSIILFFNKTDLLQEKLQS
RTTSIADYFEDFRGDPHNLQHVQRFLVDWFYEVRRNRHKPLFHHFTTAVDTENIKVVFNAVRDTILQKNITQL
MLQ

>XP_002399234 GNAO I. scapularis
MGACLSQDEEERRARLRSSQIDRQIAELARQERNVIKILLLGAGESGKSTIVKQMKIIHNEGFSDEELRSFRP
TVLDNLLGSMKFVLTGMGILRINLENPKNKAYAQTVLGCQCCFDEGTAMLPFVGSALKNLWNDKGIRLAAARG
FEYELNDSALYLFENMDRICSEKYVPSPRDVLRARVRTNGIIETHFKIDDIVFRMFDVGGQRSERRKWIQCFD
DVKALLYVVALSGYDMTLQEDPSVNRLQESLKLFASICNNMFFTDTSLVLFLNKLDLFRDKILYSERHLRYYL
PDYKGPDYDVDSGALFIQHRFQSRNRDPGKLVYPHFTTATDTSNVQVVFQVAMDTVLRGNLKTATLL

>XP_029849143 GNA12 I. scapularis
MAGLLANCACLACFYRLKYGPDELESLQRSRKIDKMIQKDKQVIRRQVKLLLLGAGESGKSTFLKQMRIIHGF
SFDEEVLCEFRTVVFQNVVKGMKVLVDARDKLCIPWGDAANAHHGQRILRYEPNLALDWDTFSRFAPSVRILW
QDDAIRTAFDRRVEFQLGDGVRYFFDNLDRIAAKDYVPTNQDILHARKATKGITEFVIPVNGIPFRFVDVGGQ
RSQRQKWFRCFDSVTSILFLVSSSEFDQVLLEDRCTNRVTESRSIFDTIVNNRCFAEVSIILFFNKTDLLQEK
LQARTTSIADYFEDFRGDPHDLQQVQRFLVDWFYSVRRNRHKPLFHHFTTAVDTENIKVVFNAVRDTILQKNI
TQLMLQ

>XP_002435048 GNAQ I. scapularis
MAPPVPCCLSEEAKLQKRVNERIERQLSRERFEAGRESKFLLLGPEEAGKTTFLKQIRLIQEGDARADRADHV
RSVFQALLGAMQRIVRAMDALGIAYSDPSNAENAMLLAAVRCESLEPPLTEPVSEALRRLWEDDGVRECYARR
SEYGGLPPFLPDVQRVTRPGYVPSREDYLRVRTPSRGVLEYAFQLDEFPVRLVDVGEQRSEPKKWIHCFEDVS
GVIFVAPLSDYDDPAKMERTRDLFSTIVSSPWFAESHLLLLLNKKDLLEQKIERHPLGAYFPAFRGPPGDPKA
AREFLRRLFADLKPERPVYFFFTCAVDTDDVKYVLPTLREIACLRSCPSRGRQPSDASTASNAAIMVDEHYEE
M

>CAL6245647 I. persulcatus
MAPPVPCCLSEEAKLQKRVNERIERQLSRERFEAGRESKFLLLGPEEAGKTTFLKQIRLIQEGDARADRADHV
RSVFQALLGAMQRIVRAMDALGIAYSDPSNAENAMLLAAVRCESLEPPLTEPVSEALRRLWEDDGVRECYARR
SEYGGLPPFLPDVQRVTRPGYVPSREDYLRVRTPSRGVLEYAFQLDEFPVRLVDVGEQRSEPKKWIHCFEDVS
GVIFVAPLSDYDDPAKMERTRDLFSTIVSSPWFAESHLLLLLNKKDLLEQKIERHPLGAYFPAFRGPPGDPKA
AREFLRRLFADLKPERPVYFFFTCAVDTDDVKYVLPTLREIACLRSCPSRGRQPSDASTASNAAAAAATAATA
AAAAFQLPHTA

>CAL6253071 I. pacificus

MAPPVPCCLSEEAKLQKRVNERIERQLSRERFEAGRESKFLLLGPEEAGKTTFLKQIRLIQEGDARADRADHV
RSVFQALLGAMQRIVRAMDALGIAYSDPSNAENAMLLAAVRCESLEPPLTEPVSEALRRLWEDDGVRECYARR
SEYGGLPPFLPDVQRVTRPGYVPSREDYLRVRTPSRGVLEYAFQLDEFPVRLVDVGEQRSEPKKWIHCFEDVS
GVIFVAPLSDYDDPAKMERTRDLFSTIVSSPWFAESHLLLLLNKKDLLEQKIERHPLGAYFPAFRGPPGDPKA
AREFLRRLFADLKPERPVYFFFTCAVDTDDVKYVLPTLREIACLRSCPSRGRQPSDASTASNAAAAAATAATA
ATAAFQLPHTA

>CAL6240404 I. pacificus
MGCAMSAEERAALARSKQIEKNLKEDGIQAAKDIKLLLLGAGESGKSTIVKQMKIIHDSGFTQEDFKQYKPVV
YSNTIQSMVAILRAMPNLGISFGNNEREADAKMVFDVVARMEDTEPFSEELLSAMKRLWTDSGVQECFGRSNE
YQLNDSAKYFQHNSDSFMCLVVGQTIPPPFFFLVPSQATVAVFFSFCMFALFDVGGQRSERKKWIHCFEDVTA
IIFCVAMSEYDQVLHEDETTNRMQESLKLFDSICNNKWFTDTSIILFLNKKDLFEEKIKKSPLTICFPEYTGA
QEYGEAAAYIQAQFEAKNKSTTKEIYCHMTCATDTTNIQFVFDAVTDVIIANNLRGCGLY

>KAG0431228 I. persulcatus
MRIVMNPSTRLGLMTAEYLTVSRDSQIIHDSGFTQEDFKQYKPVVYSNTIQSMVAILRAMPNLGISFGNNERE
ADAKMVFDVVARMEDTEPFSEELLSAMKRLWTDSGVQECFGRSNEYQLNDSAKYFLDDLDRLGKKEYMPTEQD
ILRTRVKTTGIVEVHFSFKNLNFKLFDVGGQRSERKKWIHCFEDVTAIIFCVAMSEYDQVLHEDETTNRMQES
LKLFDSICNNKWFTDTSIILFLNKKDLFEEKIKKSPLTICFPEYTGAQEYGEAAAYIQAQFEAKNKSTTKEIY
CHMTCATDTTNIQFVFDAVTDVIIANNLRGCGLY

>CAL6250599 I. persulcatus
MGACLSQDEEERRARLRSSQIDRQIAELARQERNVIKILLLGAGESGKSTIVKQMKIIHNEGFSDEELRSFRP
TVLDNLLGSMKFVLTGMGILRINLENPKNKAYAQTVLGCQCCFDEGMAMLPFVGSALKNLWNDKGIRLAAARG
FEYELNDSALYLFENMDRICSEKYVPSPRDVLRARVRTNGIIETHFKIDDIVFRMFDVGGQRSERRKWIQCFD
DVKALLYVVALSGYDMTLQEDPGVNRLQESLKLFASICNNMFFTDTSLVLFLNKLDLFRDKILYSERHLRYYL
PDYKGPDYDVDSGALFIQHRFQSRNRDPGKLVYPHFTTATDTSNVQVVFQVAMDTVLRGNLKTATLL

>CAL6181757 I. hexagonus
MGCAMSAEERAALARSKQIEKNLKEDGIQAAKDIKLLLLGAGESGKSTIVKQMKIIHDSGFTQEDFKQYKPVV
YSNTIQSMVAILRAMPNLGISFGNNEREADAKMVFDVVARMEDTEPFSEELLSAMKRLWTDSGVQECFGRSNE
YQLNDSAKYFLDDLDRLGKKEYMPTEQDILRTRVKTTGIVEVHFSFKNLNFKLFDVGGQRSERKKWIHCFEDV
TAIIFCVAMSEYDQVLHEDETTRADSRSCHRPLFQHLLNNKAVALIFNLIEMVREKLENSDKTLGQHSSAQEY
GEAAAYIQAQFEAKNKSTTKEIYCHMTCATDTTNIQFVFDAVTDVIIANNLRGCGLY

>XP_040062629 GNASS I. scapularis
MGCFGGSSSKSDAEEDKRRKEANKKIEKQIQKDKQIYRATHRLLLLGAGESGKSTIVKQMRILHVNGFSEEEK
KQKIEDIKKNIRDAILTITGAMSTLVPPVQLQKSENQWRVDYIQDVASSPDFDYPAEFYEHTEILWKDKGVQA
AFERSNEYQLIDCAKYFLDRVATIKQLDYTPNEQDILRCRVLTSGIFETKFQVDKVNFHMFDVGGQRDERRKW
IQCFNDVTAIIFVTACSSYNMVLREDPNQNRLRESLDLFKSIWNNRWLRTISVILFLNKQDLLAEKIKAGKSR
LEEYFPEFAHYQTPSDAVIETGEDPEVIRAKYFIRDEFLRISTASGDGKHYCYPHFTCAVDTENIRRVFNDCR
DIIQRMHLRQYELL

>CAL6181233 I. hexagonus
MGCFGGSSSKSDAEEDKRRKEANKKIEKQIQKDKQIYRATHRLLLLGAGESGKSTIVKQMRILHVNGFSEEEK
KQKIEDIKKNIRDAILTITGAMSTLVPPVQLQKSENQWRVDYIQDVASSPDFDYPAEFYEHTEILWKDKGVQA
AFERSNEYQLIDCAKYFLDRVATIKQPDYTPNEQDILRCRVLTSGIFETKFQVDKVNFHMFDVGGQRDERRKW
IQCFNDVTAIIFVTACSSYNMVLREDPNQNRLRESLDLFKSIWNNRWLRTISVILFLNKQDLLAEKIKAGKSR
LEEYFPEFAHYQTPSDEPRLNPFQPGTPHTVIETGEDPEVIRAKYFIRDEFLRISTASGDGKHYCYPHFTCAV
DTENIRRVFNDCRDIIQRMHLRQYELL

>CAL6177226 I. hexagonus
MGCAVSTAADKEAAERSKKIDRDLRADGERQAREVKLLLLGAGESGKSTIVKQMKIIHESGYTSDECKLYRPV
VHSNTIQSLLAIIRAMGQLKIDFKDPSRAVSRAQTAWNICPFVGLARKPPNLHLLMKAKVVVVAPQLGCRRSR

PTRCNALLPGCSYLNALDRISQASYTPTQQDVLRTRVKTTGIVETHFVFKELHFKMFDVGGQRSERKKWIHCF
EGVTAIIFCVALSGYDLVLAEDEEMNRMIESMKLFDSICNNKWFVETSIILFLNKKDLFEEKIVRSPLTICFP
EYPGSNTYEEAAAYIQMKFESLNKRRDTKEIYTHFTCATDTNNIQFVFDAVTDVIIKNNLKDCGLF

>KAG0428246 I. persulcatus
MKSSAGAGESGKSTIVKQMRILHVNGFSEEEKKQKIEDIKKNIRDAILTITGAMSTLVPPVQLQKSENQWRVD
YIQDVASSPDFDYPAEFYEHTEILWKDKGVQAAFERSNEYQLIDCAKYFLDRVATIKQLDYTPNEQDILRCRV
LTSGIFETKFQVDKVNFHMFDVGGQRDERRKWIQCFNDVTAIIFVTACSSYNMVLREDPNQNRLRESLDLFKS
IWNNRWLRTISVILFLNKQDLLAEKIKAGKSRLEEYFPEFAHYQTPSDAVIETGEDPEVIRAKYFIRDEFLRI
STASGDGKHYCYPHFTCAVDTENIRRVFNDCRDIIQRMHLRQYELL

>CAL6219429 I. pacificus
GQRVLQDDARQFFTVAGATSSQECEITPELASMMKRLWQDHGVQHCFSRSREYQLNDSASYYLNALDRISQAS
YTPTQQDVLRTRVKTTGIVETHFVFKELHFKMFDVGGQRSERKKWIHCFEGVTAIIFCVALSGYDLVLAEDEE
MNRMIESMKLFDSICNNKWFVETSIILFLNKKDLFEEKIVRSPLTICFPEYPGDNTYEEAAAYIQMKFESLNK
RRDTKEIYTHFTCATDTNNIQFVFDAVTDVIIKNNLKDCGLF

>CAL6248271 I. pacificus
MGACLSQDEEERRARLRSSQIDRQIAELARQERNVIKILLLGAGESGKSTIVKQMKIIHNEGFSDEELRSFRP
TVLDNLLGSMKFVLTGMGILRINLENPKNKAYAQTVLGCQCCFDEGTAMLPFVGSALKNLWNDKGIRLAAARG
FEYELNDSALYLFENMDRICSEKYVPSPRDVLRARVRTNGIIETHFKIDDIVFRMFDVGGQRSERRKWIQCFD
DVKALLYVVALSGYDMTLQEDPGVNRLQESLKLFASICNNMFFTDTSLVSFVLGKLF

>CAL6245999 I. pacificus
MAGLLANCACLACFYRLKYGPDELESLQRSRKIDKMIQKDKQVIRRQVKLLLLGAGESGKSTFLKQMRIIHGF
SFDEEVLCEFRTVVFQNVVKGMKVLVDARDKLCIPWGDAANAHHGERILRYEPSQALDWDTFSRFAPSVRILW
QDDAIRTAFDRRVEFQLGDGVRYFFDNLDRIAAKDYVPTNQDILHARKATKGITEFVIPVNGIPFRFVDVGGQ
RSQRQKWFRCFDSVTSILFLVSSSEFDQV

>XP_042144474 GNAS I. scapularis
MGCFKTKSPEQKVNDKLNGVIQQWKKREGGLIKILLLGAGESGKTTILKQMTILHRNGFTSEERTEKAHEIRW
NLLEAMKELTMHMATLNPPVELEDSANEESFKFVQSLYLLADYEFPQCFYFHIKRLWSDAGVQECYRRSNEFF
LIESSKYFLDQIDVISDESYIPTEQDILRCRRRTTNVKKVEFEATIPKKYGHGIQEFWMFDVGGQRGERRKWF
SVFAGIDAVLFLAATNGFDAVLREDSTVNRLQEALDLFRDVWNSKYLRGSGMILFLNKQDLLKEKVERGSKIE
DHFPAYKCFKTHKAEENSSEYTRVRTFIRELFLNITRAEVPRSCSDCAHSGLLSSESMVRECFWHYTTATDTD
NVQNVFNDVHSMIILSNLSKMGPS

>CAL6216198 I. pacificus
MGCFKTKSPEQKVNDKLNGVIQQWKKREGGLIKILLLGAGESGKTTILKQMTILHRNGFTLEERIEKAHEIRW
NLLEAMKELTMHMETLNPPVELEDSANMESFKFVQGLYLLADYEFPQCFYFHIKRLWSDAGVQECYRRSNEFF
LIESSKYFLDQIDAISDESYIPTDQDILRCRRRTTNVKKVEFEATIPKKYGHGIQEFWMFDVGGQRGERRKWF
SVFAGIDAVLFLAATNGFDAVLREDSTVNRLQEALDLFRDVWNSKYLRGSGMILFLNKQDLLKEKVERGSKIE
DHFPAYKCFKTHKAEDNSSEYTRVRTFIRELFLNITRAEVPRSCIDCAPSGLLSSESTVRECFWHYTTATDTD
NVQNVFNDVHSMIILSNLSKMGPS

>KAG0443856 I. persulcatus
MCLFCGFQGDGVRYFFDNLDRIAAKDYVPTNQDILHARKATKGITEFVIPVNGIPFRFVDVGGQRSQRQKWFR
CFDSVTSILFLVSSSEFDQVLLEDRCTNRVTESRSIFDTIVNNRCFAEVSIILFFNKTDLLQEKLQARTTSIA
DYFEDFRGDPHDLQQVQRFLVDWFYSVRRNRHKPLFHHFTTAVDTENIKVVFNAVRDTILQKN

>CAL6215671 I. persulcatus
MCVYYFIDVLIFRILYQELHFSFSRRFSLNAQFTGAGESGKTTILKQMTILHRNGFTSEERTEKAHEIRWNLL
EAMKELTMHMATLNPPVELADSANVESFKFVQSLYLLADYEFPQCFYFHIKRLWSDAGVQECYRRSNEFFLIE
SSKYFLDQIDAISDESYIPTDQDILRCRRRTTNVKKVEFEATIPKKYGHGIQEFWMFDVGGQRGERRKWFSVF

AGIDAVLFLAATNGFDAVLREDSTVNRLQEALDLFRDVWNSKYLRGSGMILFLNKQDLLKEKVERGFKIEDHF
PAYKCFKTHKAEDNSSEYTRVRSFIRELFLNITRAEVPRSCIDCAHSGLLSSESTVRECFWHYTTATDTDNVQ
NVFNDVHSMIILSNLSKMGPS

>CAL6256702_KAG0424332 I. persulcatus
RIIHESGYTSDECKLYRPVVHSNTIQSLLAIIRAMGQLKIDFKDPSRADDARQFFTVAGATSSQECEITPELA
SMMKRLWQDHGVQHCFSRSREYQLNDSASYYLNALDRISQASYTPTQQDVLRTRVKTTGIVETHFVFKELHFK
MFDVGGQRSERKNRMIESMKLFDSICNNKWFVETSIILFLNKKDLFEEKIVRSPLTICFPEYPDEDWSCRTSS
LVRFDLSLVNRKEEIYTHFTCATDTNNIQFVFDAVTDVIIKNNLKDCGLF

>CAL6199964 I. hexagonus
MGCFKTKSPEEKVNDKLNGVIQQWKKREGGLIKILLLGAGESGKTTILKQMTILHTDGFTSEERREKAHEIRW
NLLEPMKELTVHMAKLNPPIELEDPANVESFKFVKSLYPLADYDFPQCFYQHIKRLWSDAGIQECYRRSNEFF
LIESSKYFLDQVDVISDENYVPTDQDILRCRKRTSNVKKVEFEAIIPKKYGHGIQEFWGGQDIGGAGGQGGRR
WGRGGRGCCRVFVLVGSDNYWKIVTGDIHRVNEGLTGVNIIFGGMPQGSYLRGSGMILFLNKQDLLKEKVERG
SKIEDHFPEYKCFKTHKAEENSSEYTRVRAFIRELFLNITRTEVPRTSLDRAHSGLLGSESPVRECFWHYTTA
TDTDNVQNVFNDVHSMIILSNLAKMGPS

#### DNA and amino sequences used to express *Ixodes scapularis* GNAQ

>IsGNAQ1 Humanized Ixodes scapularis G protein alpha q subunit coding region based upon XP_029846564.1
ATGGCTTGTTGTATCTCCGAGGAGGCTAAAGAGCAACGGAGAATAAATCAGGAAATCGAACGCC
AGCTTAGAAAGGACAAGCGCGACGCACGCCGGGAACTCAAGCTGTTGCTGCTCGGAACTGGTGA
ATCCGGAAAGAGTACATTCATTAAGCAAATGCGCATTATACACGGTCACGGCTACAATGAGGAA
GATAAAAAGGGCTTCATTAAACTGGTTTATCAGAATATATTTATGGCTATGCAATCCATGATTA
AAGCGATGGATATGCTTAAAATTCAGTACACCAATCCAGACAACATAGAACACGCGAATCTTGT
CGGGACTGTAGACTATGAGACCGTAACAACATTTGATGGCCCCTACGTCGTAGCGATCAAAAAG
CTTTGGGCTGACGGAGGCATCCAGGAATGCTATGATCGGAGACGGGAATACCAGCTCACAGATA
GTGCGAAATATTACCTCAGTGACGTTGATCGCATCGCCGCGACTGATTATCTCCCCACGCAGCA
AGATATTCTTCGGGTGCGCGTCCCAACGACTGGCATTATCGAGTATCCCTTCGATCTCGATAGT
ATTATATTCAGGATGGTAGACGTGGGTGGACAAAGGAGCGAGCGGAGAAAATGGATCCACTGTT
TCGAGAACGTAACGTCCATAATATTCCTGGTCGCTCTTTCCGAGTATGACCAAATACTGTTCGA
ATCCGATAACGAAAACAGAATGGAGGAAAGCAAAGCCCTCTTTAAGACAATTATTACATACCCA
TGGTTCCAGAACAGTTCCGTCATACTCTTCCTTAACAAGAAAGATCTTCTTGAGGAAAAAATCA
TGTACTCTCATTTGGTCGATTATTTCCCCGAATATGACGGCCCTAAGAAAGACGCGATACACGC
CCGCGAATTTATCCTGAAGATGTTTGTCGACCTTAATCCAGATTCTGAAAAGATAATTTACAGC
CACTTCACTTGTGCGACCGATACCGAAAACATACGCTTCGTTTTTGCCGCTGTCAAGGATACGA
TACTGCAGCTTAACTTGAAGGAGTACAATCTGTAA

>XP_029846564.1 guanine nucleotide-binding protein G(q) subunit alpha isoform X1 [Ixodes scapularis]
MMACCISEEAKEQRRINQEIERQLRKDKRDARRELKLLLLGTGESGKSTFIKQMRIIHGHGYNE
EDKKGFIKLVYQNIFMAMQSMIKAMDMLKIQYTNPDNIEHANLVGTVDYETVTTFDGPYVVAIK
KLWADGGIQECYDRRREYQLTDSAKYYLSDVDRIAATDYLPTQQDILRVRVPTTGIIEYPFDLD
SIIFRMVDVGGQRSERRKWIHCFENVTSIIFLVALSEYDQILFESDNENRMEESKALFKTIITY
PWFQNSSVILFLNKKDLLEEKIMYSHLVDYFPEYDGPKKDAIHAREFILKMFVDLNPDSEKIIY
SHFTCATDTENIRFVFAAVKDTILQLNLKEYNLV

**a**

```
NP_001269468_GNB1   1   --------------------------------------------------------MSELDQLRQEAEQLKNQIRDARKACADATLSQITNNIDPVGRIQMRTRRTLRGH
NP_005264_GNB2      1   --------------------------------------------------------MSELEQLRQEAEQLRNQIRDARKACGDSTLTQITAGLDPVGRIQMRTRRTLRGH
AAH00115_GNB3       1   --------------------------------------------------------MGEMEQLRQEAEQLKKQIADARKACADVTLAELVSGLEVVGRVQMRTRRTLRGH
NP_067642_GNB4      1   --------------------------------------------------------MSELEQLRQEAEQLRNQIQDARKACNDATLVQITSNMDSVGRIQMRTRRTLRGH
XP_002411384_isGNB1 1   --------------------------------------------------------MNELDSLRQEAETLKNTIRDARKAACDTTLVQATANMEPVGRIQMRTRRTLRGH
NP_006569_GNB5      1   ------------------------------------------MATEGL------HENETLASLKSEAESLKGKLEEERAKLHDVELHQVAERVEALGQFVMKTRRTLKGH
NP_057278_GNB5L     1   MCDQTFLVNVFGSCDKCFKQRALRPVFKKSQQLSYCSTCAEIMATEGL------HENETLASLKSEAESLKGKLEEERAKLHDVELHQVAERVEALGQFVMKTRRTLKGH
XP_042144999_isGNB5 1   ------------------------------------------MATEGTTMKVDHQAPETVETLTREIEQLKARLEEERKKLNDVALSTVAQRLEAVANLNIKPRRVLKGH

NP_001269468_GNB1   55  LAKIYAMHWGTDSRLLVSASQDGKLIIWDSYTTNKVHAIPLRSSWVMTCAYAPSGNYVACGGLDNICSIYNLKT--REGNVRVSRELAGHTGYLSCCRFLD-DNQIVTSS
NP_005264_GNB2      55  LAKIYAMHWGTDSRLLVSASQDGKLIIWDSYTTNKVHAIPLRSSWVMTCAYAPSGNFVACGGLDNICSIYSLKT--REGNVRVSRELPGHTGYLSCCRFLD-DNQIITSS
AAH00115_GNB3       55  LAKIYAMHWATDSKLLVSASQDGKLIVWDSYTTNKVHAIPLRSSWVMTCAYAPSGNFVACGGLDNMCSIYNLKS--REGNVKVSRELSAHTGYLSCCRFLD-DNNIVTSS
NP_067642_GNB4      55  LAKIYAMHWGYDSRLLVSASQDGKLIIWDSYTTNKMHAIPLRSSWVMTCAYAPSGNYVACGGLDNICSIYNLKT--REGNVRVSRELPGHTGYLSCCRFLD-DSQIVTSS
XP_002411384_isGNB1 55  LAKIYAMHWGSDSRYLVSASQDGKLIVWDAYTTNKVHAIPLRSSWVMTCAYAPTGSYVACGGLDNICSIYSLKT--REGNVRVSRELPGHTGYLSCCRFVD-DNQIVTSS
NP_006569_GNB5      63  GNKVLCMDWCKDKRRIVSSSQDGKVIVWDSFTTNKEHAVTMPCTWVMACAYAPSGCAIACGGLDNKCSVYPLTFDKNENMAAKKKSVAMHTNYLSACSFTNSDMQILTAS
NP_057278_GNB5L     105 GNKVLCMDWCKDKRRIVSSSQDGKVIVWDSFTTNKEHAVTMPCTWVMACAYAPSGCAIACGGLDNKCSVYPLTFDKNENMAAKKKSVAMHTNYLSACSFTNSDMQILTAS
XP_042144999_isGNB5 69  QGKVLCSDWSLDKRHIVSSSQDGKMIIWDAFTTNKEHAVTMPTTWVMACAYAPSGNMVACGGLDNKVTVYPLSF--EEDVSTKKKAVGTHTSYMSCCLFPNSDQQILTGS

NP_001269468_GNB1   162 GDTTCALWDIETGQQTTTFTGHTGDVMSLSLAPD--TRLFVSGACDASAKLWDVREGMCRQTFTGHESDINAICFFPNGNAFATGSDDATCRLFDLRADQELMTYSHDNI
NP_005264_GNB2      162 GDTTCALWDIETGQQTVGFAGHSGDVMSLSLAPD--GRTFVSGACDASIKLWDVRDSMCRQTFIGHESDINAVAFFPNGYAFTTGSDDATCRLFDLRADQELLMYSHDNI
AAH00115_GNB3       162 GDTTCALWDIETGQQKTVFVGHTGDCMSLAVSPD--FNLFISGACDASAKLWDVREGTCRQTFTGHESDINAICFFPNGEAICTGSDDASCRLFDLRADQELICFSHESI
NP_067642_GNB4      162 GDTTCALWDIETAQQTTTFTGHSGDVMSLSLSPD--MRTFVSGACDASSKLWDIRDGMCRQSFTGHVSDINAVSFFPNGYAFATGSDDATCRLFDLRADQELLLYSHDNI
XP_002411384_isGNB1 162 GDMTCALWDIETGQQCTSFTGHTGDVMSLSLSPD--FRTFVSGACDASAKLWDVRDGMCKQTFPGHESDINAVTFFPNGYAFATGSDDATCRLFDMRADQELAMYSHDNI
NP_006569_GNB5      173 GDGTCALWDVESGQLLQSFHGHGADVLCLDLAPSETGNTFVSGGCDKKAMVWDMRSGQCVQAFETHESDINSVRYYPSGDAFASGSDDATCRLYDLRADREVAIYSKESI
NP_057278_GNB5L     215 GDGTCALWDVESGQLLQSFHGHGADVLCLDLAPSETGNTFVSGGCDKKAMVWDMRSGQCVQAFETHESDINSVRYYPSGDAFASGSDDATCRLYDLRADREVAIYSKESI
XP_042144999_isGNB5 177 GDSTCALWDVECSQLLQSFHGHSGDVMALDLSPTEMGNTFVSAGCDRQALVWDMRSGQCVQSFQGHESDINTVKFYPSGDAIATGSDDATCRLYDLRADREVAVYSKQSI

NP_001269468_GNB1   270 ICGITSVSFSKSGRLLLAGYDDFNCNVWDALKADRAGVLAGHDNRVSCLGVTDDGMAVATGSWDSFLKIWN
NP_005264_GNB2      270 ICGITSVAFSRSGRLLLAGYDDFNCNIWDAMKGDRAGVLAGHDNRVSCLGVTDDGMAVATGSWDSFLKIWN
AAH00115_GNB3       270 ICSITSVAFSLSGRLLFAGYDDFNCNVWDSMKSERVGILSGHDNRVSCLGVTADGMAVATGSWDSFLKIWN
NP_067642_GNB4      270 ICGITSVAFSKSGRLLLAGYDDFNCNVWDTLKGDRAGVLAGHDNRVSCLGVTDDGMAVATGSWDSFLRIWN
XP_002411384_isGNB1 270 ICGITSVAFSKSGRLLLAGYDDFNCNVWDSMKAERAGVLAGHDNRVSCLGVTEDGMAVATGSWDSFLKIWN
NP_006569_GNB5      283 IFGASSVDFSLSGRLLFAGYNDYTINVWDVLKGSRVSILFGHENRVSTLRVSPDGTAFCSGSWDHTLRVWA
NP_057278_GNB5L     325 IFGASSVDFSLSGRLLFAGYNDYTINVWDVLKGSRVSILFGHENRVSTLRVSPDGTAFCSGSWDHTLRVWA
XP_042144999_isGNB5 287 IFGVNSVDFSVSGRLLFAGYNDYTVNVWDALKCVRLSILYGHENRVTCLKVSPDGTALSTGSWDFTLRVWA
```

**b**

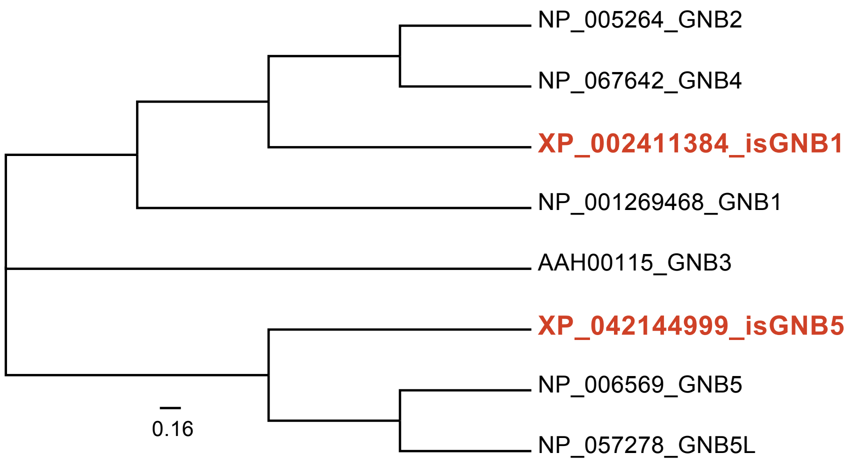

#### Percent Identity Matrix for *Ixodes* and human GNB amino acid sequences

| Species | Gene | Accession # | AAH00115 | P_002411384 | NP_005264 | NP_00126946 | KAI2532546 | P_04214499 | NP_006569 | NP_057278 |
|---|---|---|---|---|---|---|---|---|---|---|
| H. sapiens | GNB3 | AAH00115 | 100 | 79.41 | 80.59 | 82.94 | 79.41 | 51.62 | 51.18 | 51.18 |
| I. scapularis | GNB1 | XP_002411384 | 79.41 | 100 | 86.76 | 87.94 | 86.47 | 52.21 | 52.94 | 52.94 |
| H. sapiens | GNB2 | NP_005264 | 80.59 | 86.76 | 100 | 90.29 | 90 | 52.21 | 51.76 | 51.76 |
| H. sapiens | GNB1 | NP_001269468 | 82.94 | 87.94 | 90.29 | 100 | 90.88 | 52.51 | 52.65 | 52.65 |
| H. sapiens | GNB4 | KAI2532546 | 79.41 | 86.47 | 90 | 90.88 | 100 | 53.39 | 52.06 | 52.06 |
| I. scapularis | GNB5 | XP_042144999 | 51.62 | 52.21 | 52.21 | 52.51 | 53.39 | 100 | 71.51 | 71.51 |
| H. sapiens | GNB5 | NP_006569 | 51.18 | 52.94 | 51.76 | 52.65 | 52.06 | 71.51 | 100 | na |
| H. sapiens | GNB5L | NP_057278 | 51.18 | 52.94 | 51.76 | 52.65 | 52.06 | 71.51 | na | 100 |

**a** **GNAQ**

```
H. sapiens      1 MTLESIMACCLSEEAKEARRINDEIERQLRRDKRDARRELKLLLLGTGESGKSTFIKQMRIIHGSGYSDEDKRGFTKLVYQNIFTAMQAM
M. muscuculus   1 MTLESIMACCLSEEAKEARRINDEIERQLRRDKRDARRELKLLLLGTGESGKSTFIKQMRIIHGSGYSDEDKRGFTKLVYQNIFTAMQAM
I. scapularis   1 -----MMACCISEEAKEQRRINQEIERQLRKDKRDARRELKLLLLGTGESGKSTFIKQMRIIHGHGYNEEDKKGFIKLVYQNIFMAMQSM
XP_029846564

                 91 IRAMDTLKIPYKYEHNKAHAQLVREVDVEKVSAFENPYVDAIKSLWNDPGIQECYDRRREYQLSDSTKYYLNDLDRVADPAYLPTQQDVL
                 91 IRAMDTLKIPYKYEHNKAHAQLVREVDVEKVSAFENPYVDAIKSLWNDPGIQECYDRRREYQLSDSTKYYLNDLDRVADPSYLPTQQDVL
                 86 IKAMDMLKIQYTNPDNIEHANLVGTVDYETVTTFDGPYVVAIKKLWADGGIQECYDRRREYQLTDSAKYYLSDVDRIAATDYLPTQQDIL

                181 RVRVPTTGIIEYPFDLQSVIFRMVDVGGQRSERRKWIHCFENVTSIMFLVALSEYDQVLVESDNENRMEESKALFRTIITYPWFQNSSVI
                181 RVRVPTTGIIEYPFDLQSVIFRMVDVGGQRSERRKWIHCFENVTSIMFLVALSEYDQVLVESDNENRMEESKALFRTIITYPWFQNSSVI
                176 RVRVPTTGIIEYPFDLDSIIFRMVDVGGQRSERRKWIHCFENVTSIIFLVALSEYDQILFESDNENRMEESKALFKTIITYPWFQNSSVI

                271 LFLNKKDLLEEKIMYSHLVDYFPEYDGPQRDAQAAREFILKMFVDLNPDSDKIIYSHFTCATDTENIRFVFAAVKDTILQLNLKEYNLV
                271 LFLNKKDLLEEKIMYSHLVDYFPEYDGPQRDAQAAREFILKMFVDLNPDSDKIIYSHFTCATDTENIRFVFAAVKDTILQLNLKEYNLV
                266 LFLNKKDLLEEKIMYSHLVDYFPEYDGPKKDAIHAREFILKMFVDLNPDSEKIIYSHFTCATDTENIRFVFAAVKDTILQLNLKEYNLV
```

**b** **GNB1**

```
H. sapiens      1   MSELDQLRQEAEQLKNQIRDARKACADATLSQITNNIDPVGRIQMRTRRTLRGHLAKIYAMHWGTDSRLLVSASQDGKLIIWDSYTTNKV
I. scapularis   1   MNELDSLRQEAETLKNTIRDARKAACDTTLVQATANMEPVGRIQMRTRRTLRGHLAKIYAMHWGSDSRYLVSASQDGKLIVWDAYTTNKV
XP_002411384

                91  HAIPLRSSWVMTCAYAPSGNYVACGGLDNICSIYNLKTREGNVRVSRELAGHTGYLSCCRFLDDNQIVTSSGDTTCALWDIETGQQTTTF
                91  HAIPLRSSWVMTCAYAPTGSYVACGGLDNICSIYSLKTREGNVRVSRELPGHTGYLSCCRFVDDNQIVTSSGDMTCALWDIETGQQCTSF

               181  TGHTGDVMSLSLAPDTRLFVSGACDASAKLWDVREGMCRQTFTGHESDINAICFFPNGNAFATGSDDATCRLFDLRADQELMTYSHDNII
               181  TGHTGDVMSLSLSPDFRTFVSGACDASAKLWDVRDGMCKQTFPGHESDINAVTFFPNGYAFATGSDDATCRLFDMRADQELAMYSHDNII

               271  CGITSVSFSKSGRLLLAGYDDFNCNVWDALKADRAGVLAGHDNRVSCLGVTDDGMAVATGSWDSFLKIWN
               271  CGITSVAFSKSGRLLLAGYDDFNCNVWDSMKAERAGVLAGHDNRVSCLGVTEDGMAVATGSWDSFLKIWN
```

**c** **KisspeptinR + Gq (8xgs)**

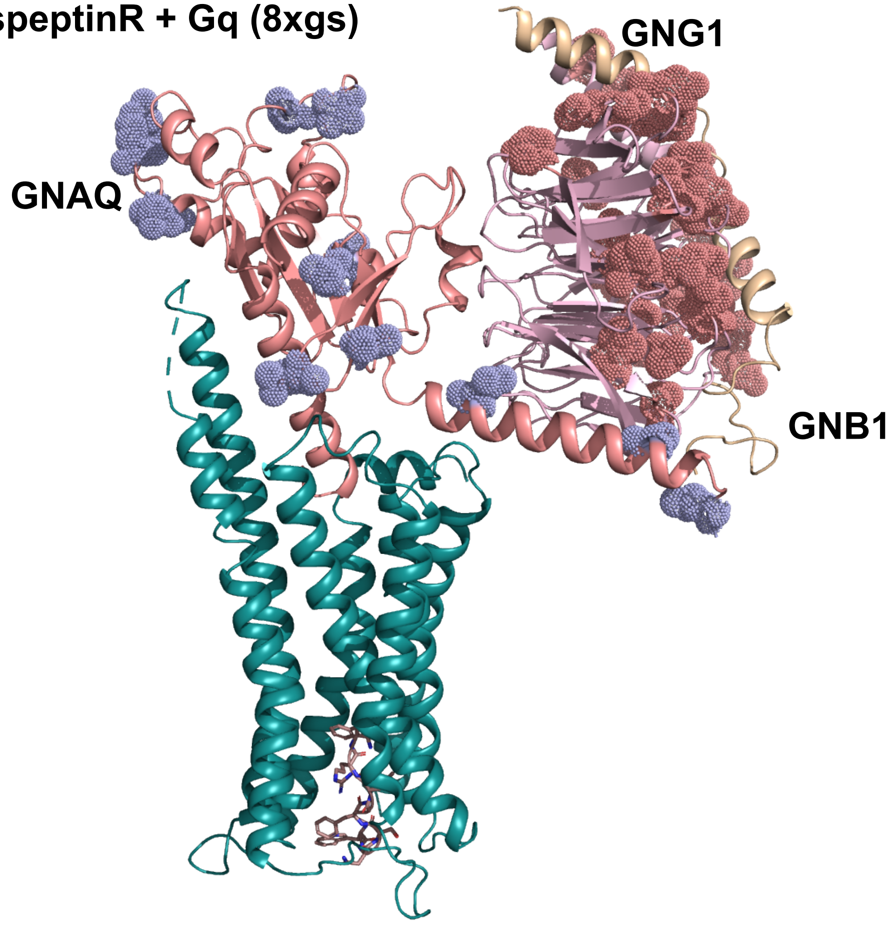

**a** **PLCβ**

```
PLCB3_HUMAN  1 MAGAQPGVHALQLEPPTVVETLRRGSKFIKWDEETSSRNLVTLRVDPNGFFLYWTGPNMEVDTLDISSIRDTRTGRYARLPKDPKIREVLGFGGPDARLEEKLMTVVSGP
XP_029835062 1 MAGAKSGVHVLQLKLISVPQSLQEGNKFVKWDDDSALGTPVTLRVDKNGFFLFWTDQNKETEFLDISSIRDTRTGKYARTPKEGKLRDSVSMGASDTPLEEKTLTVVYGP

             111 DPVNTVFLNFMAVQDDTAKVWSEELFKLAMNILAQNASRNTFLRKAYTKLKLQVNQDGRIPVKNILKMFSA---DKKRVETALESCGLKFNRSESIRPDEFSLEIFERFL
             111 DLVNVSVINFCCNSRETAQLWTDELLRMAYNLLSLNAPASRFLEKAHTKLLLTTDREGRLPVKGVLRMFAQHRDDRRRVERALDAVGLAAGKNDTLAPDRLCFDVFLRFY

             218 NKLCLRPDIDKILLEI--GAKGKPYLTLEQLMDFINQKQRDPRLNEVLYPPLRPSQARLLIEKYEPNQQFLERDQMSMEGFSRYLGGEENGILPLEALDLSTDMTQPLSA
             221 RQLVGRQEVDAIFERLCGGAK-KKAMTVDQLVDFLNKEQRDPRLNEILYPYANPARAKDIIAQYEPNKSYVAKGLFSVEGFLRYLMSADNPIVSPEKFDLSLDMDQPLNH

             326 YFINSSHNTYLTAGQLAGTSSVEMYRQALLWGCRCVELDVWKGRPPEEEPFITHGFTMTTEVPLRDVLEAIAETAFKTSPYPVILSFENHVDSAKQQAKMAEYCRSIFGD
             330 YFINSSHNTYLTGHQLTGKSSVEMYRQCLLAGCRCIELDCWTGRNSDEEPIITHGYTVVTEVLLREVIEAIAESAFKTSDFPVVLSFENHC-SPKQQAKMANYCRKLFGD

             436 ALLIEPLDKYPLAPGVPLPSPQDLMGRILVKNKKRHRPSAGGPDSAGRKRPL-EQSNSALSESSAATEPSSPQLGSPSSDSCPGLSNGEEVGLEKPSLEPQKSLGDEGLN
             439 MLVTEPMLSHPLRPGQPLPSPQQLLRKIIIKNKKKHH--------ARPHKPLASPSVAAAAAAAAVAPG----GGGEDDASPG------------------EGPDDLN

             545 RGPYVLGPADREDEEEDEEEEEQTDPKKPTTDEGTASSEVNATEEMSTLVNYIEPVKFKSFEAARKRNKCFEMSSFVETKAMEQLTKSPMEFVEYNKQQLSRIYPKGTRV
             517 -GDAKGSEADDWDTDSGTEEEEAPERSAEDQNEGTAAKESEAVAEMSALVNYIQPVRFHSFEHAEKRDRSYEISSFVETQATNLLKEHPVEFVNYNKRQLSRIYPSGTRV

             655 DSSNYMPQLFWNVGCQLVALNFQTLDVAMQLNAGVFEYNGRSGYLLKPEFMRRPDKSFDPFTEVIVDGIVANALRVKVISGQFLSDRKVGIYVEVDMFGLPVDT-RRKYR
             627 SSSNYMPQVFWNAGCQLIALNFQTLDLGMQLNLGIFEFNGRSGYLLKPDFMRRADRKFDPFTESTVDGIIAGTVSIRIISGQFLTDKHVGTYVEVDMYGLPADTVRRRFR

             764 TRTSQGNSFNPVWDEEPFDFPKVVLPTLASLRIAAFEEGGKFVGHRILPVSAIRSGYHYVCLRNEANQPLCLPALLIYTEASDYIPDDHQDYAEALINPIKHVSLMDQRA
             737 TRTVPANGINPVYDEEPFIFKKVVLPDLAVLRISVCDDSGKLLGHRILPVVGLRPGYRHISLRNESGQPLLLQTLFVHVTVKDYVPDGLSELADALANPIKYQSRIEKHA

             874 RQLAALIGESEAQAGQETCQDTQSQ-----QLGSQPSSNPTPSP--------------LDAS---------------------PRRPPGPTT---------SPASTSLSS
             847 TQLRALTDDLDEGLAAESRSPVQTQAPATSTLGACPQGASSPSPGASSGDARPSAACGIDAADTAAPSATLPVAAALNNGGLQPRSPAPPLARQDTLTRKMDPTSRSLSD

             935 PGQRD---DLIASILSEVAPTPLDELRGHKALVKLRSRQERDLRELRKKHQR---KAVTLTRRLLDGLAQAQAEGRCRL-----RPGALGGAADVEDTK----------E
             957 ECTQEKTLSLLESPEAQLSAEPLGKLREHKTVQKVLSKLDKDLSVVRKRFDKLRDKEREMQTQRADKLSQANEKHRAQLSKSHSKLGKKFSCGDVMALKQQNESQIQVLD

            1024 GEDEAKRYQEFQNRQVQSLLELREAQVDAEAQRRLEHLRQALQRLREVVLDANTTQFKRLKEMNEREKKELQKILDRKRHNSISEAKMRDKHKKEAEL----TEINRRHI
            1067 AEHRAKQ-EELQQSHNLAMMNITKEQYKAEMDIQHKYLDSLFNAMEKTMQGSQAAQMQQLQDLHDKEVSELMKRLEAQTKEEM--RSLNKKHKDKNELDRIKRELHQKMI

            1130 TESVNSIRRLEEAQKQRHDRLVAGQQQVLQQLAEEEPKLLAQLAQECQEQRARLPQEIRRSLLGEMPEGLGDGPLVACASNGH
            1174 GEAVSERQRISGLLDKKKAELEKQHEDVKKSLEEEKSQAIVKQQQE---------YELKCS---QLSSSLSDNPALFAESSDH
```

**b**

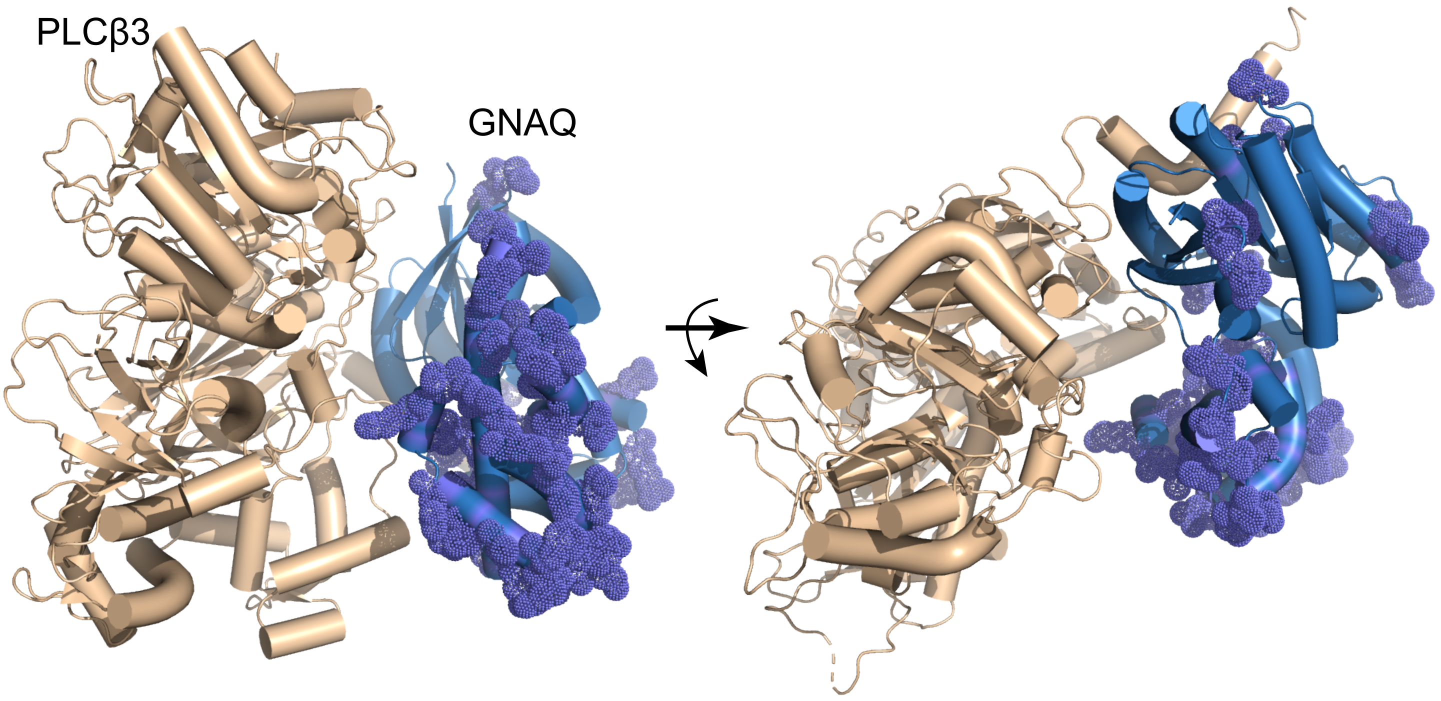

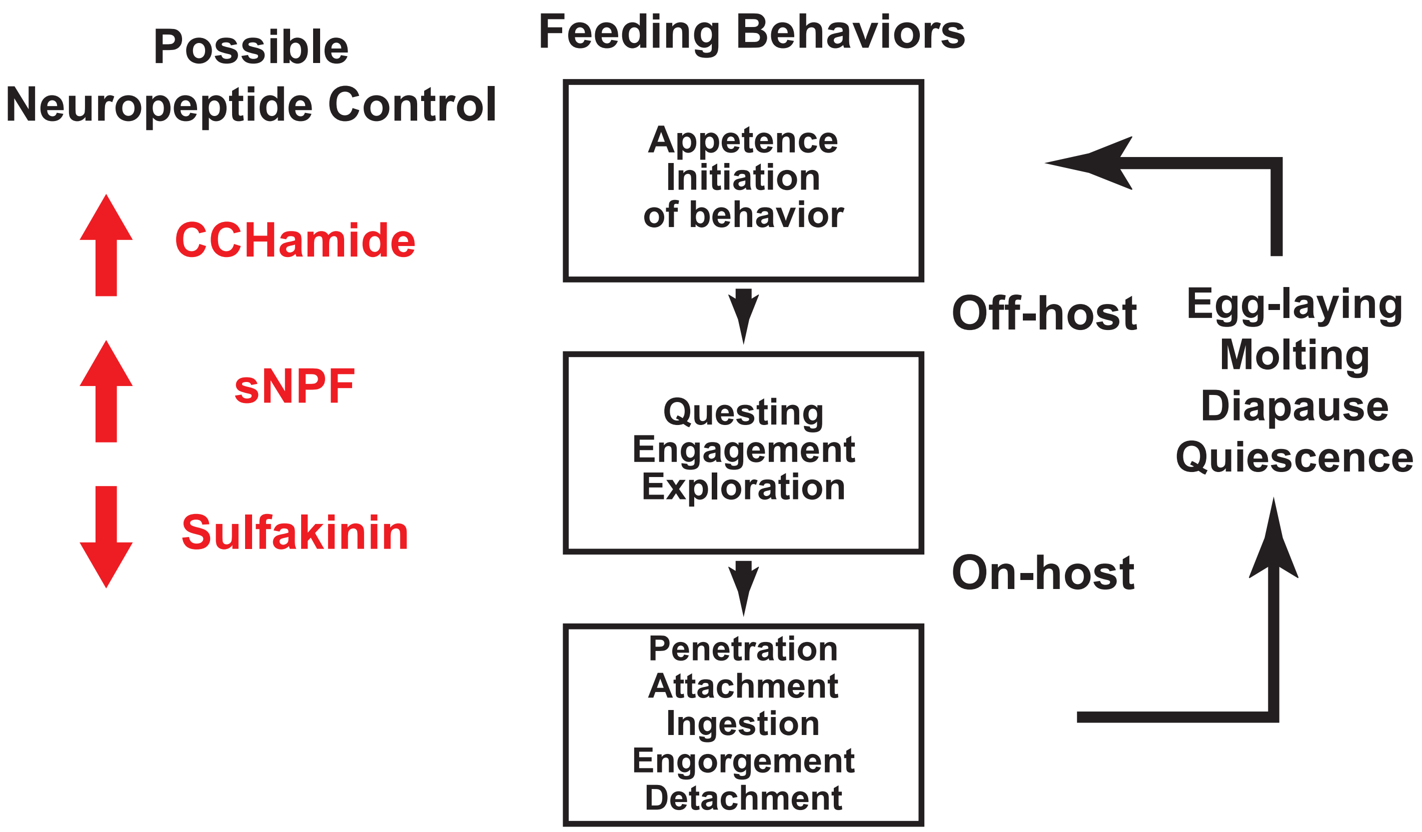
Possible
Neuropeptide Control
CCHamide
sNPF
Sulfakinin
Feeding Behaviors
Appetence
Initiation
of behavior
Questing
Engagement
Exploration
Penetration
Attachment
Ingestion
Engorgement
Detachment
Off-host
On-host
Egg-laying
Molting
Diapause
Quiescence
